# Fine-Resolution Asymmetric Migration Estimation

**DOI:** 10.1101/2025.05.29.656894

**Authors:** Hao Shen, John Novembre

## Abstract

The genetic structure of populations is often shaped by processes and events that introduce asymmetries to gene flow between geographic locations. Here, we first develop an algorithm that allows efficient computation of pairwise coalescent times in time-homogeneous models of population structure at migration-drift equilibrium. We then use the algorithm as the foundation for a new method to infer asymmetric migration rates in spatial models of population structure. The inferred equilibrium migration rates provide a novel representation of the geographic structure of genetic variation. We assess the method using a variety of simulated histories of gene flow, and apply the method to datasets from poplar trees, North American gray wolves, and human archaeogenetic samples, revealing complex asymmetric migration signals and providing a more refined view of the geographic structure of genetic variation.

## Introduction

In spatially structured populations, migration over generations shapes genetic variation and often leads to isolation-by-distance—a phenomenon in which geographically proximal individuals are more genetically similar than those farther apart (Wright, 1943, 1946). Although widespread across species and ecosystems (Slatkin, 1985; Meirmans, 2012), this phenomenon can be complex due to spatial heterogeneity in gene flow, which causes the relationship between genetic distance and geographic distance to vary across the landscape (Bradburd and Ralph, 2019).

Many methods have been developed to reveal spatial heterogeneity in isolation-by-distance (House and Hahn, 2017; Bradburd et al., 2016; Petkova et al., 2016; Marcus et al., 2021). Of particular interest here are the methods EEMS (Petkova et al., 2016) and FEEMS (Marcus et al., 2021), which model a population in terms of a large network of connected demes at migration-drift equilibrium and infer symmetric migration rates between demes. These methods produce visual maps of regions of high/low effective migration rates, offering insights into spatial variation in gene flow.

In modeling spatial heterogeneity in isolation-by-distance, as in other areas of population genetics, a coalescent-based perspective is incredibly helpful. In particular, expected pairwise coalescent times can be used to calculate the expected values of basic summaries such as *F*_*ST*_ (Slatkin, 1991), as well as in fitting models of population structure, as in EEMS and related approaches (Petkova et al., 2016; Lundgren and Ralph, 2019). For a metapopulation composed of several subpopulations or ‘demes’, the structured coalescent (Notohara, 1990; Takahata, 1991) provides a natural framework for the computation of expected pairwise coalescence times based on backward migration and coalescence rates.

However, fast computation of the expected pairwise coalescent times in the structured coalescent is a major challenge. For symmetric migration, a two-pronged strategy has been employed to reduce the time complexity. The first part of the strategy is to decompose the pairwise coalescence time into the time for two lineages to enter the same deme and the time for two lineages within the same deme to coalesce (Slatkin, 1991). The second part is to approximate the expected time for two lineages to enter the same deme as one-fourth of the expected commute time of the Markov chain defined by single-lineage migration (Matsen and Wakeley, 2006). A well-established connection between circuit theory and random walks (Nash-Williams, 1959) equates expected commute times with effective resistances (i.e., resistance distances) in the corresponding electrical network, for which efficient algorithms are available (Babić et al., 2002; Bapat, 2004). This two-pronged strategy also forms the basis of the widely used ‘isolation-by-resistance’ framework (McRae, 2006; Hanks and Hooten, 2013; Dickson et al., 2018).

In the case of isotropic symmetric migration, this strategy yields exact pairwise coalescence times. Under anisotropic but still pairwise symmetric migration, it produces approximate results and underpins methods such as EEMS (Petkova et al., 2016) and FEEMS (Marcus et al., 2021).

However, many natural systems exhibit pronounced asymmetries in gene flow, which these symmetric models fail to represent. Asymmetric migration can arise from persistent directional forces such as ocean currents (White et al., 2010; Bertola et al., 2020), prevailing winds (Kling and Ackerly, 2021), or river networks (Alp et al., 2012; Thomaz et al., 2016), as well as from historical demographic events such as range expansions (Ramachandran et al., 2005; Excoffier et al., 2009) and mass migrations (Haak et al., 2015; Lipson et al., 2017). Accurately modeling such asymmetries is crucial for reconstructing spatial population histories.

From a computational standpoint, accounting for asymmetry introduces significant challenges. As pointed out by Lundgren and Ralph (2019), approximate strategies that perform well under symmetric assumptions may lead to substantial distortions when applied to asymmetric settings. To address this issue, Lundgren and Ralph proposed directly solving the matrix equation for expected pairwise coalescent times by vectorizing it into a system of linear equations. While this approach provides exact solutions, its computational complexity scales as *O*(*d*^6^), where *d* is the number of demes, making it impractical for large-scale applications.

Here, we introduce FRAME (Fine-Resolution Asymmetric Migration Estimation). FRAME employs techniques from computational linear algebra and solves the structured coalescent equation with time complexity of *O*(*d*^4^). By integrating this approach with the efficient gradient-based optimization and penalized-likelihood in FEEMS, FRAME enables the inference of fine-resolution asymmetric migration patterns with significantly improved computational efficiency.

## Results

### Model specification and inference strategy

We model population structure using a weighted directed graph, in which vertices represent demes and directed edges represent migration between demes. A directed edge from deme *i* to deme *j* is assigned a weight *m*_*ij*_, which represents the rate at which genetic lineages in deme *i* are drawn from deme *j*. These migration rates form a migration rate matrix *M*, which is the weighted adjacency matrix of the graph and encapsulates all the parameters governing gene exchange between demes. The graph Laplacian is then given by *L* = *D* − *M*, where *D* is a diagonal matrix with entries 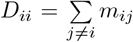. By construction, each row of *L* sums to zero, and the backward-in-time migration dynamics of the structured coalescent correspond to a continuous-time jump process on the graph, with −*L* serving as the transition-rate (generator) matrix.

We parameterize the coalescence rate per deme by *γ* = (*γ*_1_, *γ*_2_, …, *γ*_*d*_), where *γ*_*i*_ represents the coalescencerate in deme *i*. The equation for expected pairwise coalescence times at migration-drift equilibrium can then be computed from *L* and *γ* (Hill, 1972; *Strobeck, 1987; Whitlock and Barton, 1997; Tufto et al., 1998)*

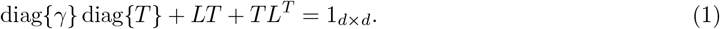

Here *T* is the matrix such that its (*i, j*)_th_ entry *T*_*ij*_ represents the expected coalescence time between two randomly chosen haploid samples, one from deme *i* and another from deme *j*, 1_*d×d*_ is a matrix of all ones.

Solving equation 1 using direct vectorized methods is computationally expensive: solving a linear system with *O*(*d*^2^) unknowns has a complexity of *O*(*d*^6^). This computational burden limits the scalability of existing structured coalescent–based methods when applied to large datasets (Lundgren and Ralph, 2019).

To overcome this limitation, we develop a method that solves it with a complexity of *O*(*d*^4^), enabling larger-scale inference in terms of the number of demes (Methods and Supplementary Note). To do so, we use techniques from computational linear algebra and exploit the structural similarity between this equation and the Lyapunov equation. The new method yields substantial improvements in computation time (see Fig. S1, mean run-time for 100 demes 0.13 seconds for the novel approach vs 3.82 seconds for the direct vectorized method).

Building upon this computational advance, we develop the inference machinery of FRAME. Detailed computational steps are provided in the Methods section and Supplementary Note. Here, we briefly outline the logic and key steps.

An important aspect of the parameterization is the relationship between *γ* and *M* . If population sizes are fully determined by migration, the jump process specified by *M* implies a corresponding stationary probability distribution vector *π* = (*π*_1_, *π*_2_, …, *π*_*d*_). In such a case, if we denote the total population size by *N*_*T*_, the population size of deme *i* can be expressed as *N*_*T*_ *π*_*i*_, and *γ*_*i*_ would be 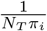 (Strobeck, 1987). However, such a model can be unrealistic in many cases, for example, when population sizes are determined by factors other than the movement of gametes/individuals (e.g., local carrying capacity).

One way to relax the relationship between coalescence rates and the stationary distribution is to set the coalescence rates as free parameters. However, this approach increases the risk of identifiability issues. An alternative is to assume all coalescence rates are equal, yet this constrains the model unrealistically. To address this, we compromise by setting 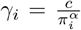, where *c* is a scaling constant greater than 0, and *α* is a constant between 0 and 1. When *α* is 0, the model assumes that all coalescence rates are equal. When *α* is 1, the model corresponds to the case where the coalescence rates *γ* are fully determined by the stationary distribution *π* implied by *M* (in which case *c* is interpretable as 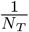).

Similar to FEEMS, FRAME uses a penalized maximum likelihood approach to estimate *M, c*, and *α*. That is, we minimize the objective function *l*(*M, c, α*) + *λ*_*m*_Ψ(*M*) where *l*(*M, c, α*) = − log *P* (*D* | *M, c, α*) denotes the negative log-likelihood of the parameters given the observed genetic distances, Ψ(*M*) represents a penalty function that favors smoothness in the inferred weight parameters, and *λ*_*m*_ is a scalar multiplier of the penalty function. For the penalty function, FRAME employs a formulation that slightly differs from that of FEEMS (Methods and Supplementary Note). For fixed values of *λ*_*m*_, we solve the optimization problem using the L-BFGS (Byrd et al., 1995) algorithm, and then we use cross-validation based on the model’s prediction of allele frequencies to choose *λ*_*m*_. Finally, as in EEMS and FEEMS, migration rates inferred by FRAME are relative rather than absolute, since the total branch length of the coalescent tree is absorbed into the model. Full computational details are provided in the Methods section and Supplementary Note.

### Example outputs

To provide an example of the outputs, Fig. 1a-d illustrates an application of FRAME using a dataset of two closely related poplar species (*Populus trichocarpa* and *Populus balsamifera*) (Geraldes et al., 2014; Lundgren and Ralph, 2019) (see more detailed discussion below). The input data consists of the latitudes, longitudes, and genotypes of sampled individuals. FRAME constructs a network of demes (detailed in Methods) to model spatial genetic structure and assigns samples to nearby demes based on proximity.

**Figure 1:**
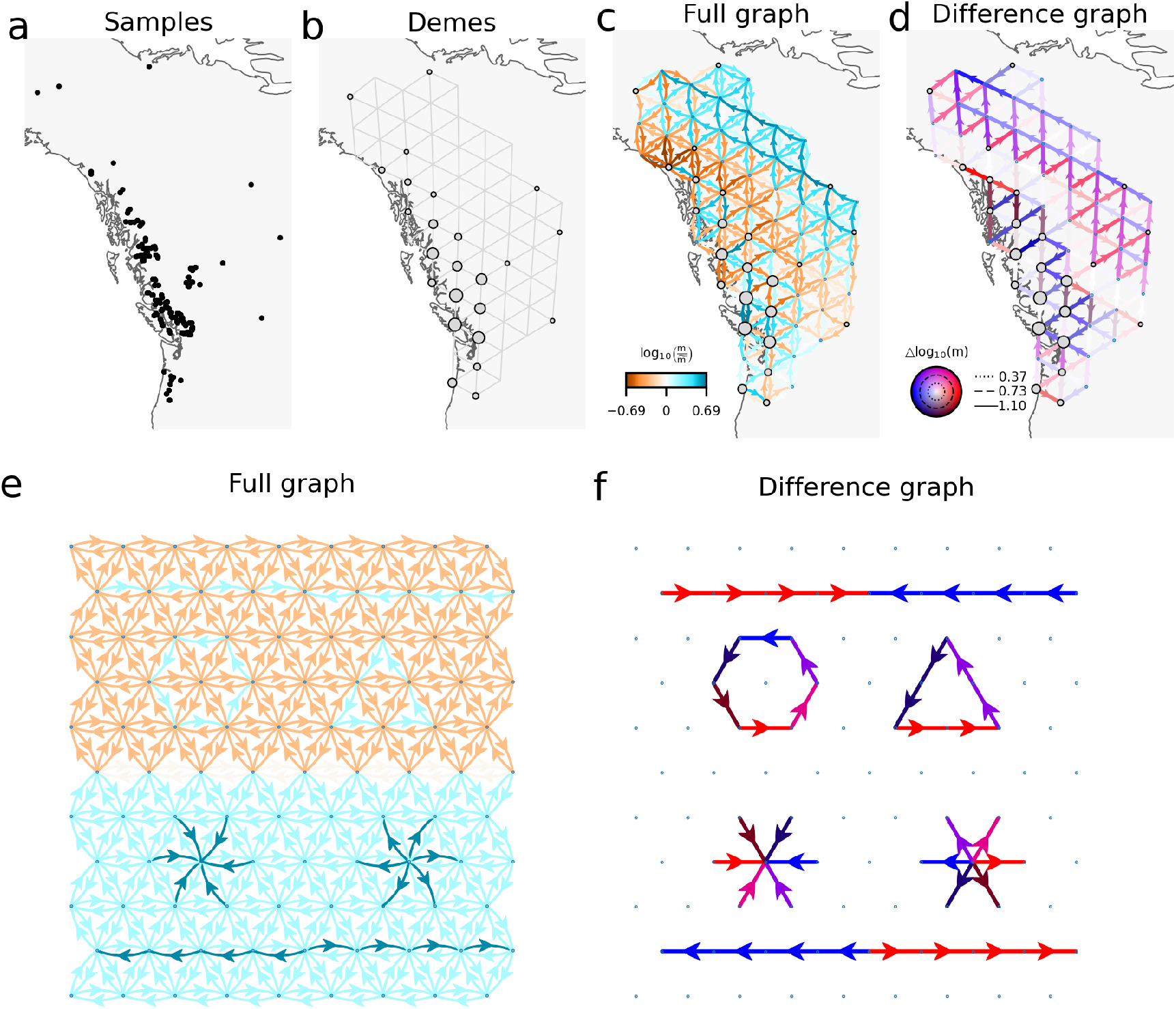
Introduction to FRAME with example outputs. **(a-d)** A schematic overview of FRAME using an example dataset from *Populus trichocarpa* and *Populus balsamifera* (Geraldes et al., 2014; Lundgren and Ralph, 2019). **(a)** The input consists of sample genotypes and their spatial positions. **(b)** FRAME constructs demes and assigns samples to their nearest demes. **(c)** The full graph representation of FRAME fit, displaying all edges. **(d)** The difference graph representation of FRAME fit, emphasizing asymmetries. **(e-f)** Various migration patterns represented by FRAME, including spatially converging lineages, spatially diverging lineages, cyclic rotating lineages, and directionally migrating lineages.

The inferred ‘full graph’ (Fig. 1c) provides an overview of all inferred migration rates across the landscape, capturing both the magnitude and direction of gene flow. In contrast, the ‘difference graph’ output (Fig. 1d) highlights migration asymmetries by calculating the difference in log-scaled migration rates between connected demes, revealing more clearly directional biases in gene flow backward in time (Additional possible summary graphs and related demographic parameters for the empirical analyses that follow are presented in the Supplementary Notes, Fig. S13-S19). It is important to note that the difference graph is shown on a log-scale: for example, migration rate differences between 0.01 and 0.1 as well as between 10^−5^ and 10^−6^, both yield Δ log *m* = 1. While this representation can be convenient in some scenarios, it may be less informative in others. We therefore encourage users to look jointly at the various outputs to aid their interpretation.

Since the graph edges represent backward migration of lineages, the patterns can be naturally interpreted in the context of the structured coalescent. Several possible patterns are shown in Fig. 1e-f. One pattern is where arrows point from surrounding demes to a central deme. This indicates that lineages from surrounding demes migrate backward to the center at higher rates than they migrate away from it. We call this a pattern of ‘spatially converging lineages’. Conversely, a pattern of ‘spatially diverging lineages’ is a pattern where lineages in the central deme migrate backward to the surrounding demes at a higher rate than they migrate to it, suggesting hybridization/admixture-like dynamics are likely to have occurred at the center. Patterns where lineages migrate backward along a path in a certain direction at higher rates represent what we term ‘directionally migrating lineages’. Cycles on the graph are also theoretically possible and for shorthand we call these patterns of ‘cyclic rotating lineages’.

### Results on simulated data from equilibrium stepping-stone models

The performance in solving equation 1 underlies the speed of the overall FRAME method, and allows us to apply the method rapidly across a range of different hypothetical scenarios. Fig. 2 illustrates FRAME’s performance in several distinct time homogeneous stepping-stone migration scenarios at migration-drift equilibrium. The first column shows the results for the combination of small-scale patterns introduced earlier (Fig. 1e,f). The remaining columns correspond to large-scale migration structures, including directionally migrating lineages (second column), spatially converging lineages (third column), and spatially diverging lineages (fourth column). Each example output is based on input consisting of 10 samples per deme and 10^5^ biallelic loci. The corresponding difference graphs and the summary of goodness of fit are shown in Figs. S2-S3.

**Figure 2:**
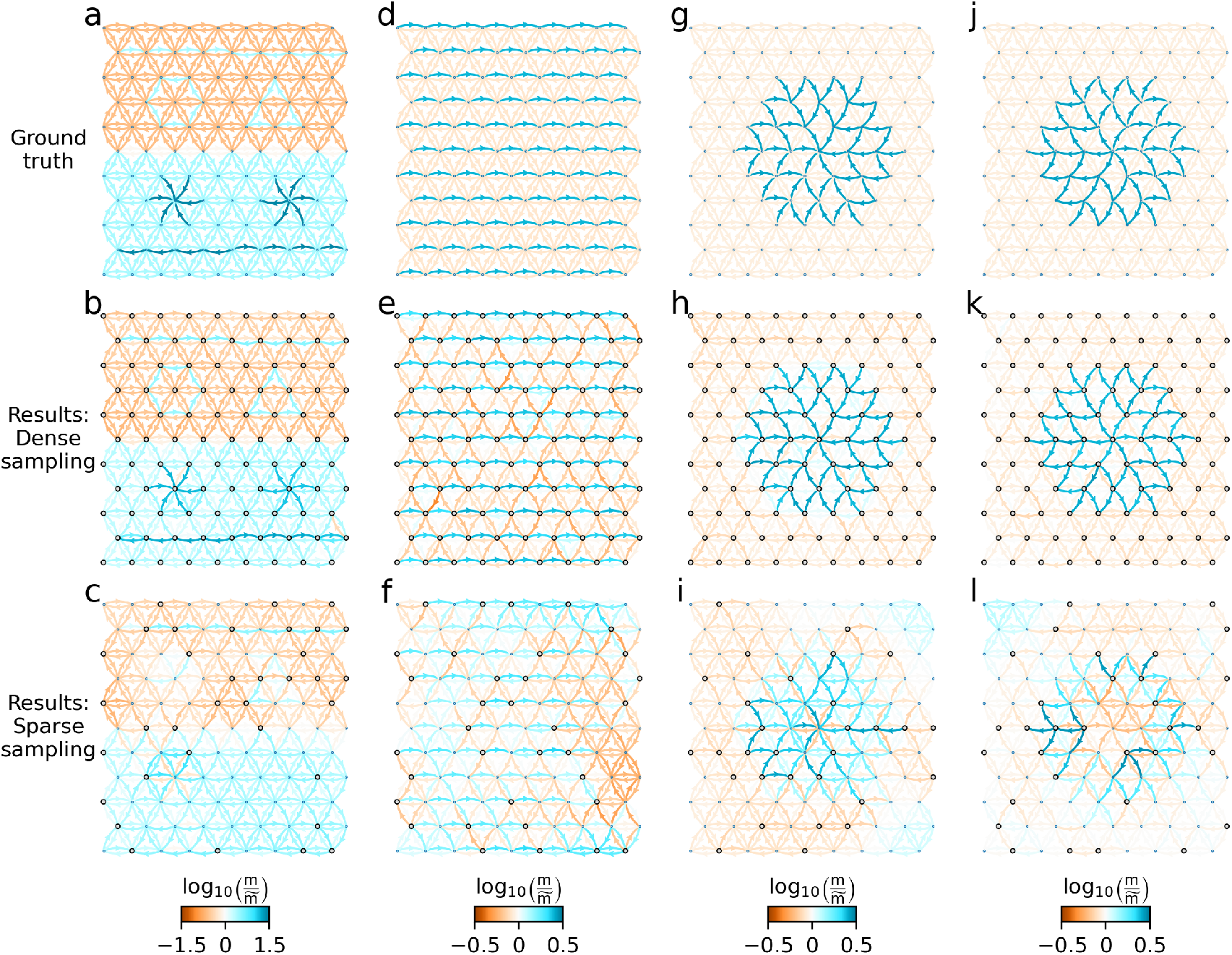
FRAME fit to equilibrium migration simulations with different patterns and sampling schemes. The first column (**a–c**) presents the ground truth and FRAME fit to coalescent simulations featuring small-scale migration patterns. The second column (**d–f**) shows results for simulations with a large scale pattern of directionally migrating lineages (‘west’ to ‘east’ here). The third column (**g–i**) illustrates cases with a large scale patterns of spatially converging lineages, while the final column (**j–l**) represents scenarios with a large scale pattern of spatially diverging lineages (For details of the simulations, see Methods).

Across all large-scale simulations, the ground-truth migration patterns (Fig. 2d–l) consist of a homogeneous baseline backward-in-time migration rate with spatially localized regions of elevated migration. In contrast, the small-scale pattern simulations (Fig. 2a) exhibit spatial heterogeneity in baseline migration rates with embedded patterns defined relative to their local background. In principle, the imposed patterns are distinguishable from surrounding migration while preserving geographic variation in the baseline structure (see Methods for detailed parameter settings).

Simulations are generated using `msprime` (Baumdicker et al., 2022) under a structured coalescent framework, proceeding backward in time. We evaluate FRAME’s performance under both dense sampling, in which all demes are sampled, and sparse sampling, in which only 25% of demes are sampled, to assess how sampling density influences the inferred migration patterns.

When sampling is dense and the dataset is large (Fig. 2b-k), both small-scale patterns and large-scale patterns are faithfully recovered, closely matching the ground truth. In contrast, when sampling is sparse (Fig. 2c-l), with fewer demes containing observed samples, FRAME’s ability to detect migration patterns depends on the scale of the patterns. In our simulations, large-scale patterns remain qualitatively detectable even with sparse sampling, despite a substantial reduction in goodness of fit (Fig. S2). However, small scale patterns are often missed or underrepresented due to the lack of samples near regions where these patterns occur. Under sparse sampling, the inferred networks recover broader migration trends, while finer details are smoothed out or lost. Fig. 2g-i and Fig. 2j-l are interesting to contrast in that the broad structure of the asymmetries is similar in all but direction. The inference results show more fidelity at recovering the true migration parameters for the case of backwards-in-time spatially converging lineages (Fig. 2g-i) vs for the diverging case (Fig. 2j-l).

All simulations in Fig. 2 assume equal population sizes across demes. In Fig. S4-S5 we demonstrate that FRAME can also effectively recover migration patterns when population sizes are proportional to the stationary distribution implied by the migration matrix.

### Results on simulated data from non-equilibrium stepping-stone models

To further evaluate the method’s performance, we extend the backward-in-time simulations to non-equilibrium conditions, as illustrated in Fig. 3 (see Supplementary Note for detailed parameter settings). As a baseline, the first row of Fig. 3 illustrates an equilibrium stepping-stone model with directionally migrating lineages from the left and right boundary to the center. The second row illustrates a range expansion scenario. Forward in time, this corresponds to a classical range expansion, where migrants originating from the central line progressively occupy new demes and undergo exponential growth until reaching carrying capacity. Migration in this scenario is strictly one-way, with no movement occurring from newly colonized demes back to the original central demes. Backward in time, lineages therefore converge toward the central line, as shown in Fig. 3b). The third row represents a scenario of a sequential pulse migration. Forward in time, this corresponds to a wave of asymmetric migration propagating outward from the central line toward both edges, eventually reaching the boundary, added upon a background of steady-state symmetric migration rate between neighboring demes. Backward in time, lineages again converge toward the central line, now in the presence of an additional uniform migration background, as illustrated in Fig. 3c.

**Figure 3:**
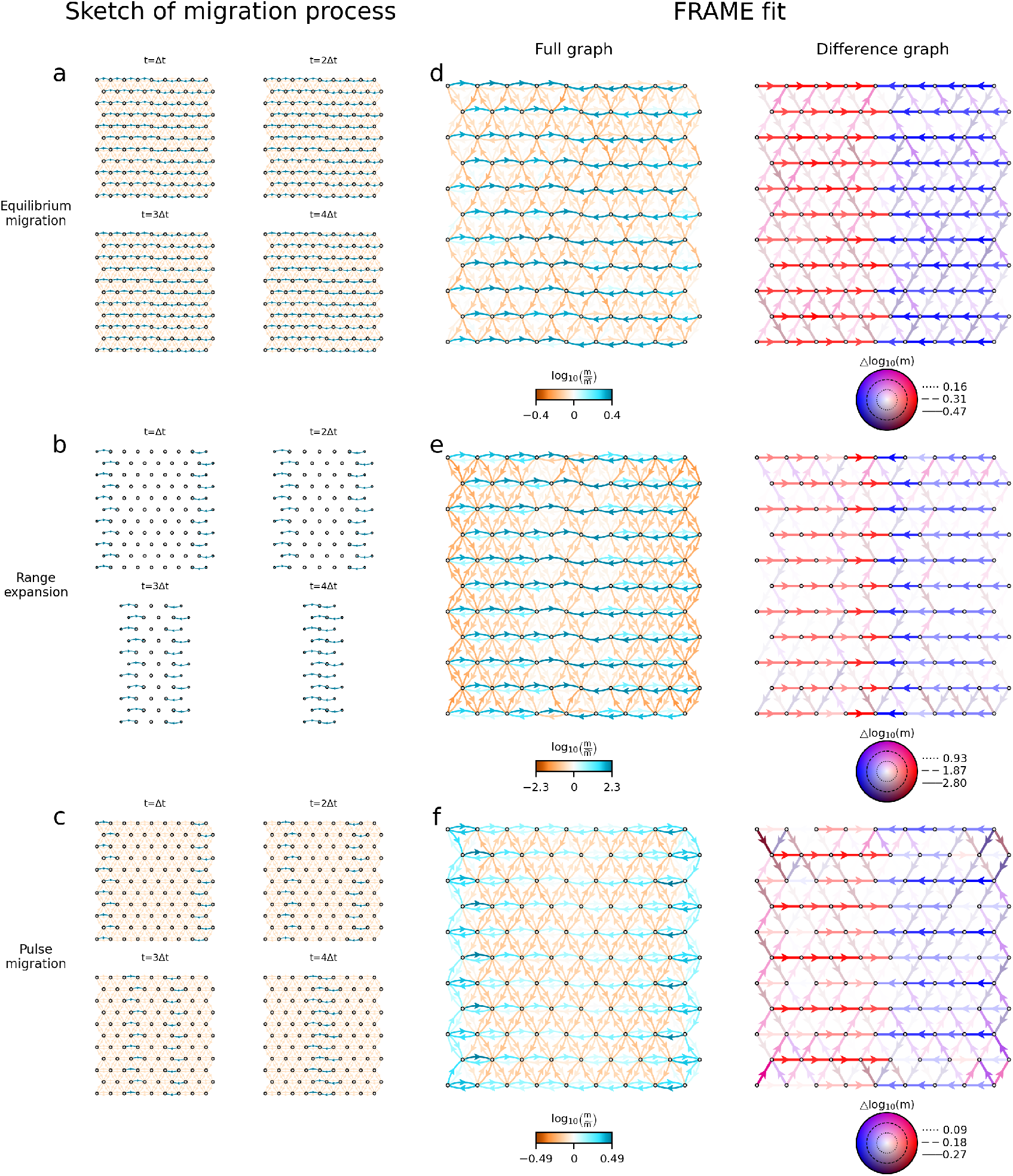
Simulations demonstrate FRAME’s ability to detect asymmetric migration signals even under non-equilibrium conditions. **(a–c)** Illustrations of the simulated migration scenarios (backward in time). **(a)** Equilibrium migration. **(b)** Range expansion. **(c)** Sequential pulse migration. (**d–f**) FRAME’s fit to the corresponding migration settings.

In all three scenarios, FRAME successfully detects signals of directional gene flow and the inferred migration networks reflect the underlying patterns of backward-in-time gene flow. As a point of caution though, the three unique demographic histories all result in difference graphs that are qualitatively similar, highlighting how interpretation of results from real data requires considering multiple possible generating processes.

### Results on simulated data from continuous space models

The above simulations are all carried out under a backward-in-time discrete-stepping-stone model setting. To evaluate the performance of our method under an alternative spatial structure model, we additionally conduct forward-in-time, continuous space, individual-based simulations using `SLiM` (Haller et al., 2025), as illustrated in Fig. 4.

**Figure 4:**
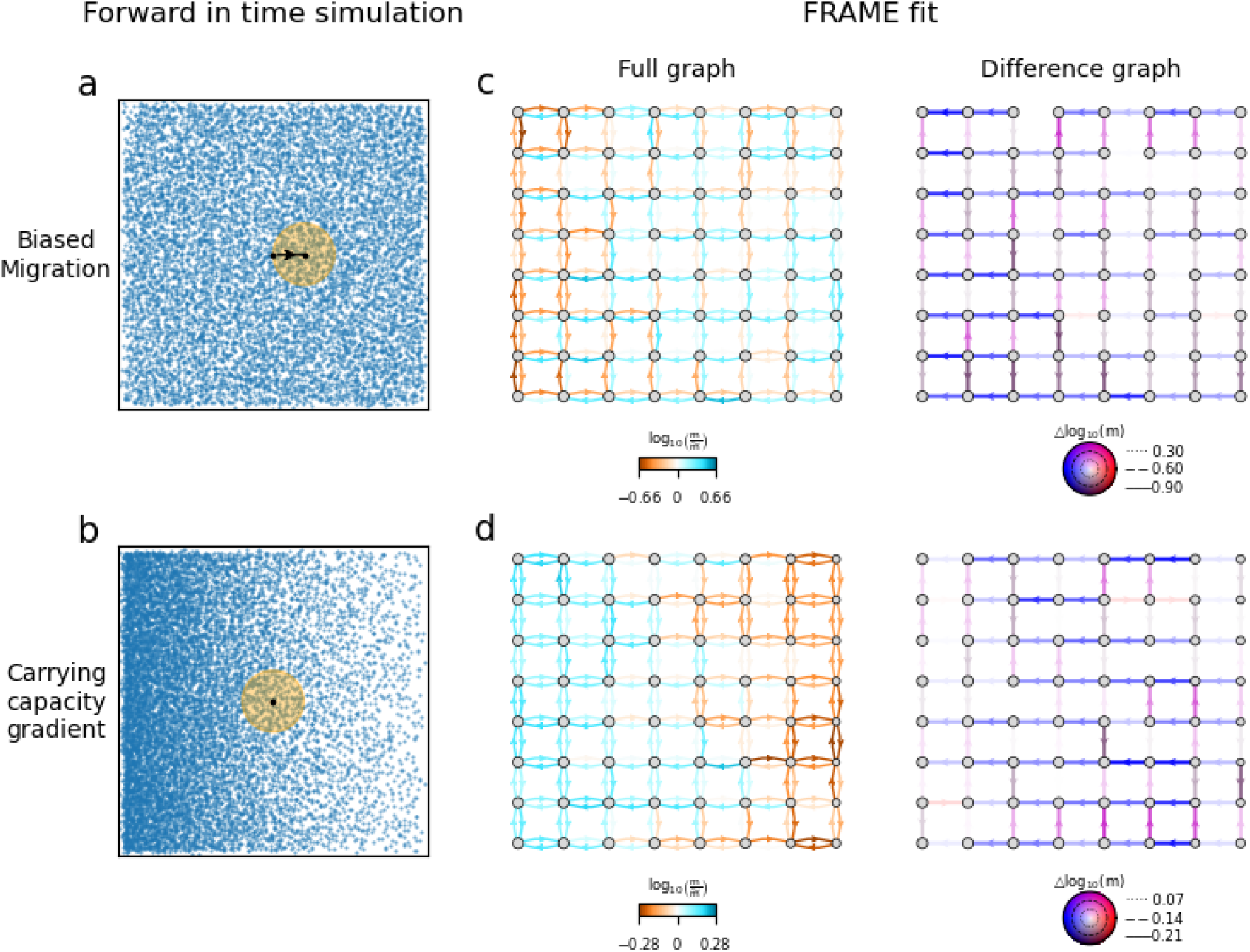
forward-in-time biased migration and local carrying capacity gradient can both lead to backward-in-time migration asymmetry. **(a-b)** Individual-based forward-in-time simulation in a continuous landscape **(a)** Uniform local carrying capacity with uniform left-to-right biased migration **(b)** Decreasing local carrying capacity from left to right with uniform unbiased migration **(c-d)** FRAME’s fit to the corresponding migration settings.

We first consider a scenario with uniform local carrying capacity and uniform left-to-right biased migration, shown in Fig. 4a with sampling carried out in a grid pattern. The simulation settings, grid construction procedure, and sampling protocol are adapted from Lundgren and Ralph (2019), with full details provided in Methods and the Supplementary Note. When applying FRAME to these simulation outputs, the method successfully recovers a clear signal of asymmetric migration: when viewed backward in time, lineages preferentially migrate from right to left, as shown in Fig. 4c.

This scenario helps illustrate a subtle but important aspect of gene flow inference, the relationship between forward and backward-in-time migration dynamics. Let *N*_*i*_ denote the effective population size of deme *i*, and let 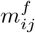 and 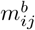 denote the forward and backward-in-time migration rates from deme *i* to deme *j*, respectively. For time-reversible migration-drift scenarios at equilibrium, these quantities satisfy the relationship 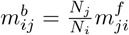, implying that larger populations tend to attract lineages when viewed backward in time. More generally, even if forward-in-time migration rates are symmetric, a spatial gradient in population size can induce directional migration asymmetry backward-in-time.

To illustrate this effect, we perform a second SLiM simulation with uniform unbiased forward-in-time dispersal but a local carrying capacity that decreases from left to right, as shown in Fig. 4b. Using the same square lattice construction and fitting the data with FRAME, we again observe asymmetric gene flow from right to left in the backward-in-time inference (Fig. 4d). Thus, an observed signal of directional migration backward in time is consistent with—but not uniquely diagnostic of—forward-in-time asymmetric migration in the opposite direction. Potential approaches for distinguishing forward-in-time migration asymmetry and gradient of local carrying capacity (or Ne) are discussed further in the Discussion (also see Fig. S11).

### Applications to empirical datasets

We first apply FRAME to a dataset comprising two poplar species, *Populus trichocarpa* (black cottonwood) and *Populus balsamifera* (balsam poplar), from North America (Fig. 5). The dataset was originally generated by Geraldes et al. (2013, 2014) and subsequently adapted by Lundgren and Ralph (2019). The adapted dataset contains genotypes for 431 individuals genotyped using a SNP genotyping array at 30, 576 SNPs originally ascertained in a panel of *P. trichocarpa* accessions (Geraldes et al., 2013). *P. trichocarpa* is distributed primarily along the Pacific coast (red samples, *n* = 423, Fig. 5a), whereas *P. balsamifera* is distributed inland (blue samples, *n* = 8, Fig. 5a). The two species are estimated to have diverged about 75, 000 years ago (Levsen et al., 2012) and admixture/introgression has been inferred (Geraldes et al., 2014). Joint analysis of the two species using this dataset was previously conducted by Lundgren and Ralph (2019) to demonstrate their method, and their method modeled the data with substantial asymmetric gene flow rates, and therefore it provides a natural benchmark for assessing the output of FRAME

**Figure 5.**
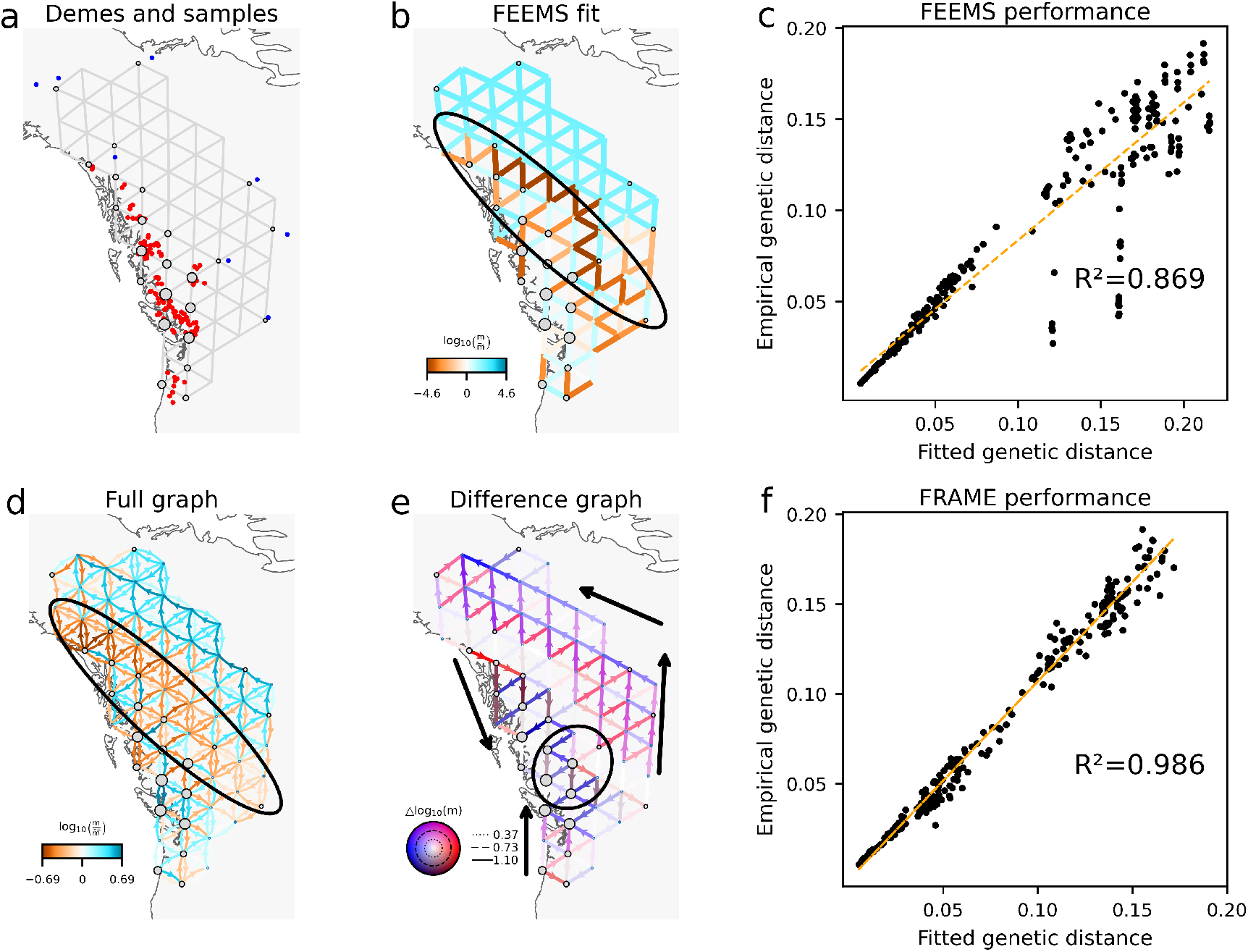
Comparison of FRAME with FEEMS on the *Populus trichocarpa/balsamifera* dataset. **(a)** Demes and samples. *P. trichocarpa* samples are shown in red and *P. balsamifera* samples in blue. **(b)** FEEMS fit. The ellipse highlights reduced migration between the two species. **(c)** FEEMS performance. **(d)** FRAME full graph. The ellipse highlights reduced migration between the two species. **(e)** FRAME difference graph. Black arrows represent inferred asymmetric gene flow; the ellipse highlights spatially diverging lineages. **(f)** FRAME performance.

As an initial analysis, we construct a spatial network of 62 demes and assign each individual to its nearest deme. The constructed demes and samples are shown in Fig. 5a. Both FEEMS and FRAME infer substantially reduced migration between the two species relative to within-species migration (Fig. 5b,d). However, FRAME achieves a markedly improved fit to the data, increasing *R*^2^ from 0.869 to 0.986 (Fig. 5c,f). This improvement likely reflects the fact that FEEMS assumes symmetric migration, whereas FRAME allows for asymmetric gene flow.

Within *P. trichocarpa*, FRAME detects asymmetric gene flow directed from both the northern and southern coastal regions toward the central coast when viewed backward in time. Within *P. balsamifera*, FRAME infers a predominant directional pattern from the southeast toward the northwest. At the location indicated by a circle in Fig. 5, we also observe spatially diverging lineages, consistent with ongoing hybridization.

These patterns of asymmetric gene flow, together with the reduced gene flow between species, are consistent with previous findings (Lundgren and Ralph, 2019) and can be interpreted in light of the known ecological and historical context of the two species. For example, glacial refugia have been proposed in western Alaska for *P. balsamifera* and in coastal British Columbia for *P. trichocarpa*, and, when viewed backward in time, the observed asymmetric gene flow (shown as black arrows in Fig. 5e) can be explained by recently colonized populations receiving more ancestry from these refugial regions than vice versa.

We further compare FRAME with the method of Lundgren and Ralph (2019) using the *P. trichocarpa/balsamifera* dataset, applying the same deme construction and sample assignment procedure (Fig. S6). Under a coarse-resolution model with 9 demes, FRAME produces results comparable to those of Lundgren’s method while requiring substantially less computation time (Fig. S1).

We next apply FRAME to a dataset of *Canis lupus*, North American gray wolves Schweizer et al. (2016); Shastry et al. (2025) with 108 individuals and 17, 729 SNPs, which was previously used as a test case in the development of FEEMS (Fig. S7). In contrast to what we find for the *P. trichocarpa/balsamifera* dataset, accounting for asymmetric migration yields only a modest improvement in model fit, with *R*^2^ increasing from 0.964 to 0.988 using FRAME versus FEEMS (Fig. S7c,f). This result suggests that, relative to the *P. trichocarpa/balsamifera* dataset, symmetric migration provides a more adequate approximation for the gray wolf dataset.

Given that many directional population movements have been inferred from ancient human DNA datasets from Europe (e.g. Lipson et al., 2017; Olalde et al., 2019; Allentoft et al., 2015; Haak et al., 2015), we applied FRAME to assess its behavior on such datasets. We analyzed ancient DNA datasets generated from the 1240k panel of the Allen Ancient DNA Resource (Mallick et al., 2024). Our analyses focused on two broad time periods: (1) 7000**-**3500 BCE, corresponding to the Neolithic expansion across Europe (Lipson et al., 2017; Olalde et al., 2019); and (2) 3500-1500 BCE, marked by the spread of genetic ancestry associated with Steppe pastoralists (Allentoft et al., 2015; Haak et al., 2015). Although the coalescent model underlying FRAME assumes contemporaneous sampling, we used broad time windows to ensure sufficient sample sizes. For comparison, we also examined narrower intervals (4500–3500 BCE and 2500–1500 BCE; see Fig. S8-S9).

To prepare these datasets, we first filter on sample geographic location and time period, followed by additional filters to reduce missingness, remove rare alleles, and reduce LD among markers (see Supplementary Note for details). After the filtering, we obtained a 7000 − 3500 BCE dataset with 975 individuals and 92, 967 SNPs, a 3500-1500 BCE dataset with 1361 individuals and 51, 306 SNPs, a 4500-3500 BCE dataset with 383 individuals and 204, 791 SNPs and a 2500-1500 BCE dataset with 916 individuals and 48, 553 SNPs.

For the 7000-3500 BCE dataset the FRAME difference graph shows an inference of directionally migrating lineages aligned in paths leading towards the area of Anatolia: one path begins in the British Isles and continues southeast; another originates in central-western France and moves eastward; and a third starts in northeast Europe and points southwest. These paths possibly reflect the observed spread of the Linear Pottery Culture (LBK) cultures (Lipson et al., 2017). We also observe directionally migrating lineages following a path that moves northeast along the Northern Mediterranean coastline and towards Anatolia, aligning with the proposed Mediterranean route of Neolithic expansion (Olalde et al., 2019). These results appear consistent with Anatolia as the geographic origin of farmer groups associated with the Mediterranean Cardial and LBK cultures (Olalde et al., 2015). The difference graph (Fig. 6a, right panel) also shows directionally migrating lineages aligned along a path northeastward out of present-day Romania, arcing westward to Central Europe, and looping back to the area of the Balkans. This trajectory may reflect gene flow processes shaped by the Carpathian Mountains, a region inferred to have reduced connectivity in the full migration graph.

**Figure 6:**
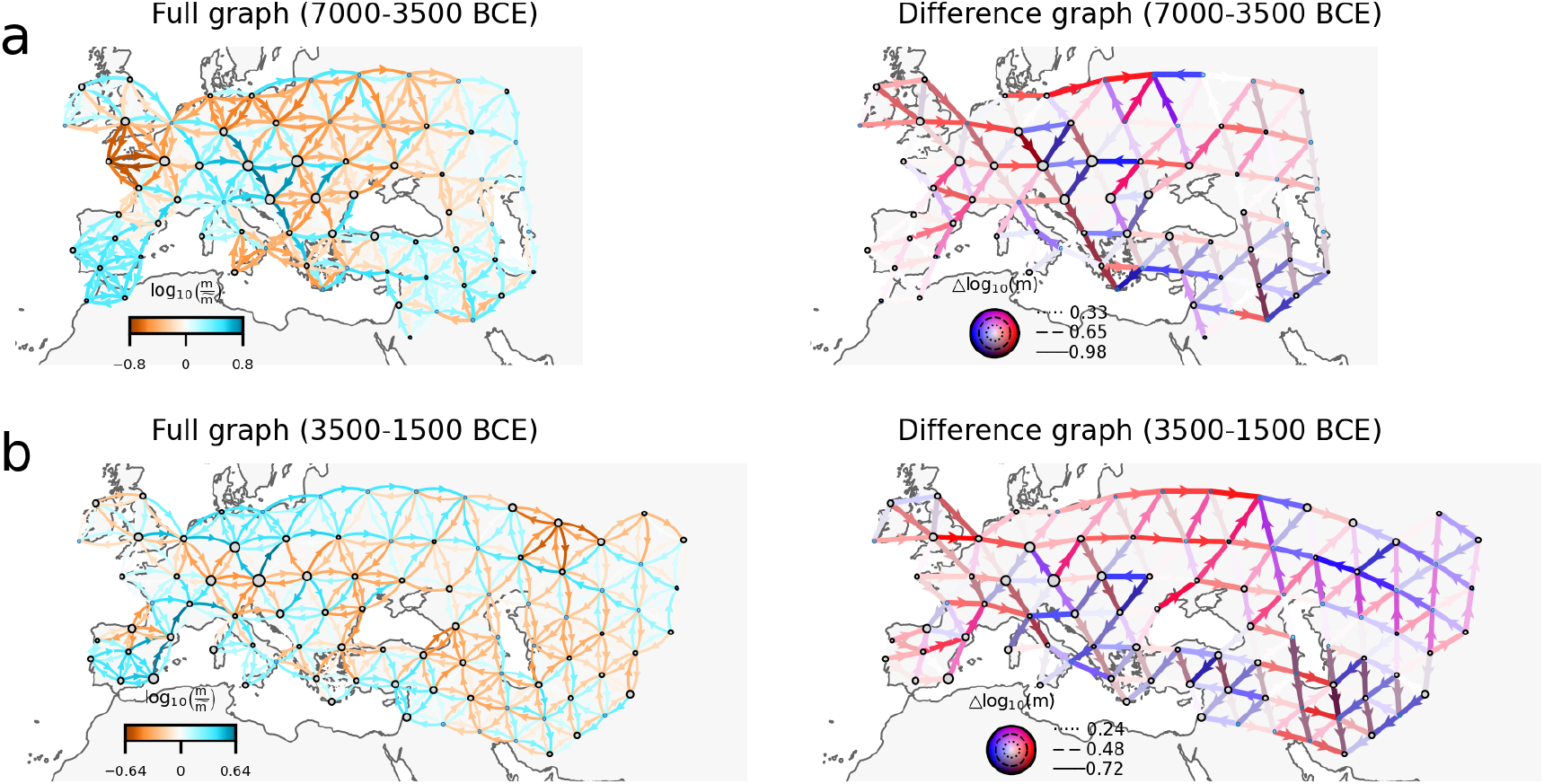
Application of FRAME to aDNA datasets. **(a)** FRAME analysis of Eurasian human populations from 7000-3500 BCE (Neolithic expansion) **(b)** FRAME analysis of Eurasian human populations from 3500-1500 BCE (Steppe expansion).

For the 3500-1500 BCE dataset, the difference graph (Fig. 6b, right panel) highlights strong signals of directionally migrating lineages from Central and Eastern Europe to the Pontic-Caspian steppe, aligning with the westward movement of Yamnaya-related populations (Haak et al., 2015). In the opposite direction, FRAME also detects directionally migrating lineages from Central Asia extending westward to the Pontic-Caspian steppe. This branch of ancestral flow potentially corresponds to movements associated with the Sintashta and Andronovo cultures (Allentoft et al., 2015).

The smaller time window analyses do not fully replicate the features observed in the broader windows, likely due to reduced sample sizes or time-varying gene flow dynamics during these periods. Comparing the 7000–3500 BCE and 4500–3500 BCE analyses (Fig. S8), we observe some shared asymmetries but also notable differences—for example along the Mediterranean, where the narrower window lacks samples from southern France. The 3500–1500 BCE and 2500–1500 BCE analyses (Fig. S9) show more agreement, with the asymmetries plausibly reflecting gene flow out of the Steppe region being a shared signal in both analyses.

## Discussion

In this work, we have developed an algorithm for efficient pairwise coalescent time computation in structured populations at migration-drift equilibrium and applied it for detecting asymmetric migration patterns in spatial models of population structure. By leveraging advanced techniques from computational linear algebra, we can compute expected coalescence times under structured coalescent with asymmetric migration with time complexity *O*(*d*^4^), a task previously limited to *O*(*d*^6^) due to the high computational cost of solving large-scale matrix equations. Our application of this algorithm allows researchers to visualize patterns of spatial structure using directional gene flow parameters, extending beyond previous symmetric approaches and offering new perspectives on population structure. Thus, the work here constitutes a computational advance in structured coalescent methodology and its application to spatial population genomic data.

In our application of the method to empirical datasets, we detect strong signals of effective asymmetric gene flow in both *P. trichocarpa/balsamifera* and ancient human datasets, consistent with known species histories. Applying FRAME to a broader range of species may help determine how widespread asymmetric gene flow is and identify the ecological and evolutionary factors that drive it.

While FRAME performs well in both simulations and empirical applications, several considerations are important when applying the method. First, the spatial network should be constructed carefully. Although FRAME successfully detects asymmetric gene flow in continuous-landscape, individual-based simulations, using excessively fine spatial resolution can be problematic. Unlike FEEMS, FRAME is not computationally feasible for networks with thousands of demes: fitting a 100-deme model requires approximately one hour, while a 300-deme model may require 3–4 days, with runtime scaling as *O*(*d*^4^). Moreover, very fine networks often result in sparse sampling, which reduces power to recover small-scale migration patterns and raises identifiability concerns. In our simulations, at least 25% of demes contained samples, and in all empirical analyses presented here, at least 40% of demes were occupied; when this proportion is substantially lower (e.g., 5–10%), we recommend using a coarser grid. Accordingly, decisions about representing internal habitat discontinuities (e.g., lakes or mountain ranges) should be guided by the biological and geographic context. For empirical applications used in this paper, we typically generate grids automatically and then refine them based on sample locations and geographic features; a comparison between automatically generated and refined networks is shown in Fig. S10.

Second, Lundgren and Ralph (2019) showed that for equation 2, the solution for *γ* and *L* given *T* need not be unique. In stepping-stone–like networks, this identifiability issue is generally less severe due to the reduced number of parameters, although it can still arise when sample locations are extremely sparse. In FRAME, we further impose strong constraints on the relationship between migration and coalescence rates to mitigate identifiability issues. As a consequence, inference of coalescence rates is relatively restricted: while the overall relationship between migration and coalescence rates is often recovered correctly, as is shown in Fig. S4-S5, deviations from the assumed relationship are difficult to infer. We therefore recommend inspecting FRAME results primarily in terms of what they reveal for backward-in-time migration patterns, and view relaxing this restriction as an important direction for future extensions of the method.

Third, translating backward-in-time dynamics into forward-in-time interpretations requires careful consideration. In our simulations, we show that both forward-in-time asymmetric migration and spatial gradients in local carrying capacity can generate asymmetric migration patterns when viewed backward in time. In principle, in a time-reversible system at equilibrium these two scenarios can be differentiated with good estimates of the model parameters by translating the backward-in-time migration rates to forward-in-time migration rates; however, due to expected departures from equilibrium, and our modeling assumptions about the *γ*_*i*_’s, we recommend caution in doing such translation, and favor focusing on the backward-in-time results. Downstream work incorporating other dimensions of the data is likely fruitful, but beyond our scope here. For example, our simulations suggest a potential way to distinguish between these two scenarios using heterozygosity (see Fig. S11). Under forward-in-time uniform left-to-right biased migration with uniform local carrying capacity, heterozygosity tends to increase from left to right, reflecting increased admixture in downstream populations. In contrast, under a left-to-right gradient of decreasing local carrying capacity with uniform unbiased migration, heterozygosity decreases from left to right, as downstream populations are smaller and thus harbor less genetic diversity. Although the relationships among heterozygosity, migration patterns, and coalescence rates can be more complex in general settings, this contrast may provide a useful first-step diagnostic for distinguishing between these two scenarios.

Finally, because FRAME assumes migration-drift equilibrium conditions, it has limited capacity to explicitly model temporally varying migration dynamics. By averaging genomic signals over the coalescent timescale, FRAME simplifies calculations, but as a result it cannot capture detailed temporal changes in migration patterns. Consequently, FRAME’s results depend significantly on the timing of past migration events. For example, if a population transitions from migration pattern A to pattern B, samples collected shortly after this transition may predominantly reflect the earlier pattern, whereas later samples would reflect the new equilibrium.

An illustrative example of this effect is the influence of divergence time on inference. In general, divergence between populations or subpopulations—when accompanied by geographic isolation and reduced hybridization—can generate a strong signal of reduced gene flow, as observed in the *P. trichocarpa/balsamifera* example. However, this signal is only detectable if the divergence has persisted for a sufficiently long time. When divergence occurs recently, the genomic signature of reduced gene flow remains weak and may be difficult to detect. In Fig. S12, we use `msprime` simulations to show that the signal of reduced gene flow induced by genetic isolation during divergence becomes detectable only when the divergence event occurred sufficiently far in the past.

Furthermore, migration events often vary in both timing and duration across regions, causing local genomic signals to emerge and decay asynchronously. Because FRAME summarizes these signals into time-averaged migration patterns, it is not well suited for resolving the temporal ordering of events in non-equilibrium scenarios. Complementary phylogenetic approaches that allow for more flexible demographic histories (Hey and Nielsen, 2007; Yang and Rannala, 2010; Gronau et al., 2011; Excoffier et al., 2013) can help address these questions, particularly when the number of demes is small.

These considerations emphasize the importance of careful dataset construction and cautious interpretation of results. While FRAME effectively identifies fine-scale migration patterns, its equilibrium-based approach constrains its ability to reconstruct deeper evolutionary histories. To improve the reconstruction of non-equilibrium dynamics, two possible future directions are: 1) develop methods that analyze time series data; 2) integrate haplotype-based information, akin to how the MAPS method (Al-Asadi et al., 2019) uses the length distribution of ‘long pairwise shared coalescence segments’ (also known as ‘identity by descent’ tracts). While either approach would require solving additional modeling and computational challenges, it would move the field closer to an ideal method that could reconstruct high-resolution spatiotemporal maps of asymmetric gene flow.

## Methods

### The expected pairwise coalescence times

The matrix *T*, whose (*i, j*)_*th*_ entry represents the expected coalescence time between a haploid individual from deme *i* and another from deme *j*, given the migration rates and coalescence rates, can be solved using equation (1). However, solving this equation with a direct vectorized approach has time complexity *O*(*d*^6^), which is computationally prohibitive for fine-resolution analyses. We reduce the computational complexity by leveraging the similarity between this equation and continuous-time Lyapunov equations (Gajic and Qureshi, 2008). In short, let *G* be an arbitrary symmetric matrix, we solve the more general equation

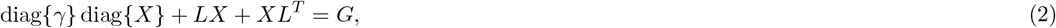

by solving a series of simpler singular continuous-time Lyapunov equations. *LX*+*XL*^*T*^ = *G* and *LX*+*XL*^*T*^ = *E*_*i*_, *i* = 1, 2, …, *d* in the least square sense, where *E*_*i*_ is the matrix with its (*i, i*)_th_ entry equal to 1 and all other entries equal to 0. The solution can then be expressed as a linear combination of the solutions to these simpler equations, along with some constants. The coefficients of this linear combination can be determined either using the method of undetermined coefficients or through a step-by-step low-rank update approach, which offers greater computational robustness. This approach reduces the complexity to *O*(*d*^4^). The detailed algorithm is provided in the Supplementary Note.

### The penalized likelihood

Assume we have *o* observed sample groups with corresponding *o* × *d* assignment matrix *J* if sample group *i* is in deme 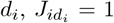, otherwise 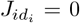. Assume in sample group *i*, we have *n*_*i*_ copies of haploid data on *p* SNPs (every diploid counts as two copies). Let allele state of sample group *i*, locus *j*, copy *k* be *Z*_*ijk*_ (coded as 0 or 1), and let 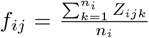 be the allele frequency at sample group *i*, locus *j*, we can define vectors *f*_*j*_ = (*f*_1*j*_, …, *f*_*oj*_)^*T*^, *j* = 1, 2, …, *p*. Each vector *f*_*j*_ can be viewed as an independent realization of the random vector of allele frequencies assuming perfect segregation (justified in the Supplementary Note). Let *T*_mrca_ and *T*_tot_ be the expected height and expected total branch length of the coalescent tree across all samples; *K* be a *o* × 1 vector such that its *i*_th_ entry is 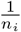, the mean of the random vector of allele frequencies is then 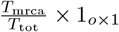 and the covariance matrix is given by (Supplementary Note)

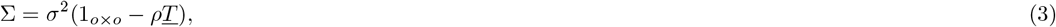

where 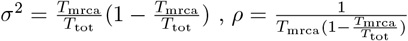 and <u>*T*</u> = *JTJ*^*T*^ − diag{*JTJ*^*T*^} diag{*K*}.

To remove the mean and focus on the covariance structure, we apply a contrast matrix *C* ∈ *R*^*o*−1*×o*^ (Supplementary Note). Assume each *f*_*j*_ obeys a multivariate normal distribution, we then have

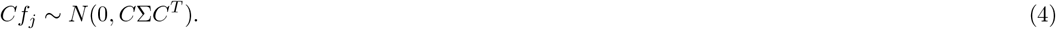

The sample covariance matrix 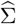 after the transformation obeys a Wishart distribution

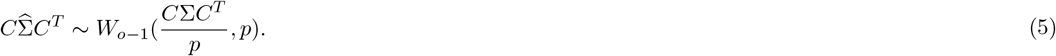

Notice that the transformed covariance matrix can be expressed as

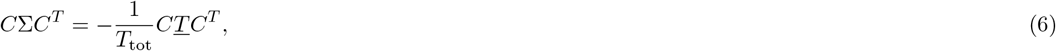

where *T*_tot_, the expected total coalescence time, acts as a nuisance parameter. *T*_tot_ is a complex function of the underlying model and the sample configuration across demes, making it challenging to disentangle directly during inference.

To address this, we absorb it into other parameters by redefining *L*^*′*^ = *T*_tot_*L*; *γ*^*′*^ = *T*_tot_*γ*, and 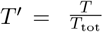. This absorption simplifies the parameterization and can be justified within the maximum likelihood framework (see Supplementary Note). Importantly, the structured coalescent equation remains valid under this rescaling. Thus, we can drop the primes and proceed by treating *L*^*′*^, *γ*^*′*^, *T* ^*′*^ as the new *L, γ, T*, respectively.

After renormalizing the time scale and removing constants, the negative log-likelihood is expressed as:

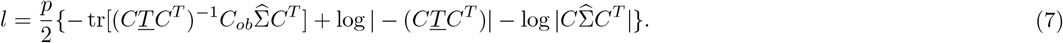

Let Ω = {(*i, j*)|*m*_*ij*_ *>* 0} represent the edge set, and let Ω_*i*_ = {(*i, t*) ∈ Ω} ∪ {(*s, i*) ∈ Ω} denote the edges connected to node *i*. Then, the penalty can be written as *λ*_*m*_Ψ, where

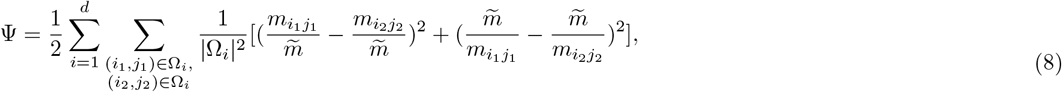

and 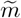 is the geometric mean of all the nonzero migration rates.

This penalty function controls the smoothness of the fitting and penalizes both low and high migration rates equally, preventing ill-conditioned situations where parameters vary across many orders of magnitude.

The objective function is then *l*+*λ*_*m*_Ψ, with *l* defined in 7 and Ψ defined in equation (8). The optimization problem then is to find parameters 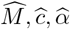 that minimize *l* + *λ*_*m*_Ψ.

### Optimization

We infer the migration rates matrix 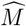 by solving the following optimization problem:

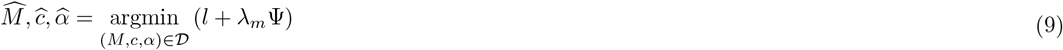

where 𝒟= 𝒟 _ℳ_ *R*^+^ [0, 1], 𝒟 _ℳ_ = {*M m*_*ij*_ *>* 0 if (*i, j*) Ω, *m*_*ij*_ = 0 if (*i, j*) */* Ω }.

Similarly to FEEMS, we solve the optimization using the L-BFGS method (Byrd et al., 1995). A crucial step in this optimization is the efficient computation of the gradient, which requires solving equations of the form (Supplementary Note):

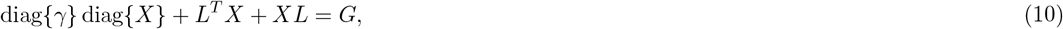

where *G* is an arbitrary symmetric matrix. Although this equation is different from equation (2), the computational techniques used to solve (2) can be applied here with appropriate modifications, as detailed in the Supplementary Note. The time complexity of computing the gradient remains *O*(*d*^4^).

### Cross-validation

The observed demes are split into a training set and a testing set. For each candidate value of *λ*_*m*_, we fit the model on the training set and predict allele frequencies (after contrasting) for the testing set. Specifically, let 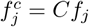 and ∑^*c*^ = *C*∑*C*^*T*^ . Then:

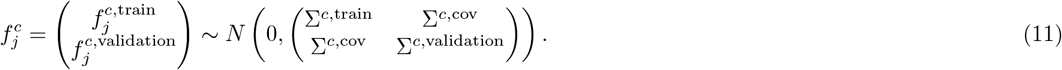

Prediction can then be made using:

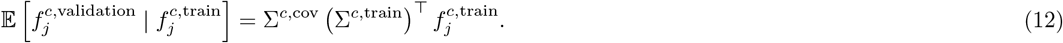

The value of *λ*_*m*_ that minimizes the prediction error is selected, and the model is then refit to the entire dataset using this optimal *λ*_*m*_.

Leaving one deme out at a time for cross-validation is the default, though for datasets with a large number of observed demes, using 10-fold cross-validation can improve efficiency.

### Simulations

The equilibrium migration simulations in Fig. 2 are performed using `msprime` (Baumdicker et al., 2022). For each simulation, we generate a dataset containing 10^5^ SNPs across all sampled individuals. Under dense sampling, individuals are sampled from every deme, whereas under sparse sampling, each deme is selected independently with probability 0.25; when selected, 10 individuals are sampled from that deme.

In all large-scale pattern simulations shown in Fig. 2(d–l), the ground-truth graph uses deep blue edges to represent migration rates that are 3 times greater than the baseline *m*_base_, which is depicted with shallow orange edges. We set the baseline migration rates and local population sizes (*N*) so that *Nm*_base_ = 0.1.

In the small-scale pattern simulations shown in Fig. 2(a–c), assuming the base migration rate (i.e., the migration rate of the edges in the middle row) is 1, the edges in the lower half of the graph have base migration rates of 3, while the edges in the upper half have base migration rates of 0.3. Additionally, all patterns exhibit migration rates that are 10 times higher than the surrounding base migration rates. Specifically, the patterns in the upper half of the graph have a migration rate of 3, while patterns in the lower half have migration rates of 30. This configuration ensures that all patterns are clearly distinguishable from the background while also maintaining geographic differentiation within the background itself.

The non-equilibrium simulations in Fig. 3 are also performed using `msprime`. The settings are more complex and are detailed in the Supplementary Note.

The forward-in-time individual-based simulations in Fig. 4 and Fig. S11 are performed using `SLiM` (Haller et al., 2025). The detailed settings are shown in the Supplementary Note.

The split and divergence simulations in Fig. S12 are performed using `msprime`. Backward in time, we assume that during [0, *T*_div_), the left four columns and right four columns of demes form two subpopulations. Within each subpopulation, local migration rates between neighboring demes (*m*) and local effective population sizes (*N*) satisfy *Nm* = 1, and there is no gene flow between the two subpopulations. At time *T*_div_, all lineages enter a panmictic population whose effective population size is the total effective population size of all demes. *T*_div_ is set to 0.01 for Fig. S12a–b and 3 for Fig. S12c–d.

### Empirical data sets

We have applied FRAME to the following datasets: a *P. trichocarpa/balsamifera* dataset with 431 individuals and 30, 756 SNPs, a North American gray wolf dataset with 108 individuals and 17, 729 SNPs, a 7000-3500 BCE ancient human dataset with 975 individuals and 92, 967 SNPs, a 3500-1500 BCE ancient human dataset with 1361 individuals and 51, 306 SNPs, a 4500-3500 BCE ancient human dataset with 383 individuals and 204, 791 SNPs and a 2500-1500 BCE ancient human dataset with 916 individuals and 48, 553 SNPs.

The *P. trichocarpa/balsamifera* dataset was originally generated by Geraldes et al. (2014) and subsequently adapted by Lundgren and Ralph (2019), and we use the adapted version. The North American gray wolf dataset was generated by Schweizer et al. (2016), with corrections introduced by Shastry et al. (2025); we use the corrected dataset. Missing genotypes in these datasets are imputed using average allele frequencies.

Filtering procedures for the ancient human datasets are more complex due to the low genotype rates typical of ancient DNA samples. Additional details on dataset construction and filtering are provided in the Supplementary Note.

### Extended Graph Representations

Beyond the primary representations—the full graph and difference graph—two other options for visualizing the migration dynamics are what we term a ‘base graph’ and ‘summary graph’ (see Fig. S13-S19).

Fig. S13 presents these additional visualizations. The ‘base graph’ illustrates mean migration rates between connected demes on a log scale, providing a complementary perspective to the difference graph. The qualitative similarity between the base graph and FEEMS outputs highlights the broader relationship between these models. The ‘summary graph’ aggregates backward migration rates at each node into a vector, summarizing overall trends in directionally migrating lineages.

Together, these representations enhance the interpretability of migration patterns and offer diverse tools for exploring population structure. A comprehensive set of visualizations, including all four graph representations and related demographic parameters for the empirical datasets, is presented in Fig. S14–S19.

Notably, these figures all show that coalescence rates are inferred as relatively uniform rather than inversely proportional to the stationary distribution, suggesting that local effective population sizes in these datasets may be largely independent of migration.

### Network Construction

FRAME leverages an automatic method to construct the network similar to that used in FEEMS (Marcus et al., 2021), which generates a dense triangular lattice over the study region. We define the outer boundary using the convex hull of sample locations with a layer of buffer, which is manually refined to align with the species’ habitat and relevant geographic features. To ensure uniform spatial coverage, we intersect the boundary with a precomputed global triangular grid projected onto the Earth’s surface (Sahr et al., 2003). Multiple grid resolutions are available, allowing users to evaluate model performance across different spatial scales. Alternatively, users can manually design the network and assign samples to demes using empirical knowledge.

### URLS

Software implementing FRAME is available at https://github.com/ShenHaotv/frame. The code to reproduce all results, along with the complete simulation data, is available at https://github.com/ShenHaotv/frame-analysis.

## Acknowledgements

FRAME is built upon the FEEMS codebase, which was originally developed by Joe Marcus and Wooseok Ha, and further expanded upon and developed by Vivaswat Shastry. We would further like to thank Vivaswat Shastry for helpful conversations and for advice on software development. We also thank members of the Novembre Lab collectively for input on figure design. The research was supported by funding from NIH NIGMS grant R35-GM149521 to JN.

