## Supplementary Note for "Fine-Resolution Asymmetric Migration Estimation"

**1 Mathematical details of the FRAME method** **2**

### 1 Mathematical details of the FRAME method

#### 1.1 Overview

The structured coalescent process combines backward migration among demes and coalescence events when lineages co-occur in the same deme. This process canbe modeled in terms of a weighted directed graph. The nodes of the graph are denoted by $\{1, 2, ..., d\}$, each representing a deme. The weight associated with the edge from node $i$ to node $j$ is $m_{ij}$, representing the backward migration rate from $i$ to $j$. Biologically, $m_{ij}$ is the rate at which deme $i$ receives ancestry from deme $j$. The Laplacian of the weighted directed graph is then

$$L = \begin{pmatrix} \sum_{i\neq 1} m_{1i} & -m_{12} & \dots & -m_{1d} \\ -m_{21} & \sum_{i\neq 2} m_{2i} & \dots & -m_{2d} \\ \vdots & \vdots & \dots & \vdots \\ -m_{d1} & -m_{d2} & \dots & \sum_{i\neq d} m_{di} \end{pmatrix}.$$

The transition rate matrix of the jump process is defined as $Q = -L$. The set of migration rates is encoded in the weighted adjacency matrix $M$, which is given by $M = Q - \operatorname{diag} Q$.

The coalescence rate per deme is represented by the vector $\gamma = (\gamma_1, \gamma_2, ..., \gamma_d)$, where $\gamma_i$ is the coalescence rate of deme $i$. Rather than fit the coalescent rates per deme as fully independent parameters, we take an approach that has the potential to favor coalescent rates that are consistent with the stationary distribution implied by the migration process (as one might expect when a system is truly at equilibrium). Let $\pi = (\pi_1, \pi_2, ..., \pi_d)^T$ denote the stationary distribution of the of the backward migration process. We then model the coalescence rate in deme $i$ as:

$$\gamma_i = \frac{c}{\pi_i^\alpha}, \tag{S1}$$

where $c$ is a constant scaling parameter, and $\alpha \in [0, 1]$ is a parameter that controls the relationship between coalescence rates and migration dynamics. This formulation captures a continuum of scenarios—from deme sizes being entirely independent of migration rates and constant ($\alpha = 0$) to being fully determined by the stationary distribution ($\alpha = 1$).

In this framework, the parameters $(M, c, \alpha)$ can be used to compute the matrix of expected pairwise coalescent times ($T$) and in turn the negative log likelihood $l$. This can be conceptualized as moving through the sequence $(M, c, \alpha) \to (M, \gamma) \to T \to l$. Here $T$ is the matrix where the $(i, j)_{\text{th}}$ entry is the expected pairwise coalescence time between haploid samples from deme $i$ and $j$, and $l$ denotes the negative log likelihood (up to a constant). Computing $T$ consistent with a given $M$ and $\gamma$ involves solving the equation (Strobeck, 1987)

$$\operatorname{diag}\{\gamma\} \operatorname{diag}\{T\} + LT + TL^T = 1_{d\times d}, \tag{S2}$$

where $1_{d\times d}$ is a matrix of ones. Section 1.4 will provide a detailed explanation of the solution methods for this equation.

The final step involves formulating the negative log likelihood. Since our approach employs a penalized maximum likelihood method, the objective function is expressed as $l(M, c, \alpha) + \lambda_m \Psi(M)$, where $\lambda_m \Psi(M)$ serves as a penalty term to regulate overfitting using $M$. The optimization task is to minimize this objective over an appropriate domain $D$. The explicit form of $l$, $\lambda_m \Psi$ and $D$, along with the rationale behind this construction are discussed in section 1.3. Section 1.2 examines the covariance structure of allele frequencies, providing essential groundwork for understanding section 1.3.

To solve the optimization problem, we utilize the L-BFGS algorithm (Byrd et al., 1995). A crucial component of this method is gradient computation, which presents notable challenges in our context. Section 1.5 elaborates on the procedures for calculating the gradient efficiently.

To select the optimal hyperparameter $\lambda_m$, we employ cross-validation based on the model's predictive performance in estimating allele frequencies. The details of this cross-validation procedure are provided in the Methods section of the main text; for completeness, we restate it in section 1.6.

#### 1.2 Covariance matrix of allele frequencies

Assume we have $o$ observed sample groups with corresponding $o \times d$ assignment matrix $J$ if sample group $i$ is in deme $d_i$, $J_{id_i} = 1$, otherwise $J_{id_i} = 0$. Assume in sample group $i$ , we have $n_i$ copies of haploid data on $p$ SNPs (every diploid counts as two copies). Let allele state of sample group $i$ , locus $j$ , copy $k$ be $Z_{ijk}$ (coded as 0 or 1). As in McVean (2009), we condition on the event $S$ that the SNP segregates in the sample due to a single mutation and take the limit as the mutation rate $\theta$ approaches zero. The expected allele state is then

$$E(Z_{ijk}|S) = \frac{\Pr(Z_{ijk} = 1, S)}{P(S)} = \lim_{\theta \to 0} \frac{E(\theta t_{\text{mrca}} e^{-\theta t_{\text{tot}}})}{E(\theta t_{\text{tot}} e^{-\theta t_{\text{tot}}})} = \frac{T_{\text{mrca}}}{T_{\text{tot}}}, \tag{S3}$$

where the $t$'s are random coalescence times and the $T$'s are their expectations, under the coalescent process, or be more specific, $T_{\text{mrca}}$ and $T_{\text{tot}}$ are the expected height and expected total branch lengthof the coalescent tree across all samples.

Similarly, the probability that two randomly sampled (with repetition) alleles share a derived mutation is

$$E(Z_{ijk}^2|S) = \frac{T_{\text{mrca}}}{T_{\text{tot}}}, \tag{S4a}$$

$$E(Z_{ijk_1} Z_{ijk_2}|S) = \frac{T_{\text{mrca}} - T_{d_i d_i}}{T_{\text{tot}}}, \quad k_1 \neq k_2, \tag{S4b}$$

$$E(Z_{i_1 jk_1} Z_{i_2 jk_2}|S) = \frac{T_{\text{mrca}} - T_{d_{i_1} d_{i_2}}}{T_{\text{tot}}}, \quad i_1 \neq i_2. \tag{S4c}$$

Combining equation (S3) and (S4), the covariance structure of allele states is then

$$\text{Var}(Z_{ijk}^2|S) = \frac{T_{\text{mrca}}}{T_{\text{tot}}}(1 - \frac{T_{\text{mrca}}}{T_{\text{tot}}}), \tag{S5a}$$

$$\text{Cov}(Z_{ijk_1}, Z_{ijk_2}|S) = \frac{T_{\text{mrca}}}{T_{\text{tot}}}(1 - \frac{T_{\text{mrca}}}{T_{\text{tot}}}) - \frac{T_{d_i d_i}}{T_{\text{tot}}}, \quad k_1 \neq k_2, \tag{S5b}$$

$$\text{Cov}(Z_{i_1 jk_1}, Z_{i_2 jk_2}|S) = \frac{T_{\text{mrca}}}{T_{\text{tot}}}(1 - \frac{T_{\text{mrca}}}{T_{\text{tot}}}) - \frac{T_{d_{i_1} d_{i_2}}}{T_{\text{tot}}}, \quad i_1 \neq i_2. \tag{S5c}$$

Now let $f_{ij} = \frac{\sum_{k=1}^{n_i} Z_{ijk}}{n_i}$ be the allele frequency for sample group $i$, locus $j$. We can define vectors $f_j = (f_{1j}, ..., f_{oj})^T, j = 1, 2, \ldots, p$, which can be viewed as independent realizations of the random vector of allele frequencies assuming perfect segregation. Let $K$ be a $o \times 1$ vector such that its $i_{\text{th}}$ entry is $\frac{1}{n_i}$, from equation (S5) and definition of $f_j$, the covariance matrix for each $f_j$ can be written as

$$\Sigma = \sigma^2(1_{o \times o} - \rho \underline{T}), \tag{S6}$$

where

$$\sigma^2 = \frac{T_{\text{mrca}}}{T_{\text{tot}}}(1 - \frac{T_{\text{mrca}}}{T_{\text{tot}}}), \tag{S7a}$$

$$\rho = \frac{1}{T_{\text{mrca}}(1 - \frac{T_{\text{mrca}}}{T_{\text{tot}}})}, \tag{S7b}$$

$$\underline{T} = JTJ^T - \text{diag}\{JTJ^T\} \, \text{diag}\{K\}. \tag{S7c}$$

#### 1.3 The penalized likelihood function

To remove the mean and focus on the covariance structure, we apply a contrast matrix $C \in \mathbb{R}^{o-1\times o}$, where $C$ is a matrix of rank $o-1$ whose rows sum to 0 . In practice, we construct $C$ by choosing one column and setting it to $-1$. In each row, we then select a unique column, distinct from the chosen column, and set the corresponding entry to 1.

Assume each $f_j$ obeys a multivariate normal distribution, the transformed data is distributed as

$$Cf_j \sim N(0, C\Sigma C^T). \tag{S8}$$

The sample covariance matrix $\widehat{\Sigma}$ after the transformation then obeys a Wishart distribution

$$C\widehat{\Sigma}C^T \sim W_{o-1}(\frac{C\Sigma C^T}{k_p}, k_p), \tag{S9}$$

where $k_p$ represents the degrees of freedom and equals $p$ (the number of SNPs) under perfect independence of loci (see EEMS, Petkova et al., 2016).

The negative log likelihood can then be expressed as

$$\begin{aligned} l_{\text{raw}} = &\frac{k_p}{2}\{\text{tr}[(C\Sigma C^T)^{-1}C_{ob}\widehat{\Sigma}C^T] + \log|(C\Sigma C^T)| - \log|C\widehat{\Sigma}C^T|\} \\ &+ \log[\Gamma_{o-1}(\frac{k_p}{2})] - \frac{k_p(o-1)}{2}\log\frac{k_p}{2} + \frac{o}{2}\log|C\widehat{\Sigma}C^T|. \end{aligned} \tag{S10}$$

Let $\Omega = \{(i,j)|m_{ij} > 0, i = 1,\ldots,d, j = 1,\ldots,d\}$ denote the edge set, and let $\Omega_i = \{(i,t) \in \Omega\} \cup \{(s,i) \in \Omega\}$ denote the edges connected to node $i$. The penalty term is then written as $\lambda_m\Psi$, where

$$\Psi = \frac{1}{2}\sum_{i=1}^{d}\sum_{\substack{(i_1,j_1)\in\Omega_i,\\(i_2,j_2)\in\Omega_i}} \frac{1}{|\Omega_i|^2}[(\frac{m_{i_1j_1}}{\widetilde{m}} - \frac{m_{i_2j_2}}{\widetilde{m}})^2 + (\frac{\widetilde{m}}{m_{i_1j_1}} - \frac{\widetilde{m}}{m_{i_2j_2}})^2], \tag{S11}$$

and $\widetilde{m}$ is the geometric mean of all the nonzero migration rates across the whole graph. The summand penalizes large differences between edge weights $m_{i_1j_1}$ and $m_{i_2j_2}$ that share a common node.

The migration rates matrix $\widehat{M}$ is estimated by solving the following penalized likelihood optimization problem:

$$\widehat{M}, \widehat{c}, \widehat{\alpha}, \widehat{k}_p = \underset{(M,c,\alpha,k_p)\in\mathcal{D}_{\text{raw}}}{\text{argmin}} (l_{\text{raw}} + \lambda_m\Psi) \tag{S12}$$

where $\mathcal{D}_{\text{raw}} = \mathcal{D}_{\mathcal{M}} \times R^+ \times [0,1] \times (0,p]$, $\mathcal{D}_{\mathcal{M}} = \{M|m_{ij} > 0 \text{ if } (i,j) \in \Omega,\ m_{ij} = 0 \text{ if } (i,j) \notin \Omega\}$.

Directly solving this raw optimization problem is difficult and we do several simplifications.

The first simplification involves replacing $k_p$ with $p$ in the negative log-likelihood function, even when there is no perfect independence of loci. Consider the function

$$F(k_p) = \log[\Gamma_{o-1}(\frac{k_p}{2})] - \frac{k_p(o-1)}{2}\log\frac{k_p}{2}, \tag{S13}$$

we would like to show that

$$\lim_{k_p\to\infty} F(k_p) - F(k_p - 1) = \frac{1-o}{2} \tag{S14}$$

Using the definition of multivariate gamma function that

$$\Gamma_a(x) = \pi^{\frac{a(a-1)}{4}}\prod_{i=1}^{a}\Gamma(x + \frac{1-i}{2}), \tag{S15}$$

the difference of the first term in $F(k_p)$ is

$$\log[\Gamma_{o-1}(\frac{k_p}{2})] - \log[\Gamma_{o-1}(\frac{k_p-1}{2})] = \log\frac{\Gamma_{o-1}(\frac{k_p}{2})}{\Gamma_{o-1}(\frac{k_p-1}{2})} = \log\frac{\Gamma(\frac{k_p}{2})}{\Gamma(\frac{k_p-o+1}{2})}. \tag{S16}$$

For the second term, we have

$$k_p\log\frac{k_p}{2} - (k_p-1)\log\frac{(k_p-1)}{2} = \log\frac{k_p}{2} + \log(1+\frac{1}{k_p-1})^{k_p-1}. \tag{S17}$$

Combining the above two equations and applying Stirling's formula yields

$$\begin{aligned}&\lim_{k_p\to\infty} F(k_p) - F(k_p-1)\\ &= \lim_{k_p\to\infty}\log\frac{\Gamma(\frac{k_p}{2})}{\Gamma(\frac{k_p-o+1}{2})(\frac{k_p}{2})^{\frac{o-1}{2}}(1+\frac{1}{k_p-1})^{\frac{(o-1)(k_p-1)}{2}}}\\ &= \frac{1-o}{2}\end{aligned} \tag{S18}$$

This means for large $k_p$ the negative log likelihood function (neglecting constants) approximates

$$\frac{k_p}{2}[-\operatorname{tr}[(C\underline{T}C^T)^{-1}C\widehat{\Sigma}C^T] + \log|-(C\underline{T}C^T)| - \log|C\widehat{\Sigma}C^T| - \frac{o-1}{2}] \tag{S19}$$

This asymptotic linearity means that for large $k_p$, the optimization problem is largely independent of $k_p$.Therefore we can replace $k_p$ with $p$ and and the resulting constant can be absorbed into the penalty parameter $\lambda_m$. The updated optimization problem becomes

$$\widehat{M}, \widehat{c}, \widehat{\alpha} = \underset{(M,c,\alpha)\in\mathcal{D}}{\operatorname{argmin}}\ (l_p + \lambda_m\Psi) \tag{S20}$$

where $\mathcal{D} = \mathcal{D}_{\mathcal{M}} \times R^+ \times [0,1]$ and

$$l_p = \frac{p}{2}\{\operatorname{tr}[(C\Sigma C^T)^{-1}C_{ob}\widehat{\Sigma}C^T] + \log|(C\Sigma C^T)| - \log|C\widehat{\Sigma}C^T|\} \tag{S21}$$

The second simplification involves handling the term $C\Sigma C^T = -\frac{1}{T_{\text{tot}}}C\underline{T}C^T$, where $T_{\text{tot}}$ is the expected total coalescence time. $T_{\text{tot}}$ is a complicated function of the underlying model and the samples and is in general very difficult to compute. To address this, we absorb it into the model by redefining $L' = T_{\text{tot}}L$; $\gamma' = T_{\text{tot}}\gamma$, and $T' = \frac{T}{T_{\text{tot}}}$. This adjustment is justified within the maximum likelihood framework. To see this, we replace $T_{\text{tot}}$ with its MLE given other parameters. The derivative of $l_p$with respect to $T_{\text{tot}}$ is $\frac{p}{2}\{\operatorname{tr}[(-C\underline{T}C^T)^{-1}C\widehat{\Sigma}C^T] - \frac{o-1}{T_{\text{tot}}}\}$. Setting it to be 0, we obtain

$$T_{\text{tot}}^{\text{MLE}} = \frac{o-1}{\operatorname{tr}[(-C\underline{T}C^T)^{-1}C\widehat{\Sigma}C^T]}. \tag{S22}$$

Replacing $T_{\text{tot}}$ in equation (S21) with this expression, the negative log likelihood function (neglecting constants) becomes

$$l_{\text{MLE}} = \frac{p}{2}\{(o-1)\log(\operatorname{tr}[(-C\underline{T}C^T)^{-1}C\widehat{\Sigma}C^T]) + \log|-C\underline{T}C^T|\} \tag{S23}$$

Notice that both $l_{\text{MLE}}$ and $\lambda_m\Psi$ are invariant under the scaling of $(M,c)$ since they are both dimensionless quantity. That means for any $(M,c,\alpha)$, we can define scaled parameters $M' = T_{\text{total}}^{\text{MLE}}M$, $c' = T_{\text{total}}^{\text{MLE}}c$. The new parameters $(M', c', \alpha)$ preserve the objective function value and satisfy $T_{\text{total}}^{\text{'MLE}} = 1$. Therefore, we can first do the optimization over $\mathcal{D}_{\text{MLE}} = \{(M,c,\alpha)|T_{\text{total}}^{\text{MLE}} = 1\}$, yielding $(\widehat{M}_{\text{MLE}}, \widehat{c}_{\text{MLE}})$, and write the solution set of minimizing $l_{\text{MLE}} + \lambda_m\Psi$ as $\{(x\widehat{M}_{\text{MLE}}, x\widehat{c}_{\text{MLE}}, \alpha), x \in R^+\}$. The optimization problem over $\mathcal{D}_{\text{MLE}}$ can be interpreted as finding a representative element within the solution set. By applying the constraint

$\frac{o-1}{\mathrm{tr}[(-C\underline{T}C^T)^{-1}C\widehat{\Sigma}} = 1$ and negelecting the constants, this problem becomes minimizing $\frac{p}{2}(\log|-C\underline{T}C^T|)+\lambda_m\Psi$ over $\bar{D}_{\mathrm{MLE}}$.

On the other hand, the negative log likelihood after $T_{\mathrm{tot}}$ is absorbed can be written as

$$l = \frac{p}{2}\{-\mathrm{tr}[(C\underline{T}C^T)^{-1}C\widehat{\Sigma}C^T] + \log|-C\underline{T}C^T|\} \tag{S24}$$

The optimization problem is then minimizing $l + \lambda_m\Psi$ over $\mathcal{D}$. For any $(M, c, \alpha)$, the parameters can be rescaled as $(x_0M_0, x_0c_0, \alpha)$, where $x_0 \in R^+$ and $(M_0, c_0, \alpha) \in \mathcal{D}_{\mathrm{MLE}}$. The optimization problem is thus equivalent to first minimizing the function over $x_0$ given $(M_0, c_0, \alpha)$, and then minimizing over all possible possible parameters in $\mathcal{D}_{\mathrm{MLE}}$. By taking the derivative of the objective function with respect to $x_0$ and setting it to zero, we obtain $x_0 = 1$. The solution of minimizing $l + \lambda_m\Psi$ over $\mathcal{D}$ thus lies in $\mathcal{D}_{\mathrm{MLE}}$, and the problem is equivalent to minimize $l + \lambda_m\Psi$ over $D_{\mathrm{MLE}}$. By applying the constraint $\frac{o-1}{\mathrm{tr}[(-C\underline{T}C^T)^{-1}C\widehat{\Sigma}} = 1$ and negelecting the constants, the problem also reduces to minimizing $\frac{p}{2}(\log|-C\underline{T}C^T|) + \lambda_m\Psi$ over $\mathcal{D}_{\mathrm{MLE}}$. This is equivalent to selecting a representative of the solution set obtained after the MLE adjustment.

After these two simplifications, the final penalized likelihood function is $l + \lambda_m\Psi$ and the optimization problem becomes

$$\widehat{M}, \widehat{c}, \widehat{\alpha} = \underset{(M,c,\alpha)\in\mathcal{D}}{\mathrm{argmin}}\ (l + \lambda_m\Psi) \tag{S25}$$

#### 1.4 Computing of the expected pairwise coalescence time

Here we would like to compute $T|M,\gamma$ from equation (S2) . We develop an algorithm to solve the more general equation

$$\operatorname{diag}\{\gamma\}\operatorname{diag}\{X\} + LX + XL^T = G. \tag{S26}$$

with time complexity $O(d^4)$, where $G$ is an arbitrary symmetric matrix. We now introduce a few notations and give the outline of the method.

| Notation | Meaning |
|---|---|
| $\otimes$ | Kronecker Product |
| $\dagger$ | Moore-Penrose inverse |
| $\|\|\|\|_F$ | Frobenius Norm |
| vec | Vectorization |
| $(e_1,\ldots,e_d)$ | Standard basis of $R^d$ |
| $E_i, i=1,2,\ldots,d$ | $e_ie_i^T$ |
| $g$ | $\mathrm{vec}(G)$ |
| $\epsilon_i, i=1,2,\ldots,d$ | $\mathrm{vec}(E_i)$ |
| $S^d$ | The space of $d\times d$ symmetric matrix |
| $X_A^0, \forall A\in S^d$ | Least square solution of the equation: $LX+XL^T=A$ |
| $X_A^1, \forall A\in S^d$ | Solution of the equation $\gamma_1E_1\operatorname{diag}\{X\}+LX+XL^T=A$ |
| $X_A^d, \forall A\in S^d$ | Solution of the equation $\operatorname{diag}\{\gamma\}\operatorname{diag}\{X\}+LX+XL^T=A$ |
| $\chi_A^0, \forall A\in S^d$ | $\mathrm{vec}(X_A^0)$ |
| $\chi_A^1, \forall A\in S^d$ | $\mathrm{vec}(X_A^1)$ |
| $\chi_A^d, \forall A\in S^d$ | $\mathrm{vec}(X_A^d)$ |
| $\pi$ | Stationary distribution, a row vector |
| $\hat\pi$ | Normalized stationary distribution, i.e. $\frac{\pi}{\sqrt{\pi\pi^T}}$ |
| $\psi$ | $\pi\otimes\pi$ |
| $\hat\psi$ | $\hat\pi\otimes\hat\pi$ |
| $z$ | $1_{d\times1}$ |
| $Z$ | $zz^T$ |
| $\zeta$ | $z\otimes z$ |

**Table S1:** Notations

The remaining notations, as well as further explanations, will be provided in context. With the above notation, we proceed with the following steps:

1. Compute $X_G^0$ and $X_{E_i}^0$, for $i=1,2,\ldots,d$ ;

2. Compute $X_G^1$ and $X_{E_i}^1$, for $i=1,2,\ldots,d$ ;

3. Compute $X_G^d$

Before presenting the details of the three steps, we state two corollaries.

**Corollary 1**: The multiplicity of eigenvalue 0 is 1 for $L \otimes I + I \otimes L$, with the corresponding right eigenvector $\zeta$ and left eigenvector $\psi$.

Proof: In our cases, $L$ is the Laplacian of a strongly connected graph, thus the multiplicity for eigenvalue 0 is 1, and all other eigenvalues have positive real part (Caughman and Veerman, 2006). Noting that $L = -Q$, where $Q$ is the transition-rate matrix, we know $z = 1_{d\times 1}$ is a right eigenvector associated with eigenvalue 0.

By property of Kronecker product (Chapter 13, Laub, 2005), the eigenvalues of the matrix $L \otimes I + I \otimes L$ are of the form $\lambda_i + \lambda_j$, where $\lambda_i$ and $\lambda_j$ are eigenvalues of $L$. Since $L$ has eigenvalue 0 with multiplicity 1, and all other eigenvalues have positive real part, the matrix $L \otimes I + I \otimes L$ must also have eigenvalue 0 with multiplicity 1; the right-eigenvector corresponding to eigenvalue 0 is $z \otimes z = 1_{d^2\times 1} = \zeta$, and the corresponding left-eigenvector is $\pi \otimes \pi = \psi$.

**Corollary 2:** $\gamma_1 E_1 \operatorname{diag}(X) + LX + XL^T = A$ has a unique solution for any symmetric matrix $A$.

Proof: Rewriting the equation in vectorized form gives

$$(\gamma_1 \epsilon_1 \epsilon_1^T + L \otimes I + I \otimes L)\operatorname{vec}(X) = \operatorname{vec}(A). \tag{S27}$$

Thus, it suffices to show that the matrix $\gamma_1 \epsilon_1 \epsilon_1^T + L \otimes I + I \otimes L$ is non-singular. Since $L \otimes I + I \otimes L$ has rank $d^2 - 1$, we only need to verify that $\epsilon_1$ is not in the column space of $L \otimes I + I \otimes L$. If it were, it would be orthogonal to the left eigenvector $\pi \otimes \pi$. However, this is impossible because $\pi$, being the stationary distribution, has all positive entries.

##### Step 1

We aim to compute $X_G^0$ and $X_{E_i}^0$, $i = 1, 2, ..., d$ , it suffices to provide an algorithm that solves the equation

$$LX + XL^T - A = 0 \tag{S28}$$

for an arbitrary symmetric matrix $A$.

To solve this problem, we note that getting a least square solution of equation (S28) is equivalent to solving

$$LX + XL^T = A_{\parallel} \tag{S29}$$

exactly, where $A_{\parallel}$ is the projection of $A$ onto the range of the operator $L() + ()L^T$. This is equivalent to projecting $\operatorname{vec}(A)$ onto the range of $L \otimes I + I \otimes L$. Let $V$ denote the range of the operator $L \otimes I + I \otimes L$, and let $V^{\perp}$ be its orthogonal complement. The projection of $\operatorname{vec}(A)$ onto $V$ can then be written as $\operatorname{vec}(A)$ minus its projection onto $V^{\perp}$. As for $V^{\perp}$, it is simply the span of $\psi^T$, since

$$\begin{aligned}
& x \in V^{\perp} \\
\Leftrightarrow & \langle x, (L \otimes I + I \otimes L)y \rangle = 0, \forall y \in R^{d^2} \\
\Leftrightarrow & x^T (L \otimes I + I \otimes L)y = 0, \forall y \in R^{d^2} \\
\Leftrightarrow & x^T (L \otimes I + I \otimes L) = 0 \\
\Leftrightarrow & \exists k \in R, x^T = k\psi^T \quad \text{(Corollary 1)} \\
\Leftrightarrow & x \in \operatorname{Span}(\psi^T).
\end{aligned} \tag{S30}$$

Thus, we can write

$$\operatorname{vec}(A_{\parallel}) = \operatorname{vec}(A) - \hat{\psi}\operatorname{vec}(A)\hat{\psi}^T, \tag{S31}$$

or in the matrix form

$$A_{\parallel} = A - \hat{\psi}\operatorname{vec}(A)\hat{\pi}^T\hat{\pi}. \tag{S32}$$

Now, to solve the equation $LX + XL^T = A_{\parallel}$ exactly, we perform a transformation. Let $P_1 = \frac{z}{\sqrt{d}}$, $P = \begin{pmatrix} P_1 & P_2 \end{pmatrix}$ be a orthonormal matrix extended from vector $P_1$. Define

$$L' = P^T L P, \tag{S33a}$$

$$X' = P^T X P, \tag{S33b}$$

$$A'_{\parallel} = P^T A_{\parallel} P. \tag{S33c}$$

Partition these matrices as

$$L' = \begin{pmatrix} 0 & L'_{12} \\ 0 & L'_{22} \end{pmatrix}, \tag{S34a}$$

$$X' = \begin{pmatrix} X'_{11} & X'_{12} \\ X'_{21} & X'_{22} \end{pmatrix}, \tag{S34b}$$

$$A'_{\|} = \begin{pmatrix} (A'_{\|})_{11} & (A'_{\|})_{12} \\ (A'_{\|})_{21} & (A'_{\|})_{22} \end{pmatrix}, \tag{S34c}$$

where the $()_{11}$ is the entry in row 1, column 1. From block matrix multiplication, we derive

$$X'_{22}(L'_{12})^T + L'_{22}X'_{21} = (A'_{\|})_{21}, \tag{S35a}$$

$$L'_{22}X'_{22} + X'_{22}(L'_{22})^T = (A'_{\|})_{22}. \tag{S35b}$$

Since $L$ has eigenvalue 0 with multiplicity 1 and all other eigenvalues have positive real part, $-L$ is semi-stable. Consequently, $-L'_{22}$ is stable, ensuring that equation (S35b) has a unique solution. We solve (S35b) directly using Bartels–Stewart algorithm (Bartels and Stewart, 1972)), a standard method for solving Sylvester equations. $X'_{21}$ can then be obtained from (S35a) . By setting $X_{11} = 0$ and using the fact that $X'_{12} = X'^T_{21}$, we fully recover $X'$, the original solution is given by

$$X = PX'P^T. \tag{S36}$$

By choosing $A = G$ and $A = E_i$, $i = 1, 2, \ldots, d$ we can compute $X^0_G$ and $X^0_{E_i}$.

**Step 2**

In this step, we compute $X^1_G$ and $X^1_{E_i}$, for $i = 1, 2, \ldots, d$. Since $X^0_G$ and $X^0_{E_i}$ have already been computed, it suffices to show how to compute $X^1_A$ from $X^0_A$ and $X^0_{E_i}$, $i = 1, 2, \ldots, d$ for arbitrary matrix $A$.

Let $\alpha = \mathrm{vec}(A)$, From corollary 2, equation

$$(\gamma_1\epsilon_1\epsilon_1^T + L \otimes I + I \otimes L)\chi = \alpha \tag{S37}$$

.has a unique solution

$$\chi^1_A = (\gamma_1\epsilon_1\epsilon_1^T + L \otimes I + I \otimes L)^{-1}\alpha. \tag{S38}$$

Because the Moore-Penrose pseudoinverse coincides with the inverse in this case, we can also write

$$\chi^1_A = (\gamma_1\epsilon_1\epsilon_1^T + L \otimes I + I \otimes L)^{\dagger}\alpha. \tag{S39}$$

Using the rank-one update formula for the Moore-Penrose inverse developed by Carl D. Meyer (Meyer, 1973), we can efficiently compute $X^1_A$ . For any matrix $B$ and vectors $c_1$ and $c_2$ where $c_1$ is not in the column space of $B$ and $c_2$ is not in the row space of $B$ the update is

$$(B + c_1c_2^T)^{\dagger} = B^{\dagger} - k_1u^{\dagger} - v^{\dagger}k_2^T + (1 + c_2^TB^{\dagger}c_1)v^{\dagger}u^{\dagger}, \tag{S40}$$

where

$$k_1 = B^{\dagger}c_1, \tag{S41a}$$

$$k_2^T = c_2^TB^{\dagger}, \tag{S41b}$$

$$u = (I - BB^{\dagger})c_1, \tag{S41c}$$

$$v = c_2^T(I - B^{\dagger}B). \tag{S41d}$$

The expression can be transformed to

$$\begin{aligned} &B^{\dagger} - k_1u^{\dagger} - v^{\dagger}k_2^T + (1 + c_2^TB^{\dagger}c_1)v^{\dagger}u^{\dagger} \\ &= (B^{\dagger} - k_1u^{\dagger}) + v^{\dagger}(c_2^TB^{\dagger}c_1u^{\dagger} - k_2^T) + v^{\dagger}u^{\dagger} \\ &= B^{\dagger}(I - c_1u^{\dagger}) - v^{\dagger}c_2^TB^{\dagger}(I - c_1u^{\dagger}) + v^{\dagger}u^{\dagger} \\ &= (I - v^{\dagger}c_2^T)B^{\dagger}(I - c_1u^{\dagger}) + v^{\dagger}u^{\dagger} \end{aligned} \tag{S42}$$

Since $u^\dagger$ is a vector, we have $u^\dagger = \frac{u^T}{u^T u} = \frac{c_1^T(I-BB^\dagger)^T}{c_1^T(I-BB^\dagger)^T(I-BB^\dagger)c_1}$. From the definition of Moore-Penrose inverse, we obtain

$$(I - BB^\dagger)^T = I - BB^\dagger, \tag{S43a}$$

$$BB^\dagger(I - BB^\dagger) = 0. \tag{S43b}$$

This means

$$u^\dagger = \frac{c_1^T(I - BB^\dagger)}{c_1^T(I - BB^\dagger)c_1}. \tag{S44}$$

Similarly,

$$v^\dagger = \frac{(I - B^\dagger B)c_2}{c_2^T(I - B^\dagger B)c_2}. \tag{S45}$$

Now let $B = L \otimes I + I \otimes L$ , $c_1 = \gamma_1 \epsilon_1$ and $c_2 = \epsilon_1$. Since $\gamma\epsilon_1$ is not in the column space of $L \otimes I + I \otimes L$ and $\epsilon_1^T$ is not in the row space of $L \otimes I + I \otimes L$, the rank-one update formula applies. From the property of Moore-Penrose inverse, we know $B(I - B^\dagger B) = 0$, which implies $B(I - B^\dagger B)c_2 = 0$. From previous analysis, we know the kernel of $L \otimes I + I \otimes L$ is the space spanned by $\zeta = 1_{d^2\times 1} = z \otimes z$. Therefore $(I - B^\dagger B)c_2 = k\zeta$ for some $k$. Replace $c_2$ with $\epsilon_1$ and $(I - B^\dagger B)c_2$ with $k\zeta$ , we obtain

$$v^\dagger = \frac{\zeta}{\epsilon_1^T \zeta}. \tag{S46}$$

Similarly,

$$u^\dagger = \frac{\psi^T}{\gamma_1 \psi^T \epsilon_1}. \tag{S47}$$

Substituting $v^\dagger$ and $u^\dagger$ back into the rank-one update formula gives

$$\begin{aligned}&(\gamma_1\epsilon_1\epsilon_1^T + L \otimes I + I \otimes L)^\dagger \\ &= (I - \frac{\zeta\epsilon_1^T}{\epsilon_1^T\zeta})(L \otimes I + I \otimes L)^\dagger(I - \frac{\epsilon_1\psi}{\psi\epsilon_1}) + \frac{\zeta\psi}{\gamma_1(\psi\epsilon_1)(\epsilon_1^T\zeta)}.\end{aligned} \tag{S48}$$

We can then calculate the expression of $\chi_A^1$

$$\begin{aligned}\chi_A^1 =&(\gamma_1\epsilon_1\epsilon_1^T + L \otimes I + I \otimes L)^\dagger\alpha \\ =&(I - \frac{\zeta\epsilon_1^T}{\epsilon_1^T\zeta})[(L \otimes I + I \otimes L)^\dagger\alpha - (\frac{\psi\alpha}{\psi\epsilon_1})(L \otimes I + I \otimes L)^\dagger\epsilon_1] \\ &+ \frac{\psi\alpha}{\gamma_1(\psi\epsilon_1)(\epsilon_1^T\zeta)}\zeta.\end{aligned} \tag{S49}$$

Since $(I - \frac{\zeta\epsilon_1^T}{\epsilon_1^T\zeta})\zeta = 0$, we can use any representative in $\{(L\otimes I+I\otimes L)^\dagger\alpha+k\zeta\}$ to replace $(L\otimes I+I\otimes L)^\dagger\alpha$ in the expression. From the property of Moore-Penrose pseudoinverse (Barata and Hussein, 2012) and the fact that the space spanned by $\zeta$ is the kernel of $L \otimes I + I \otimes L$, we know that finding a representative in $\{(L \otimes I + I \otimes L)^\dagger\alpha + k\zeta\}$ is equivalent to finding a symmetric matrix $X$ such that $\mathrm{vec}(X)$ minimizes $\| (L \otimes I + I \otimes L)\mathrm{vec}(X) - \alpha \|$. This is further equivalent to finding a symmetric matrix $X$ that minimizes $\| LX + XL^T - A \|_F$, which is simply $X_A^0$ solved in step 1. This means that we can replace $(L\otimes I+I\otimes L)^\dagger\alpha$ with $\chi_A^0$ . $(L \otimes I + I \otimes L)^\dagger\epsilon_1$ can be replaced by $\chi_{E_1}^0$,with the same logic.

Using these substitutions, the expression for $\chi_A^1$ simplifies to

$$\chi_A^1 = (I - \frac{\zeta\epsilon_1^T}{\epsilon_1^T\zeta})[\chi_A^0 - (\frac{\psi\alpha}{\psi\epsilon_1})\chi_{E_1}^0] + \frac{\psi\alpha}{\gamma_1(\psi\epsilon_1)(\epsilon_1^T\zeta)}\zeta. \tag{S50}$$

In matrix form, this becomes

$$X_A^1 = X_A^0 - \frac{\psi\alpha}{\pi_1^2}X_{E_1}^0 + k_0 Z, \tag{S51}$$

where $k_0 = \frac{\psi^T\alpha}{\gamma_1\pi_1^2} - X_A^0(1,1) + \frac{\psi^T\alpha}{\pi_1^2}X_{E_1}^0(1,1)$, $\pi_i$ represents the $i_{\text{th}}$ entry of $y$ and $X_A^0(1,1)$ represents the element in row 1 column 1 of $X_A^0$.

Notice that we can always adjust $X_A^0$ in step 1 such that $X_A^0(1,1) = 0$. This can be done by substracting $X_A^0(1,1)Z$ from $X_A^0$. With this adjustment, the expression for $X_A^1$ simplifies to

$$X_A^1 = X_A^0 - \frac{\psi\alpha}{\pi_1^2}X_{E_1}^0 + \frac{\psi\alpha}{\gamma_1\pi_1^2}Z. \tag{S52}$$

Now by letting $A = G$ and $A = E_i$, $i = 1, 2, \ldots, d$, we obtain

$$X_G^1 = X_G^0 - \frac{\psi g}{\pi_1^2}X_1^0 + \frac{\psi g}{\gamma_1\pi_1^2}Z, \tag{S53a}$$

$$X_{E_i}^1 = X_{E_i}^0 - \frac{\pi_i^2}{\pi_1^2}X_1^0 + \frac{\pi_i^2}{\gamma_1\pi_1^2}Z. \quad i = 1, 2, \ldots, d. \tag{S53b}$$

**Step 3**

In this step, we compute the final solution $X_G^d$ .

Define the matrices

$$H_1 = \gamma_1\epsilon_1\epsilon_1^T + L\otimes I + I\otimes L, \tag{S54a}$$

$$H_d = \sum_{i=1}^{d}\gamma_i\epsilon_i\epsilon_i^T + L\otimes I + I\otimes L. \tag{S54b}$$

Since $H_1$ is non-singular, we can apply the Sherman-Morrison-Woodbury formula to effeciently compute $H_d$. Let $U \in R^{d^2\times d-1}$ be a matrix where its $i_{th}$ column is $\epsilon_{i+1}$, and $U_\gamma \in R^{d^2\times d-1}$be a matrix where its $i_{th}$ column is $\gamma_i\epsilon_{i+1}$, then

$$H_d = H_1 + UU_\gamma^T. \tag{S55}$$

Applying the Sherman-Morrison-Woodbury formula, we obtain

$$H_d^{-1} = H_1^{-1} - H_1^{-1}U(I + U_\gamma^T H_1^{-1}U)^{-1}U_\gamma^T H_1^{-1}. \tag{S56}$$

We assume $I + U_\gamma^T H_1^{-1}U$ is invertible, which is valid for almost all parameter values.

Now, we compute $X_G^d$

$$\begin{aligned}\chi_G^d &= H_d^{-1}g \\ &= H_1^{-1}g - H_1^{-1}U(I + U_\gamma^T H_1^{-1}U)^{-1}U_\gamma^T H_1^{-1}g \\ &= \chi_G^1 - H_1^{-1}U(I + U_\gamma^T H_1^{-1}U)^{-1}U_\gamma^T\chi_G^1.\end{aligned} \tag{S57}$$

$(I + U_\gamma^T H_1^{-1}U)^{-1}U_\gamma^T\chi_G^1$ can computed by solving the equation

$$(I + U_\gamma^T H_1^{-1}U)x = U_\gamma^T\chi_G^1, \tag{S58}$$

where $U_\gamma^T\chi_G^1$ is a $d-1\times 1$ vector whose $i_{th}$ entry is $\gamma_{i+1}X_G^1(i+1, i+1)$.

This equation can be written explicitly as

$$\begin{pmatrix} X_{E_2}^1(2,2) + \frac{1}{\gamma_2} & \ldots & X_{E_d}^1(2,2) \\ \vdots & \ldots & \vdots \\ X_{E_2}^1(d,d) & \ldots & X_{E_d}^1(d,d) + \frac{1}{\gamma_d}\end{pmatrix} x = \begin{pmatrix} X_G^1(2,2) \\ \vdots \\ X_G^1(d,d)\end{pmatrix}. \tag{S59}$$

After we get the solution we then compute $Ux$. Since $i_{\text{th}}$ column of $U$ is $\epsilon_{i+1}$, this simplifies to compute

$$Ux = \sum_{i=2}^{d} x_{i-1}\epsilon_i, \tag{S60}$$

where $x_i$ represents the $i_{\text{th}}$ entry of $x$. This means

$$H_1^{-1}Ux = \sum_{i=2}^{d} x_{i-1}(H_1^{-1}\epsilon_i). \tag{S61}$$

Or in matrix form

$$X_G^d = X_G^1 - \sum_{i=2}^{d} x_{i-1}X_{E_i}^1. \tag{S62}$$

**A direct method**

The above method is based on rank one update of Moore-Penrose inverse and Sherman-Morrison-Woodbury formula, here we also give a direct way to find the solution. The first step is the same as the above method, i.e. computing $X_G^0$ and $X_{E_i}^0$, for $i = 1, 2, ..., d$ . The solutions satisfy

$$LX_G^0 + X_G^0 L^T = G - \hat{\psi} g \hat{\pi}^T \hat{\pi}, \tag{S63a}$$
$$LX_{E_i}^0 + X_{E_i}^0 L^T = E_i - \hat{\pi}_i^2 \hat{\pi}^T \hat{\pi}, \quad i = 1, 2, \ldots, d. \tag{S63b}$$

The final solution is assumed to be a linear combination of $X_G^0$, $X_{E_i}^0$ and $Z$

$$X_G^d = X_G^0 - \sum_{i=1}^{d} a_i X_{E_i}^0 - bZ \tag{S64}$$

The first equation we get is the constraint that the projection cancels out

$$\hat{\psi} g - \sum_{i=1}^{d} a_i \hat{\pi}_i^2 = 0. \tag{S65}$$

Apply the operator $L() + ()L^T$ to $X_G^d$ we get

$$LX_G^d + X_G^d L^T = G - \sum_{i=1}^{d} a_i E_i. \tag{S66}$$

Combining with equation (S26) , from the diagonal elements we obtain another $d$ equations

$$X_G^0(j,j) - \sum_{i=1}^{d} a_i X_{E_i}^0(j,j) - b = \frac{a_j}{\gamma_j}, \quad i = 1, 2, \ldots, d. \tag{S67}$$

The $d+1$ linear equations of the $d+1$ unknowns $\{b, a_i, 1 \le i \le d\}$ can then be written in matrix notation

$$\begin{pmatrix} 0 & \hat{\pi}_1^2 & \ldots & \hat{\pi}_d^2 \\ 1 & X_{E_1}^0(1,1) + \frac{1}{\gamma_1} & \ldots & X_{E_d}^0(1,1) \\ \vdots & \vdots & \ldots & \vdots \\ 1 & X_{E_1}^0(d,d) & \ldots & X_{E_d}^0(d,d) + \frac{1}{\gamma_d} \end{pmatrix} \begin{pmatrix} b \\ a_1 \\ \vdots \\ a_d \end{pmatrix} = \begin{pmatrix} \hat{\psi} g \\ X_G^0(1,1) \\ \vdots \\ X_G^0(d,d) \end{pmatrix}. \tag{S68}$$

The solution is obtained by solving the above linear system, which provides values for $b$ and $a_i$, $i = 1, 2, \ldots, d$. These coefficients then fully determine $X_G^d$.

**Summary**

The total time complexity of both algorithms is $O(d^4)$, primarily due to solving $O(d)$ Lyapunov equations in step 1. Similarly, the total space complexity is $O(d^3)$, as it is dominated by storing $O(d)$ matrices during the same step. In practice, the first method is preferred for its robustness. By selecting the deme with the largest $\pi_i$ and performing a rank-1 update on it first, we effectively control the numerical computation's size, ensuring stability and efficiency. The second method, while easier to understand, can serve as an intuitive illustration.

#### 1.5 Computing of the gradient

The gradient of the object function is the sum of the gradient of $l$, and $\lambda_m \Psi_{\lambda_m}$. In this section we show how to compute the gradient of the three components.

**Computating $\nabla l$**

We start by showing how to compute $\frac{\partial l}{\partial m_{st}}$ for a given $m_{st}$ where $(s,t) \in \Omega$. Let $T$ be the solution of equation (S2) and let $\tau = \text{vec}(T)$. Using the chain rule, we have

$$\frac{\partial l}{\partial m_{st}} = \frac{\partial l}{\partial \tau}\frac{\partial \tau}{\partial m_{st}}, \tag{S69}$$

where $\frac{\partial l}{\partial \tau}$ is a $1 \times d^2$ row vector and $\frac{\partial \tau}{\partial L_{st}}$ is a $d^2 \times 1$ column vector.

We first compute $\frac{\partial l}{\partial \tau}$. From the expression

$$l = \frac{p}{2}\{-\operatorname{tr}[(C\underline{T}C^T)^{-1}C\widehat{\Sigma}C^T] + \log|-C\underline{T}C^T|\}, \tag{S70}$$

we know

$$\frac{\partial l}{\partial C\underline{T}C^T} = \frac{p}{2}[(C\underline{T}C^T)^{-1}C\widehat{\Sigma}C^T(C\underline{T}C^T)^{-1} + (C\underline{T}C^T)^{-1}]. \tag{S71}$$

From this and the fact that

$$\frac{\partial l}{\partial \underline{T}} = C^T\frac{\partial l}{\partial C\underline{T}C^T}C, \tag{S72}$$

we can compute $\frac{\partial l}{\partial T}$ using

$$\frac{\partial l}{\partial T} = J^T\left[\frac{\partial l}{\partial \underline{T}} - \operatorname{diag}\left\{\frac{\partial l}{\partial \underline{T}}\right\}\operatorname{diag}\{K\}\right]J, \tag{S73}$$

(this is because $\underline{T} = JTJ^T - \operatorname{diag}\{JTJ^T\}\operatorname{diag}\{K\}$). $\frac{\partial l}{\partial \tau}$ can then be computed from

$$\frac{\partial l}{\partial \tau} = [\text{vec}(\frac{\partial l}{\partial T})]^T. \tag{S74}$$

Next, we compute $\frac{\partial l}{\partial \tau}\frac{\partial \tau}{\partial m_{st}}, (s,t) \in \Omega$ given $\frac{\partial l}{\partial \tau}$. From the last section, we know $H_d\tau = \zeta$. This implies

$$\frac{\partial H_d}{\partial m_{st}}\tau + H_d\frac{\partial \tau}{\partial m_{st}} = 0. \tag{S75}$$

Thus,

$$H_d\frac{\partial \tau}{\partial m_{st}} = -\frac{\partial H_d}{\partial m_{st}}\tau, \tag{S76}$$

This means

$$\frac{\partial \tau}{\partial m_{st}} = -H_d^{-1}\frac{\partial H_d}{\partial m_{st}}\tau, \tag{S77}$$

and

$$\frac{\partial l}{\partial \tau}\frac{\partial \tau}{\partial m_{st}} = -(\frac{\partial l}{\partial \tau}H_d^{-1})\frac{\partial H_d}{\partial m_{st}}\tau. \tag{S78}$$

Notice that

$$\frac{\partial H_d}{\partial m_{st}} = \frac{\partial H_d}{\partial L_{ss}} - \frac{\partial H_d}{\partial L_{st}} + \sum_{i=1}^{d}\frac{\partial H_d}{\partial \gamma_i}\frac{\partial \gamma_i}{\partial m_{st}}, \tag{S79}$$

we have

$$-(\frac{\partial l}{\partial \tau}H_d^{-1})\frac{\partial H_d}{\partial m_{st}}\tau = (\frac{\partial l}{\partial \tau}H_d^{-1})(\frac{\partial H_d}{\partial L_{st}} - \frac{\partial H_d}{\partial L_{ss}})\tau - \sum_{i=1}^{d}\frac{\partial \gamma_i}{\partial m_{st}}(\frac{\partial l}{\partial \tau}H_d^{-1})\frac{\partial H_d}{\partial \gamma_i}\tau. \tag{S80}$$

The computation of the right-hand side can be divided into three parts, i.e. computing $\frac{\partial l}{\partial \tau} H_d^{-1}$, computing $(\frac{\partial H_d}{\partial L_{ss}} - \frac{\partial H_d}{\partial L_{st}})\tau$ as well as $\frac{\partial H_d}{\partial \gamma_i}\tau$, and computing $\frac{\partial \gamma_i}{\partial m_{st}}$.

For the second part, the expressions

$$(\frac{\partial H_d}{\partial L_{st}} - \frac{\partial H_d}{\partial L_{ss}})\tau = \mathrm{vec}(e_s e_t^T T + T e_t e_s^T) - \mathrm{vec}(e_s e_s^T T + T e_s e_s^T) \tag{S81}$$

and

$$\frac{\partial H_d}{\partial \gamma_i}\tau = \epsilon_i \epsilon_i^T \tau = (\epsilon_i^T \tau)\epsilon_i \tag{S82}$$

are easy to calculate as long as we get $T$.

For the third part, since $\gamma_i = \frac{c}{\pi_i^\alpha}$, we know

$$\frac{\partial \gamma_i}{\partial m_{st}} = \frac{-c}{\pi_i^{\alpha+1}} \frac{\partial \pi_i}{\partial m_{st}} = \frac{-\gamma_i}{\pi_i} \frac{\partial \pi_i}{\partial m_{st}}. \tag{S83}$$

On the other hand,

$$\frac{\partial \pi}{\partial m_{st}} = \frac{\partial \pi}{\partial L_{ss}} - \frac{\partial \pi}{\partial L_{st}}. \tag{S84}$$

Since $\pi L = 0$ and $\pi z = 1$, we have

$$\pi \frac{\partial L}{\partial L_{st}} + \frac{\partial \pi}{\partial L_{st}} L = 0, \tag{S85}$$

and

$$\frac{\partial \pi}{\partial L_{st}} z = 0. \tag{S86}$$

This means

$$\frac{\partial \pi}{\partial L_{st}}(L + Z) = -\pi \frac{\partial L}{\partial L_{st}}, \tag{S87}$$

which implies

$$\frac{\partial \pi}{\partial L_{st}} = -\pi \frac{\partial L}{\partial L_{st}}(L + Z)^{-1}, \tag{S88}$$

and

$$\frac{\partial \pi}{\partial m_{st}} = -\pi(\frac{\partial L}{\partial L_{ss}} - \frac{\partial L}{\partial L_{st}})(L + Z)^{-1}. \tag{S89}$$

We then obtain

$$\frac{\partial \pi_i}{\partial m_{st}} = -\pi(\frac{\partial L}{\partial L_{ss}} - \frac{\partial L}{\partial L_{st}})(L + Z)^{-1} e_i. \tag{S90}$$

Now we only need to compute the first part

$$\frac{\partial l}{\partial \tau} H_d^{-1} = [(H_d^T)^{-1}(\frac{\partial l}{\partial \tau})^T]^T, \tag{S91}$$

which further reduces to the computation of

$$(H_d^T)^{-1}(\frac{\partial l}{\partial \tau})^T. \tag{S92}$$

However, we know

$$H_d^T = \sum_{i=1}^{d} \gamma_i \epsilon_i \epsilon_i^T + L^T \otimes I + I \otimes L^T, \tag{S93}$$

thus we only need to solve the equation

$$\mathrm{diag}\{\gamma\}\,\mathrm{diag}\{X\} + L^T X + X L = \frac{\partial l}{\partial T}. \tag{S94}$$

Let $T^{\nabla}$ be the solution to the equation. Then $(\frac{\partial l}{\partial \tau} H_d^{-1})\frac{\partial H_d}{\partial L_{st}}\tau$ is the summation of entries of $T^{\nabla} \odot (e_s e_t^T T + T e_t e_s^T)$, where $\odot$ represents the Hadamard product. This is simply twice the inner product of the $s_{th}$ column of $T^{\nabla}$ and the $t_{th}$ column of $T$. The expression $(\frac{\partial l}{\partial \tau} H_d^{-1})(\frac{\partial H_d}{\partial L_{st}} - \frac{\partial H_d}{\partial L_{ss}})\tau$ can be computed. Similarly, since

$$\frac{\partial H_d}{\partial \gamma_i}\tau = (\epsilon_i^T \tau)\epsilon_i, \tag{S95}$$

we have

$$(\frac{\partial l}{\partial \tau} H_d^{-1})\frac{\partial H_d}{\partial \gamma_i}\tau = T_{ii}^{\nabla} T_{ii}, \tag{S96}$$

where $T_{ii}^{\nabla}$ and $T_{ii}$ represents the $(i,i)_{\text{th}}$ entry of $T^{\nabla}$ and $T$. Combining this equation with equation (S83), we have

$$-\sum_{i=1}^{d} \frac{\partial \gamma_i}{\partial m_{st}} (\frac{\partial l}{\partial \tau} H_d^{-1})\frac{\partial H_d}{\partial \gamma_i}\tau = \pi(\frac{\partial L}{\partial L_{st}} - \frac{\partial L}{\partial L_{ss}})(L+Z)^{-1}[\sum_{i=1}^{d} \frac{\gamma_i}{\pi_i} T_{ii}^{\nabla} T_{ii} e_i] \tag{S97}$$

Let $x$ be a vector such that

$$(L+Z)x = \delta, \tag{S98}$$

where $\delta = \sum_{i=1}^{d} \frac{\gamma_i}{\pi_i} T_{ii}^{\nabla} T_{ii} e_i$ is a vector such that its $i_{\text{th}}$ element is $\frac{\gamma_i}{\pi_i} X_{ii}^{\nabla} T_{ii}$. The expression $-\sum_{i=1}^{d} \frac{\partial \gamma_i}{\partial m_{st}} (\frac{\partial l}{\partial \tau} H_d^{-1})\frac{\partial H_d}{\partial \gamma_i}\tau$ can then be reduced to $\pi_s(x_t - x_s)$, where $x_t$ represents the $t_{\text{th}}$ entry of $x$.

Now we only need to solve the equation (S94). The method is similar to that shown in section 1.4. The only difference is that for $L^T$ the right-eigenvector corresponding to eigenvalue 0 is $\pi^T$, and the corresponding left eigenvector is $z^T$. After we correct for this difference, the procedures are exactly the same. We can compute this with time complexity $O(d^4)$ and space complexity $O(d^3)$. To be more precise, we go through the following procedure to compute the solution of the equation

$$\operatorname{diag}\{\gamma\} \operatorname{diag}\{X\} + L^T X + XL = G. \tag{S99}$$

**1.**Compute the exact least square solutions of

$$L^T X + XL = G, \tag{S100a}$$

$$L^T X + XL = E_i, \quad i = 1, 2, \ldots, d. \tag{S100b}$$

It suffices to solve

$$L^T X + XL = A \tag{S101}$$

for arbitrary symmetric matrix $A$ in the least square sense.

To so this, we project $A$ onto the range of $L^T() + ()L$. Similar to the previous analysis, we see that the projection is

$$A_{\parallel} = A - \frac{1}{d^2}\zeta^T \operatorname{vec}(A) Z. \tag{S102}$$

We then perform the same transformation, i.e.

$$(L^T)' = P^T L^T P = (L')^T, \tag{S103a}$$

$$X' = P^T X P, \tag{S103b}$$

$$A'_{\parallel} = P^T A_{\parallel} P, \tag{S103c}$$

and use the following block division

$$L' = \begin{pmatrix} 0 & 0 \\ (L'_{12})^T & (L'_{22})^T \end{pmatrix}, \tag{S104a}$$

$$X' = \begin{pmatrix} X'_{11} & X'_{12} \\ X'_{21} & X'_{22} \end{pmatrix}, \tag{S104b}$$

$$A'_{\parallel} = \begin{pmatrix} (A'_{\parallel})_{11} & (A'_{\parallel})_{12} \\ (A'_{\parallel})_{21} & (A'_{\parallel})_{22} \end{pmatrix}. \tag{S104c}$$

The equations to be solved are then

$$(L'_{12})^T X'_{11} + (L'_{22})^T X'_{21} = (A'_{\|})_{21}, \tag{S105a}$$

$$L'_{22} X'_{22} + X'_{22} (L'_{22})^T = (A'_{\|})_{22} - (L'_{12})^T (X'_{21})^T - X'_{21} L'_{12}. \tag{S105b}$$

Setting $X_{11} = 0$, we can then get $X'_{21}$ from the first equation. The second equation is a canonical Lyapunov equation, and we can solve $X'_{22}$ from it. Using the fact that $X'_{12} = X'^T_{21}$, we get $X'$, and the solution $X$ can then be obtained from $PX'P^T$. By setting $A = G$ and $A = E_i$, $i = 1, 2, \ldots, d$, we obtain the solutions.

**2.** Perform a rank-1 update similar to step 2 in section 1.4, switching between $L$ and $L^T$, as well as between $\zeta$ and $\psi^T$ in $v^\dagger$ and $u^\dagger$. Define $X^0_A$ as the least-squares solution of $L^T X + XL = A$, and $\chi^0_A$ as its vectorization. Consistent with the conventions established in section 1.4 (except for replacing $L$ with $L^T$), introduce $X^1_A, \chi^1_A, X^d_A, \chi^d_A$. We obtain

$$\chi^1_G = \left(I - \frac{\psi^T \epsilon_1^T}{\psi \epsilon_1}\right) \left[\chi^0_G - \left(\frac{\zeta^T g}{\zeta^T \epsilon_1}\right) \chi^0_{E_1}\right] + \frac{\zeta^T g}{\gamma_1 (\zeta^T \epsilon_1)(\psi \epsilon_1)} \psi^T, \tag{S106a}$$

$$\chi^1_{E_i} = \left(I - \frac{\psi^T \epsilon_1^T}{\psi \epsilon_1}\right) \left[\chi^0_{E_i} - \left(\frac{\zeta^T \epsilon_i}{\zeta^T \epsilon_1}\right) \chi^0_{E_1}\right] + \frac{\zeta^T \epsilon_i}{\gamma_1 (\zeta^T \epsilon_1)(\psi \epsilon_1)} \psi^T, \quad i = 1, 2, \ldots, d. \tag{S106b}$$

By adjusting $X^0_{E_i}$ such that $X^0_{E_i}(1,1) = 0$ (through subtracting $\frac{X^0_{E_i}(1,1)}{\pi_1^2} \psi$ from $X^0_{E_i}$), the equation in matrix form can then be written as

$$X^1_G = X^0_G - (\zeta^T \epsilon_0) X^0_{E_1} + \frac{\zeta^T \epsilon_0}{\gamma_1 \pi_1^2} \pi^T \pi, \tag{S107a}$$

$$X^1_{E_i} = X^0_{E_i} - X^0_{E_1} + \frac{1}{\gamma_1 \pi_1^2} \pi^T \pi, \quad i = 1, 2, \ldots, d. \tag{S107b}$$

**3.** Use the Sherman-Morrison-Woodbury formula similar to step 3 in the last section, solving

$$\left[I + U_\gamma^T (H_1^{-1})^T U\right] x = U_\gamma^T \chi^1_G \tag{S108}$$

and obtaining the solution through

$$X^d_G = X^1_G - \sum_{i=2}^{d} x_{i-1} X^1_{E_i}. \tag{S109}$$

By setting $G = \frac{\partial l}{\partial T}$, we can compute the solution $T^\nabla$.

For the gradient, we also need to compute $\frac{\partial l}{\partial c}$ and $\frac{\partial l}{\partial \alpha}$. Similar to previous calculations, we know

$$\frac{\partial l}{\partial \gamma_i} = -\left(\frac{\partial l}{\partial \tau} H_d^{-1}\right) \frac{\partial H_d}{\partial \gamma_i} \tau = -T^\nabla_{ii} T_{ii}. \tag{S110}$$

Thus

$$\frac{\partial l}{\partial c} = \sum_{i=1}^{d} \frac{\partial l}{\partial \gamma_i} \frac{\partial \gamma_i}{\partial c} = \sum_{i=1}^{d} \frac{\partial l}{\partial \gamma_i} \frac{1}{\pi_i^\alpha} = -\sum_{i=1}^{d} \frac{1}{\pi_i^\alpha} T^\nabla_{ii} T_{ii}, \tag{S111a}$$

$$\frac{\partial l}{\partial \alpha} = \sum_{i=1}^{d} \frac{\partial l}{\partial \gamma_i} \frac{\partial \gamma_i}{\partial \alpha} = -\sum_{i=1}^{d} \frac{\partial l}{\partial \gamma_i} \frac{c}{\pi_i^\alpha} \log(\alpha) = \sum_{i=1}^{d} \frac{c \log(\alpha)}{\pi_i^\alpha} T^\nabla_{ii} T_{ii}. \tag{S111b}$$

The gradients of $l$ are then fully determined.

**Computing $\nabla\Psi$**

The penalty function is

$$\Psi = \frac{1}{2}\sum_{i=1}^{d} \sum_{\substack{(i_1,j_1)\in\Omega_i,\\(i_2,j_2)\in\Omega_i}} \frac{1}{|\Omega_i|^2}\left[\left(\frac{m_{i_1j_1}}{\widetilde{m}} - \frac{m_{i_2j_2}}{\widetilde{m}}\right)^2 + \left(\frac{\widetilde{m}}{m_{i_1j_1}} - \frac{\widetilde{m}}{m_{i_2j_2}}\right)^2\right]. \tag{S112}$$

To compute its gradient, we first consider the gradient of the first term $\frac{1}{2}\sum\limits_{i=1}^{d} \sum\limits_{\substack{(i_1,j_1)\in\Omega_i,\\(i_2,j_2)\in\Omega_i}} \frac{1}{|\Omega_i|^2}\left(\frac{m_{i_1j_1}}{\widetilde{m}} - \frac{m_{i_2j_2}}{\widetilde{m}}\right)^2$.

To simplify the notation, we let $g_{st} = \frac{m_{st}}{\widetilde{m}}$. The first term then becomes $\frac{1}{2}\sum\limits_{i=1}^{d} \sum\limits_{\substack{(i_1,j_1)\in\Omega_i,\\(i_2,j_2)\in\Omega_i}} \frac{1}{|\Omega_i|^2}(g_{i_1j_1} - g_{i_2j_2})^2$.

Using the identity

$$\frac{1}{2}\sum_{\substack{(i_1,j_1)\in\Omega_i,\\(i_2,j_2)\in\Omega_i}} (g_{i_1j_1} - g_{i_2j_2})^2 = |\Omega_i|\left(\sum_{(s,t)\in\Omega_i} g_{st}^2\right) - \left(\sum_{(s,t)\in\Omega_i} g_{st}\right)^2, \tag{S113}$$

we find

$$\begin{aligned}&\frac{1}{2}\sum_{i=1}^{d} \sum_{\substack{(i_1,j_1)\in\Omega_i,\\(i_2,j_2)\in\Omega_i}} \frac{1}{|\Omega_i|^2}(g_{i_1j_1} - g_{i_2j_2})^2 \\ &= \sum_{i=1}^{d}\left[\frac{1}{|\Omega_i|}\left(\sum_{(s,t)\in\Omega_i} g_{st}^2\right) - \frac{1}{|\Omega_i|^2}\left(\sum_{(s,t)\in\Omega_i} g_{st}\right)^2\right].\end{aligned} \tag{S114}$$

This expression, denoted as $\Psi^g$, represents the penalty on the variance of edge weights within nodes. The gradient with respect to $m_{ij}$ is then given by

$$\frac{\partial \Psi^g}{\partial m_{ij}} = \sum_{(s,t)\in\Omega} \frac{\partial \Psi^g}{\partial g_{st}}\frac{\partial g_{st}}{\partial m_{ij}}, \tag{S115}$$

where

$$\frac{\partial \Psi^g}{\partial g_{st}} = 2\left[\left(\frac{1}{|\Omega_s|} + \frac{1}{|\Omega_t|}\right) g_{st} - \frac{1}{|\Omega_s|}\sum_{(i',j')\in\Omega_s} g_{i'j'} - \frac{1}{|\Omega_t|}\sum_{(i',j')\in\Omega_t} g_{i'j'}\right]. \tag{S116}$$

For $\frac{\partial g_{st}}{\partial m_{ij}}$, we use the equation

$$\log(\widetilde{m}) = \frac{1}{|\Omega|}\sum_{(i',j')\in\Omega} \log(m_{i'j'}) \tag{S117}$$

and obtain

$$\frac{\partial g_{st}}{\partial m_{ij}} = \frac{1}{\widetilde{m}}\left(I\{(i,j) = (s,t)\} - \frac{1}{|\Omega|}\frac{m_{st}}{m_{ij}}\right), \tag{S118}$$

where $I\{(i,j) = (s,t)\}$ is the indicator function. Similarly, let $h_{st} = \frac{\widetilde{m}}{m_{st}}$ and denote the second term by $\Psi^h$, we have

$$\frac{\partial \Psi^h}{\partial m_{ij}} = \sum_{(s,t)\in\Omega} \frac{\partial \Psi^h}{\partial h_{st}}\frac{\partial h_{st}}{\partial m_{ij}}, \tag{S119}$$

where

$$\frac{\partial \Psi^h}{\partial h_{st}} = 2\left[\left(\frac{1}{|\Omega_s|} + \frac{1}{|\Omega_t|}\right) h_{st} - \frac{1}{|\Omega_s|}\sum_{(i',j')\in\Omega_s} h_{i'j'} - \frac{1}{|\Omega_t|}\sum_{(i',j')\in\Omega_t} h_{i'j'}\right]. \tag{S120}$$

For $\frac{\partial h_{st}}{\partial m_{ij}}$, we obtain

$$\frac{\partial h_{st}}{\partial m_{ij}} = \frac{-h_{st}^2}{\widetilde{m}} \left( I\{(i,j) = (s,t)\} - \frac{1}{|\Omega|} \frac{m_{st}}{m_{ij}} \right). \tag{S121}$$

The total gradient $\frac{\partial \Psi}{\partial m_{ij}}$ is then given by

$$\frac{\partial \Psi}{\partial m_{ij}} = \frac{\partial \Psi^g}{\partial m_{ij}} + \frac{\partial \Psi^h}{\partial m_{ij}}. \tag{S122}$$

### 2 Settings of simulations and empirical datasets

#### 2.1 Non-equilibrium simulations

The non-equilibrium stepping stone simulations are performed using `msprime` (Baumdicker et al., 2022). For the pulse migration scenario, we assume a uniform base migration rate across the graph. Forward in time, the migration process takes place at discrete time points $4i\Delta t_m$, where $i = 1, 2, \ldots$, and $\Delta t_m$ represents the time interval between mass migration events. At each of these time points, the frontier populations of the migration wave—beginning with the populations in the central column and progressing outward until reaching the boundary columns—split and send individuals to adjacent demes. During each mass migration event, let population $A$ denote the source population and population $B$ the recipient population. After the migration, 50% of the individuals in population $B$ will originate from population $A$.

Backward in time, this scenario represents populations migrating from the boundary columns towards the central column. In each mass migration event, 50% of the individuals in population $B$ will migrate backward to population $A$, effectively reversing the direction of movement observed in forward time.

It's important to mention that the uniform base migration happens parallel to mass migration events, and will effectively dilute the effect of the mass migration events. This is different from the case of range expansion, as we shall see.

Forward in time, the range expansion process occurs at discrete time points $4i\Delta t_r$, where $n = 1, 2, \ldots$, and $\Delta t_r$ represents the time interval between range expansion events. At each of these time points, the frontier populations of the migration wave—starting from the populations in the central column and spreading outward until reaching the boundary columns—split and send 10% of their individuals to occupy the adjacent vacant demes. Initially, all demes, except for those in the central column, are vacant. The newly occupied demes then undergo a period of exponential growth from $4i\Delta t_r$ to $(4i+1)\Delta t_r$, during which their population size reaches that of their ancestral population. These demes remain at this size until the next range expansion event at $4(i+1)\Delta t_r$.

Backward in time, this process corresponds to the newly occupied demes undergoing exponential shrinkage and merging with their ancestral deme, such that the merged population consists of 10% lineages from the newly occupied deme and 90% lineages from the ancestral deme.

Parameters of our range expansion model are adapted from the human Out-of-Africa model (Ramachandran et al., 2005), where $\Delta t_r$ is very small due to the rapid expansion. However, because our model includes only a small number of demes, the total duration is short—approximately $20\Delta t_r$—and the effect of base migration would be minor, even when included. This differs from the original Out-of-Africa model, which involves over 200 demes, making careful model comparisons that account for base migration necessary.

#### 2.2 Forward in time simulations

The forward-in-time individual-based simulations shown in Fig. 4 and Supplementary Fig. 11 are performed using `SLiM` (Haller et al., 2025). The simulation settings are adapted from Lundgren and Ralph (2019). In each migration scenario, simulations are conducted on an $8 \times 8$ continuous square landscape. Each diploid individual carries one pair of chromosomes of length $10^8$ base pairs and has a mean lifetime of $l = 4$ generations.

For each spatial position $(x, y)$, we define a local carrying capacity $K(x, y)$. For each individual in each generation, local density dependence is modeled using a gaussian competition kernel with standard deviation $\sigma = 0.1$, truncated at $s = 3\sigma = 0.3$. Individual fitness is defined using a logistic form, $w = \frac{2}{1+\frac{l+1}{l-1}\frac{a}{2\pi s^2 K(x,y)}}$, where $a$ denotes the number of individuals within the competition kernel. When $K(x, y)$ is spatially constant with value $K$ and $a = 2\pi s^2 K$, the fitness simplifies to $w = \frac{2}{1+\frac{l+1}{l-1}} = \frac{l-1}{l}$. If the survival probability equals $\frac{l-1}{l}$, the expected lifetime of an individual is $l$, ensuring demographic consistency.

To convert fitness into a survival probability, we apply two additional steps. First, individuals close to the boundary receive a boundary penalty defined as $p_b = \min(1, 5x) \min(1, 5y) \min(1, 40 - 5x) \min(1, 40 - 5y)$, which reduces survival for individuals located within 0.2 units of the landscape boundary. Second, to prevent excessive spatial clustering, following Felsenstein (1975), the survival probability is capped as $p_s = \min(0.9, wp_b)$.

In each generation, after the survivorship step, mating and reproduction occur. Each surviving individual mates with another surviving individual according to a gaussian mating kernel with standard deviation $\sigma = 0.1$, truncated at $s = 3\sigma = 0.3$. After mating, the number of offspring is drawn from a Poisson distribution with mean $\frac{1}{l} = 0.25$, so that on average each individual produces one offspring over its lifetime.

For each offspring, the recombination rate is set to $10^{-8}$ per base pair per generation. Offspring dispersal occurs once at birth: if individual $A$ mates with individual $B$ to produce offspring $C$, then $C$ disperses from the spatial position of $A$ according to a Gaussian dispersal kernel.

The two simulations shown in Fig. 4 and Supplementary Fig. 11 differ in their specifications of $K(x, y)$ and the dispersal kernel. In the uniform local carrying capacity with left-to-right biased migration scenario, $K(x, y) = 200$ everywhere, and the gaussian dispersal kernel has mean $(0.1, 0)$ and standard deviation 0.1. In the left-to-right decreasing local carrying capacity with uniform unbiased migration scenario, the carrying capacity is defined as $K(x, y) = \frac{240\beta e^{-\frac{\beta x}{8}}}{1-e^{-\beta}}$ with $\beta = \log 30$, so that the mean carrying capacity equals 240 and decreases exponentially from left to right, with the carrying capacity at the left boundary being 30 times larger than that at the right boundary. In this case, the gaussian dispersal kernel has mean $(0, 0)$ and standard deviation 0.1.

Each simulation is run for 10,000 generations. We then construct a square lattice network with lattice points located at $(x + 0.5, y + 0.5)$, where $x, y \in \{0, 1, \ldots, 7\}$. For each lattice point, we randomly sample 50 individuals from a smaller square centered at the lattice point with side length 0.75. If fewer than 50 individuals are present within this region, all available individuals are sampled.

During the forward-time simulations, the mutation rate is set to 0. After obtaining the tree sequences for the sampled individuals, we use `msprime` to add mutations. From each tree, we then retain exactly one SNP and then generate the SNP profiles required for FRAME.

#### 2.3 Empirical datasets

The *P. trichocarpa/balsamifera* dataset originates from Geraldes et al. (2013, 2014) and was later adapted by Lundgren and Ralph (2019) for their analysis. In our research, we use this adapted version.

The gray wolf dataset originates from Schweizer et al. (2016). It was initially used in FEEMS (Marcus et al., 2021) and later corrected by Shastry et al. (2025). In our research, we utilize the corrected version.

All ancient human datasets are derived from the 1240k dataset provided by the Allen Ancient DNA Resource (Mallick et al., 2024; Mallick and Reich, 2023), with a focus on autosomal data. Initially, we filter the dataset based on temporal and geographic criteria, including time period, longitude, and latitude.

For the $7000 - 3500$ BCE and $4500 - 3500$ BCE datasets, we include all individuals from the respective periods within the Europe block (longitude $\in [-10°, 35°]$, latitude $\in [32°, 55°]$) and the Eurasia block (longitude $\in [35°, 55°]$, latitude $\in [25°, 55°]$).

For the $3500 - 1500$ BCE and $2500 - 1500$ BCE dataset, we select all individuals from the respective periods within the Europe sample block (lon $\in [-10°, 35°]$, lat $\in [32°, 55°]$) and the Extended Eurasia sample block (lon $\in [35°, 75°]$, lat $\in [25°, 55°]$).

For all four datasets, we then apply the following filters using plink (Purcell et al., 2007):

| Filter (plink name) | Parameters |
|---|---|
| Minor allele frequency (maf) | 0.001 |
| SNP missingness (geno) | 0.4 |
| Individual missingness (mind) | 0.4 |
| Minor allele frequency (maf) | 0.01 |
| Linkage disequilibrium (indep-pairwise) | $50, 5, 0.2$ |

**Table S2:** Filters

The first minor allele frequency filter is applied to remove loci with no variation in the smaller dataset. The SNP missingness filter and the individual missingness filter are used to increase the genotype rate. The second minor allele frequency filter is applied to remove newly occurring mutations that have not been fully spread and thus do not contain sufficient geographical information. The linkage disequilibrium filter is applied to retain SNPs that exhibit nearly perfect segregation. After filtering, all datasets have achieved a genotype rate around 90%.

The final step after the filtering procedures involve removing duplicate samples from the dataset, as some samples were sequenced multiple times using different methods and sequencing depths. References for the samples used in each dataset are provided below. And lists of samples used in each dataset are available at https://github.com/ShenHaotv/frame-analysis.

| Study(Abbreviation in AADR) | Samples | Reference |
|---|---|---|
| AllentoftNature2024 | 17 | (3) |
| AntonioGaoMootsScience2019 | 4 | (5) |
| BraceDiekmannNatureEcologyEvolution2019 | 9 | (11) |
| CassidyNature2020 | 18 | (15) |
| ChildebayevaHaakMolBioEvo2022 | 24 | (17) |
| FeldmanNatureCommunications2019 | 1 | (20) |
| FernandesNatureEcologyEvolution2020 | 7 | (23) |
| FernandesScientificReports2018 | 11 | (22) |
| FowlerOlaldeNature2021 | 25 | (24) |
| FreilichPinhasiScientificReports2021 | 15 | (25) |
| FurtwanglerNatureCommunications2020 | 5 | (26) |
| GambaNatureCommunications2014 | 4 | (27) |
| GelabertBioRxiv2023 | 170 | (29) |
| GelabertSciRep2022 | 9 | (28) |
| GokhmanNatureCommunications2020 | 1 | (32) |
| GonzalesFortesCurrentBiology2017 | 4 | (33) |
| HarneyCheronetGenomeResearch2021 | 4 | (38) |
| HarneyMayNatureCommunications2018 | 13 | (37) |
| HofmanovaPNAS2016 | 5 | (39) |
| JensenSchroederNatureCommunications2019 | 1 | (43) |
| LazaridisAlpaslanRoodenbergScience2022 | 62 | (51) |
| LazaridisNature2014 | 3 | (48) |
| LazaridisNature2016 | 12 | (49) |
| LazaridisNature2017 | 1 | (50) |
| LipsonNature2017 | 42 | (53) |
| MarchiExcoffierCell2022 | 13 | (59) |
| MarcusNatureCommunications2020 | 3 | (61) |
| MarotiTorokCurrBio2022 | 1 | (63) |
| MartinianoPLoSGenetics2017 | 4 | (62) |
| MathiesonNature2015 | 35 | (64) |
| MathiesonNature2018 | 79 | (65) |
| MattilaCommBio2023 | 5 | (66) |
| NarasimhanPattersonScience2019 | 3 | (71) |
| NovakPLoSOne2021 | 29 | (72) |
| OlaldeMBE2015 | 1 | (73) |
| OlaldeNature2018 | 15 | (74) |
| OlaldeScience2019 | 4 | (75) |
| PapacScienceAdvances2021 | 26 | (76) |
| PattersonNature2021 | 15 | (77) |
| PenskeHaakNature2023 | 79 | (78) |
| PosthYuNature2023 | 13 | (81) |
| RivollatNature2023 | 64 | (85) |
| RivollatScienceAdvance2020 | 64 | (84) |
| RohlandMallickGenomeResearch2022 | 3 | (86) |
| SanchezQuintoPNAS2019 | 3 | (98) |
| ScheibAnnHumBio2019 | 1 | (88) |
| SimoesPNAS2024 | 7 | (94) |
| SimoesNature2023 | 5 | (93) |
| SkourtaniotiCell2020 | 16 | (95) |
| VillalbaMoucoCurrentBiology2019 | 4 | (100) |
| WangNatureCommunications2019 | 7 | (102) |
| YuvandeLoosdrechtiScience2022 | 9 | (104) |

**Table S3:** References for samples used in the 7000-3500 BCE dataset

| Study (Abbreviation in AADR) | Samples | Reference |
|---|---|---|
| AgranatTamirWaldmanCell2020 | 17 | (1) |
| AllentoftNature2015 | 7 | (2) |
| AllentoftNature2024 | 3 | (3) |
| AmorimNatureCommunications2018 | 1 | (4) |
| AntonioGaoMootsScience2019 | 3 | (5) |
| ArianoCell2022 | 3 | (6) |
| BlocherPNAS2023 | 32 | (10) |
| BraceDiekmannNatureEcologyEvolution2019 | 4 | (11) |
| BrunelPNAS2020 | 1 | (12) |
| CassidyNature2020 | 19 | (15) |
| CassidyPNAS2016 | 1 | (14) |
| ClementeCell2021 | 6 | (18) |
| DamgaardScience2018 | 9 | (19) |
| FernandesNatureEcologyEvolution2020 | 19 | (23) |
| FernandesScientificReports2018 | 6 | (22) |
| FowlerOlaldeNature2021 | 1 | (24) |
| FreilichPinhasiScientificReports2021 | 7 | (25) |
| FurtwanglerNatureCommunications2020 | 69 | (26) |
| GambaNatureCommunications2014 | 1 | (27) |
| GelabertSciRep2022 | 1 | (28) |
| GonzalesFortesCurrentBiology2017 | 1 | (33) |
| GonzalesFortesProcRoyalSocB2019 | 3 | (34) |
| HaberAJHG2017 | 4 | (35) |
| HarneyCheronetGenomeResearch2021 | 1 | (38) |
| ImmelCommBiol2021 | 33 | (41) |
| ImmelNature2020 | 2 | (40) |
| IngmanStockhammerPLoS2021 | 8 | (42) |
| JeongNatureEcologyEvolution2019 | 1 | (44) |
| KoptekinCurrentBiology2023 | 7 | (45) |
| KrzewinskaScienceAdvances2018 | 6 | (46) |
| LazaridisAlpaslanRoodenbergScience2022 | 90 | (51) |
| LazaridisNature2016 | 6 | (49) |
| LazaridisNature2017 | 4 | (50) |
| LinderholmNatureScientificReports2020 | 4 | (52) |
| LipsonNature2017 | 16 | (53) |
| MaierFlegontovELife2023 | 1 | (55) |
| MalmstromProcBiolSci2019 | 1 | (58) |
| MarcusNatureCommunications2020 | 21 | (61) |
| MartinianoPLoSGenetics2017 | 6 | (62) |
| MathiesonNature2015 | 37 | (64) |
| MathiesonNature2018 | 23 | (65) |
| MittnikScience2019 | 20 | (69) |
| MootsNatEcolEvol2023 | 2 | (70) |
| NarasimhanPattersonScience2019 | 130 | (71) |
| OlaldeNature2018 | 157 | (74) |
| OlaldeScience2019 | 59 | (75) |
| PapacScienceAdvances2021 | 140 | (76) |
| PattersonNature2021 | 74 | (77) |
| PenskeHaakNature2023 | 12 | (78) |
| PenskeSciRep2024 | 40 | (79) |
| PosthYuNature2023 | 2 | (81) |
| RivollatScienceAdvance2020 | 1 | (84) |
| RohlandMallickGenomeResearch2022 | 4 | (86) |
| SaupeScheibCurrBio2021 | 1 | (87) |
| ScheibAnnHumBio2019 | 1 | (88) |
| SchroederPNAS2019 | 17 | (89) |
| SeguinOrlandoCurrBio2021 | 9 | (91) |
| SkourtaniotiCell2020 | 41 | (95) |
| SkourtaniotiRingbauerNatureEcologyEvolution2023 | 28 | (96) |
| ValdioseraPNAS2018 | 3 | (99) |
| VillalbaMoucoSciAdv2021 | 88 | (101) |
| WangKrauseCellGenomics2023 | 1 | (103) |
| WangNatureCommunications2019 | 28 | (102) |
| | | *Continued on next page* |

| Study (Abbreviation in AADR) | Samples | Reference |
|---|---|---|
| ZegaracScientificReports2021 | 18 | (105) |

**Table S4:** References for samples used in the 3500-1500 BCE dataset

| Study (Abbreviation in AADR) | Samples | Reference |
|---|---|---|
| AllentoftNature2024 | 3 | (3) |
| BraceDiekmannNatureEcologyEvolution2019 | 15 | (11) |
| CassidyNature2020 | 17 | (15) |
| FernandesNatureEcologyEvolution2020 | 3 | (23) |
| FernandesScientificReports2018 | 10 | (22) |
| FowlerOlaldeNature2021 | 27 | (24) |
| FreilichPinhasiScientificReports2021 | 13 | (25) |
| FurtwanglerNatureCommunications2020 | 2 | (26) |
| GambaNatureCommunications2014 | 1 | (27) |
| GelabertBioRxiv2023 | 1 | (29) |
| GelabertSciRep2022 | 9 | (28) |
| HarneyCheronetGenomeResearch2021 | 2 | (38) |
| HarneyMayNatureCommunications2018 | 11 | (37) |
| HofmanovaPNAS2016 | 2 | (39) |
| JensenSchroederNatureCommunications2019 | 1 | (43) |
| LazaridisAlpaslanRoodenbergScience2022 | 45 | (51) |
| LazaridisNature2016 | 8 | (49) |
| LipsonNature2017 | 12 | (53) |
| MarcusNatureCommunications2020 | 3 | (61) |
| MartinianoPLoSGenetics2017 | 4 | (62) |
| MathiesonNature2018 | 15 | (65) |
| MattilaCommBio2023 | 5 | (66) |
| NarasimhanPattersonScience2019 | 1 | (71) |
| NovakPLoSOne2021 | 27 | (72) |
| OlaldeNature2018 | 16 | (74) |
| OlaldeScience2019 | 2 | (75) |
| PapacScienceAdvances2021 | 26 | (76) |
| PattersonNature2021 | 9 | (77) |
| PenskeHaakNature2023 | 46 | (78) |
| RivollatNature2023 | 3 | (85) |
| RivollatScienceAdvance2020 | 24 | (84) |
| SanchezQuintoPNAS2019 | 2 | (98) |
| ScheibAnnHumBio2019 | 1 | (88) |
| SimoesNature2023 | 1 | (93) |
| SkourtaniotiCell2020 | 10 | (95) |
| VillalbaMoucoCurrentBiology2019 | 2 | (100) |
| WangNatureCommunications2019 | 4 | (102) |

**Table S5:** References for samples used in the 4500-3500 BCE dataset

| Study (Abbreviation in AADR) | Samples | Reference |
|---|---|---|
| AgranatTamirWaldmanCell2020 | 17 | (1) |
| AllentoftNature2015 | 5 | (2) |
| AmorimNatureCommunications2018 | 1 | (4) |
| AntonioGaoMootsScience2019 | 1 | (5) |
| ArianoCell2022 | 2 | (6) |
| BlocherPNAS2023 | 32 | (10) |
| ClementeCell2021 | 6 | (18) |
| DamgaardScience2018 | 3 | (19) |
| FernandesNatureEcologyEvolution2020 | 19 | (23) |
| FernandesScientificReports2018 | 5 | (22) |
| FreilichPinhasiScientificReports2021 | 6 | (25) |
| FurtwanglerNatureCommunications2020 | 26 | (26) |
| GelabertSciRep2022 | 1 | (28) |
| GonzalesFortesProcRoyalSocB2019 | 1 | (34) |
| HaberAJHG2017 | 4 | (35) |
| HarneyCheronetGenomeResearch2021 | 1 | (38) |
| IngmanStockhammerPLoS2021 | 8 | (42) |
| KrzewinskaScienceAdvances2018 | 6 | (46) |
| LazaridisAlpaslanRoodenbergScience2022 | 63 | (51) |
| LazaridisNature2016 | 3 | (49) |
| LazaridisNature2017 | 4 | (50) |
| LinderholmNatureScientificReports2020 | 4 | (52) |
| MarcusNatureCommunications2020 | 19 | (61) |
| MartinianoPLoSGenetics2017 | 2 | (62) |
| MathiesonNature2015 | 20 | (64) |
| MathiesonNature2018 | 5 | (65) |
| MittnikScience2019 | 44 | (69) |
| MootsNatEcolEvol2023 | 2 | (70) |
| NarasimhanPattersonScience2019 | 92 | (71) |
| OlaldeNature2018 | 148 | (74) |
| OlaldeScience2019 | 24 | (75) |
| PapacScienceAdvances2021 | 95 | (76) |
| PattersonNature2021 | 61 | (77) |
| PenskeSciRep2024 | 40 | (79) |
| RohlandMallickGenomeResearch2022 | 1 | (86) |
| SeguinOrlandoCurrBio2021 | 3 | (91) |
| SkourtaniotiCell2020 | 20 | (95) |
| SkourtaniotiRingbauerNatureEcologyEvolution2023 | 27 | (96) |
| VillalbaMoucoSciAdv2021 | 70 | (101) |
| WangNatureCommunications2019 | 7 | (102) |
| ZegaracScientificReports2021 | 18 | (105) |

**Table S6:** References for samples used in the 2500-1500 BCE dataset

**Figure S1: Runtimes for FRAME method vs the direct solution of the linear system of Equation S2**

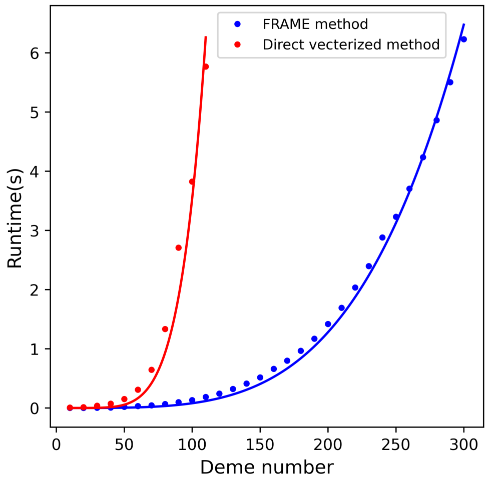

Average runtimes for solving equation (S2) using FRAME and brute-force methods are shown, for a given number of demes ($x$-axis) connected via random migration rates in a fully connected graph and with random coalescence rates (see Methods).

**Figure S2: FRAME fit to equilibrium migration simulations with different patterns and sampling schemes (Difference graph)**

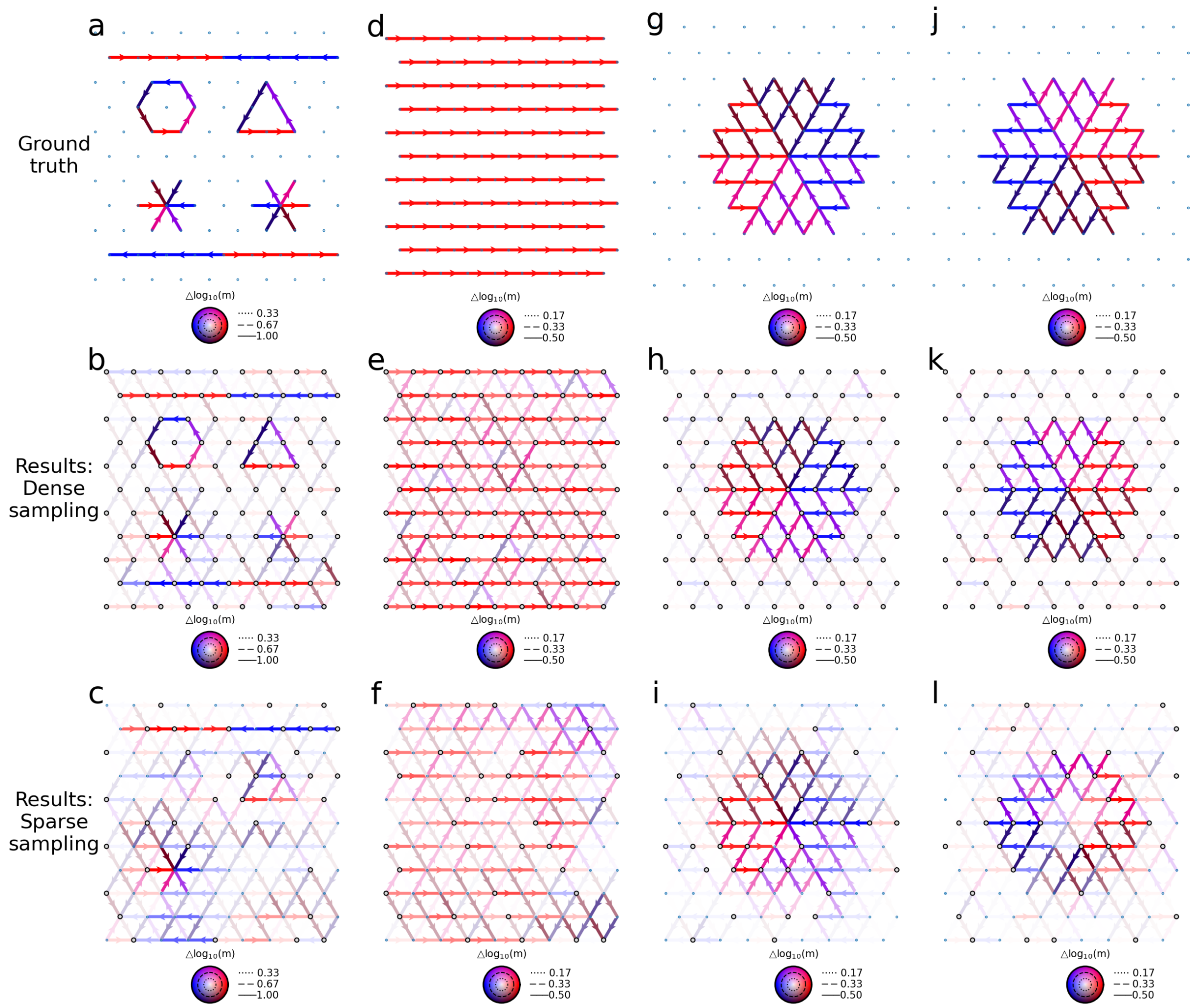

(**a–l**) Difference graphs corresponding to full graphs in Figure 3.

**Figure S3: FRAME fit to equilibrium migration simulations with different patterns and sampling schemes (Goodness of fit)**

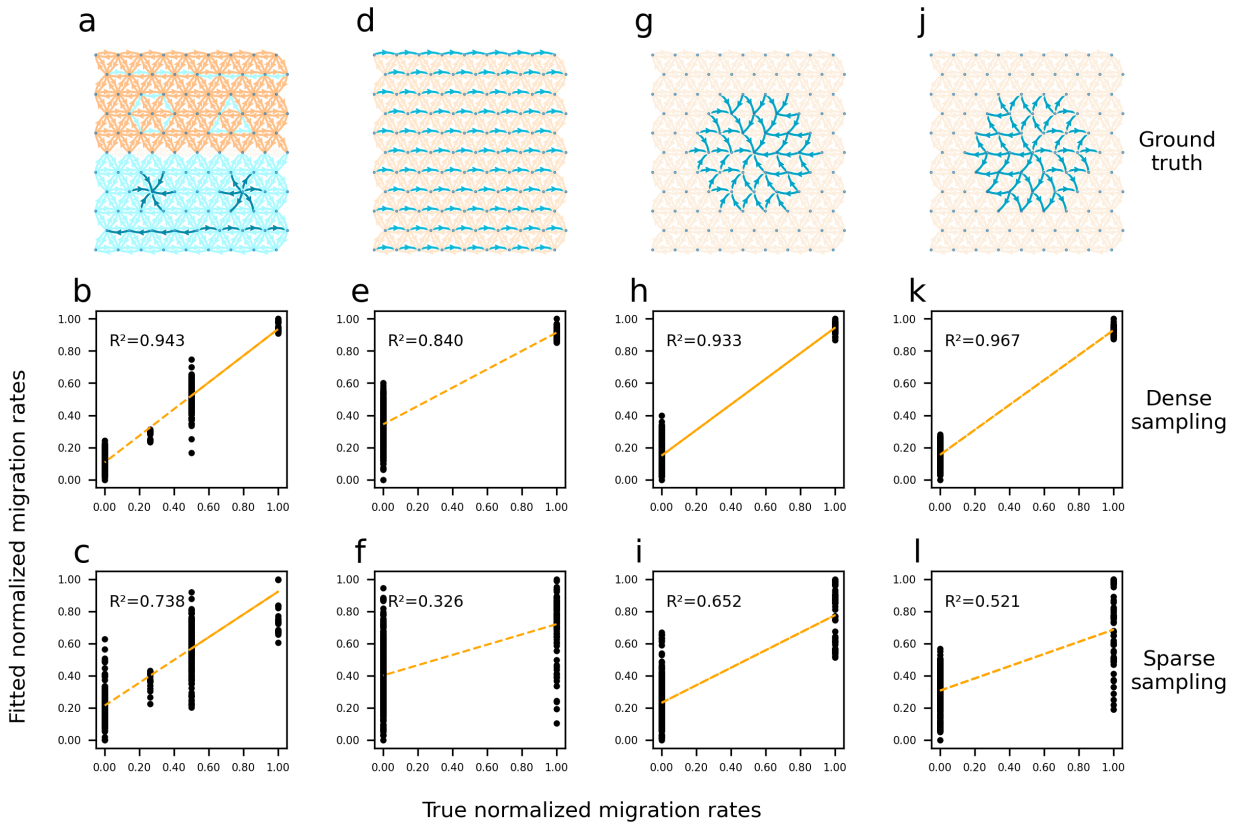

(**a–l**) Ground truth and comparison between fitted and true migration rates for the full graphs shown in Fig.3. Migration rates are shown on the log scale and normalized to the interval $[0, 1]$. The coefficient of determination ($R^2$) quantifies the goodness of fit.

**Figure S4: FRAME fit to simulations of large scale directional flow with equal coalescence rates**

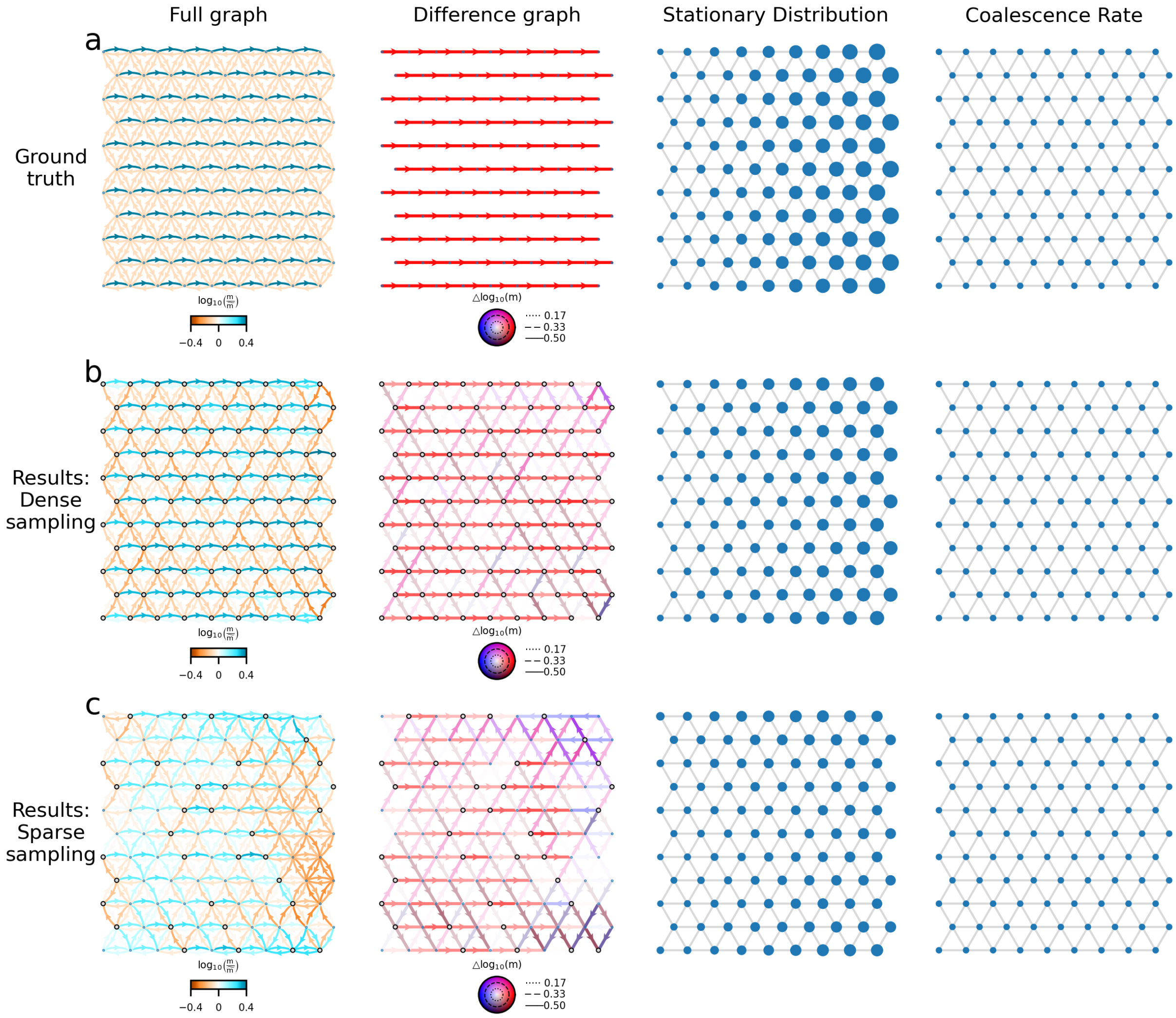

**(a-c)** Ground truth and FRAME fit to simulations of large scale directional flow with equal coalescence rates, varying the sampling design

**Figure S5: FRAME fit to simulations of large scale directional flow with coalescence rates inversely proportional to stationary distribution**

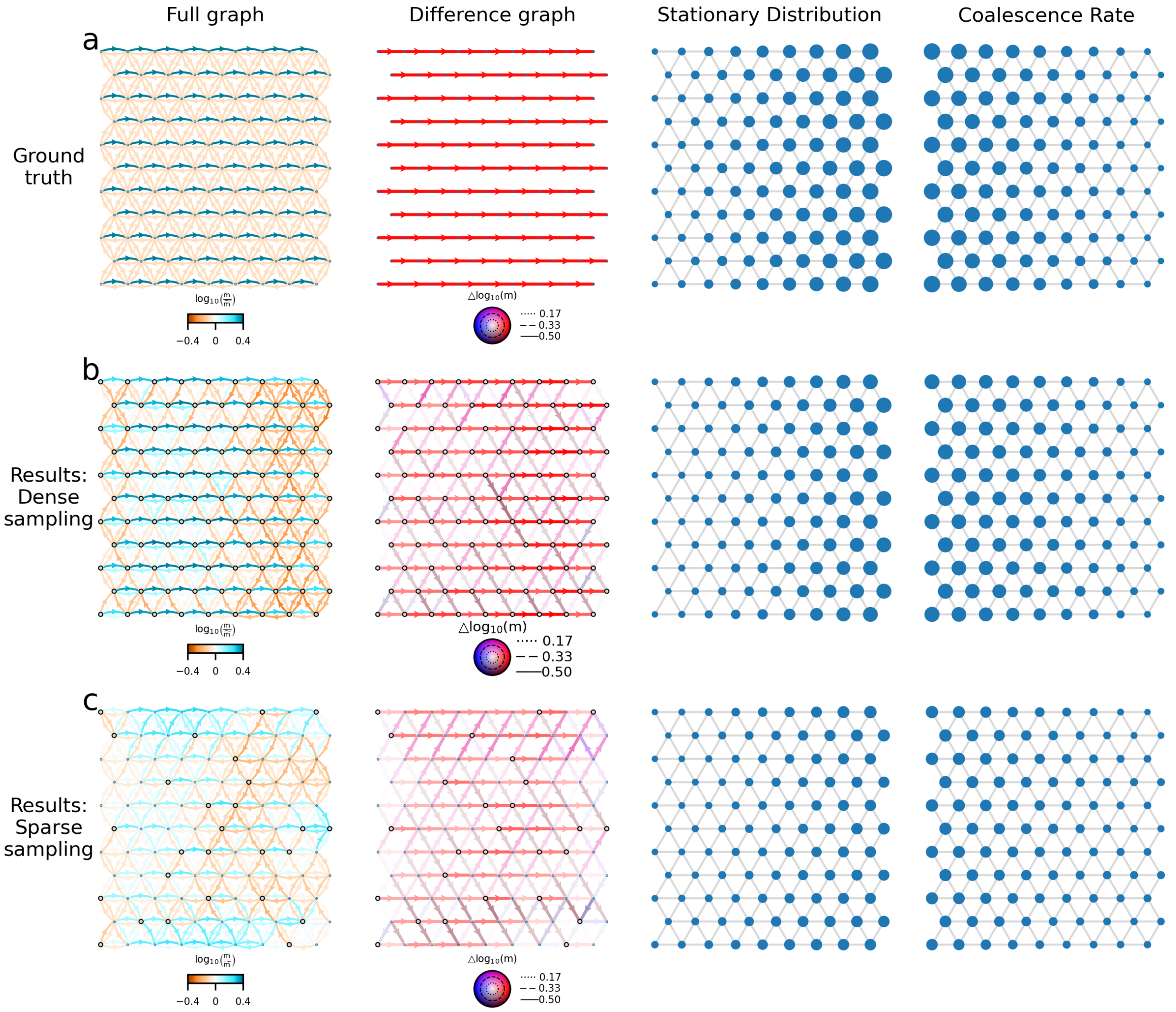

**(a-c)** Ground truth and FRAME fit to simulations of large scale directional flow with coalescence rates inversely proportional to stationary distribution, varying the sampling design

**Figure S6: Comparison of FRAME with Lundgren and Ralph (2019) on the *P. trichocarpa/balsamifera* dataset**

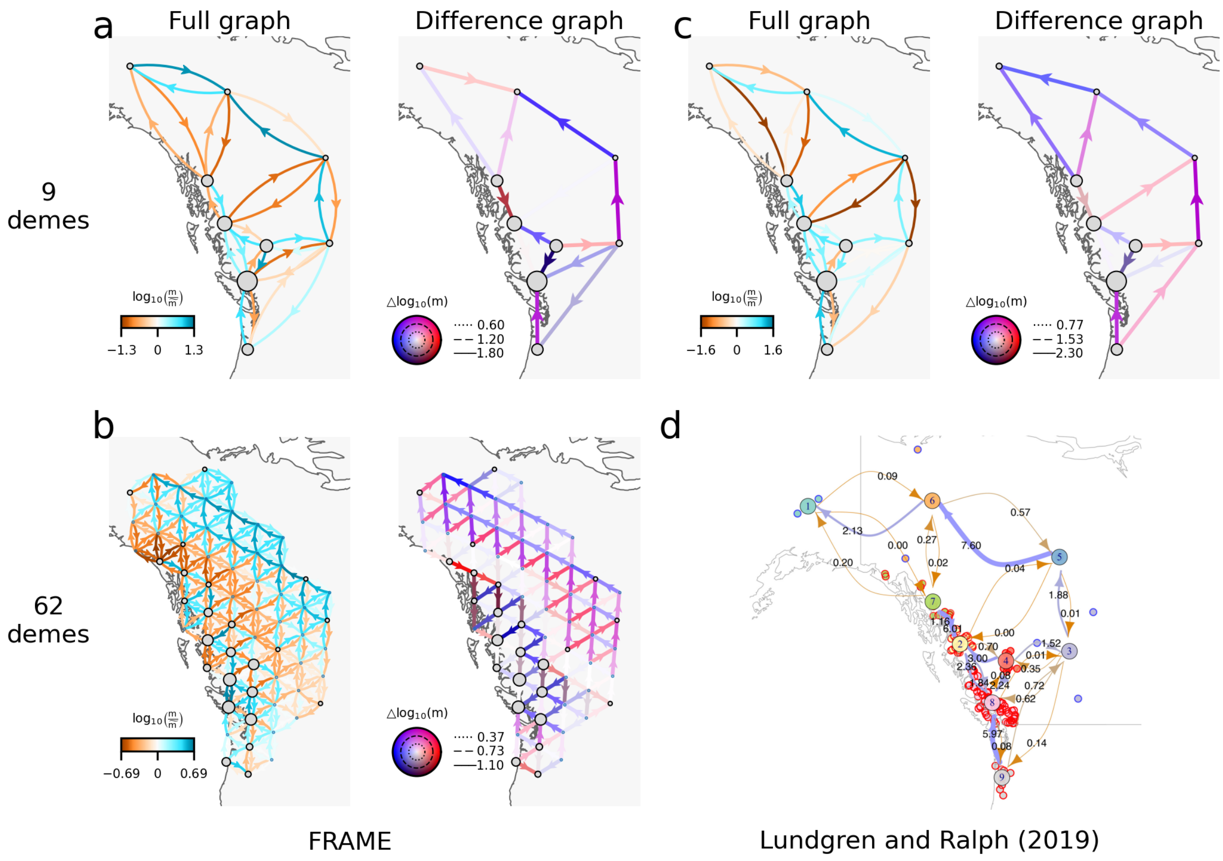

**(a–b)** FRAME analysis of the *P. trichocarpa/balsamifera* datasets dataset at different resolutions. **(c)** Lundgren and Ralph (2019) results visualized in FRAME's representation, with zero edges replaced by a minimal migration rate of 0.01. **(d)** Lundgren and Ralph (2019) results in their original representation.

**Figure S7: Comparison of FRAME with FEEMS on the North American wolf dataset**

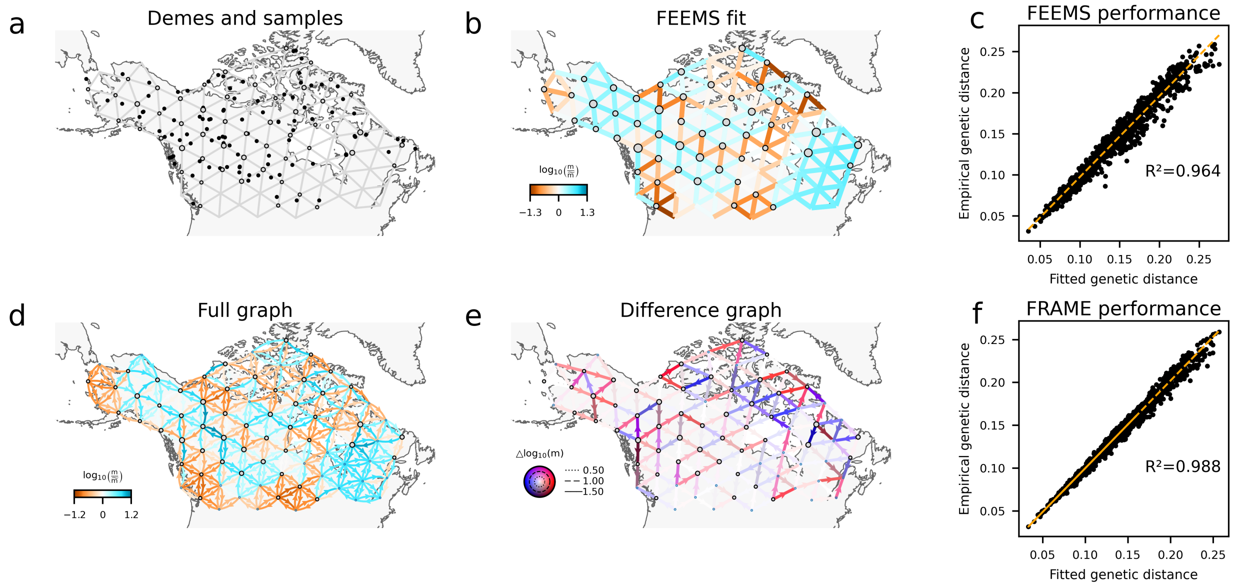

**(a)** Demes and samples. **(b)** FEEMS fit of the dataset. **(c)** FEEMS performance. **(d-e)** FRAME fit of the dataset. **(f)** FRAME performance.

**Figure S8: Comparison of the 7000-3500 BCE dataset with the 4500-3500 BCE dataset**

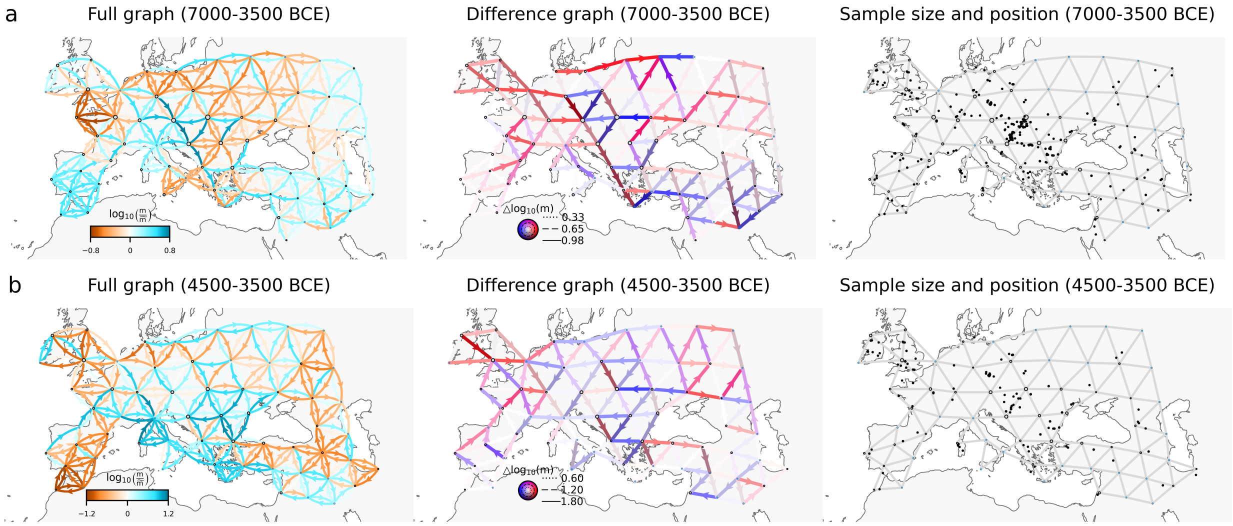

**(a)** FRAME analysis of the 7000-3500 BCE dataset. **(b)**FRAME analysis of the 4500-3500 dataset. Although FRAME analysis of this dataset shares some asymmetric signals with the FRAME analysis of the 7000-3500 BCE dataset, it detects fewer patterns consistent with our current knowledge Neolithic expansion. This discrepancy is likely due to the smaller sample size (383 individuals versus 975 in the broader window) and time-varying gene flow dynamics during these periods. For instance, the 4500-3500 BCE dataset lacks samples from southern France, and we observe different signals along the northern Mediterranean coast. Similarly, the significant reduction of number of samples from Germany probably leads to notable differences in the inferred migration patterns in that region compared to the 7000-3500 BCE dataset.

**Figure S9: Comparison of the 3500-1500 BCE dataset with the 2500-1500 BCE dataset**

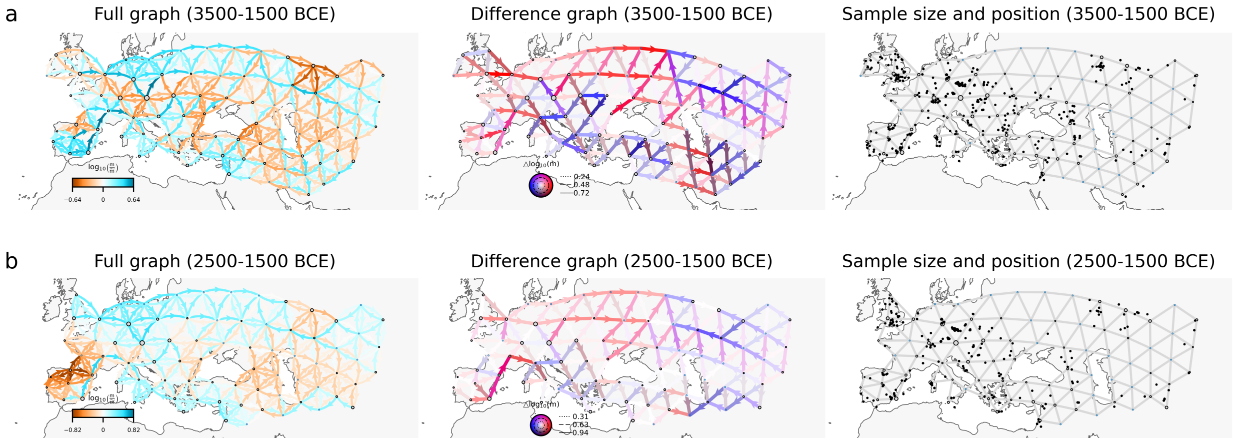

**(a)** FRAME analysis of the 3500-1500 BCE dataset. **(b)** FRAME analysis of the 2500-1500 dataset. Compared with Figure S5, FRAME analysis of the 2500-1500 BCE dataset shows more similarity with FRAME analysis of the $3500 - 1500$ BCE dataset. The asymmetries plausibly reflecting gene flow out of the Steppe region is a shared signal in both analyses. This similarity is likely due to relative large sample sizes of the $2500 - 1500$ BCE dataset (916 individuals versus 1361 individuals in the $3500 - 1500$ BCE dataset).

**Figure S10: Comparison between automatically generated and refined networks**

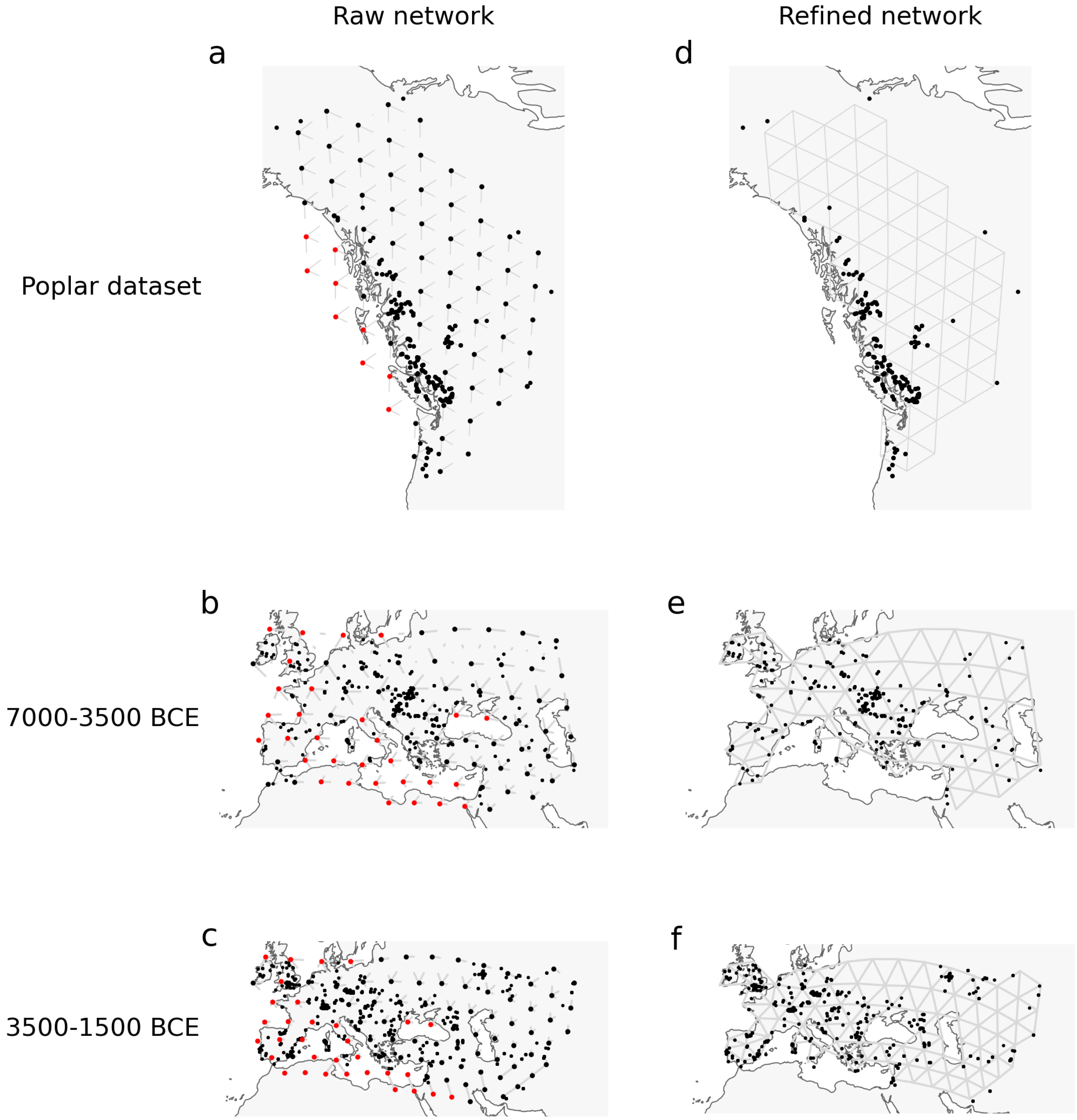

**(a–c)** Automatically generated networks for the *P. trichocarpa/balsamifera* dataset, the 7000–3500 BCE dataset, and the 3500–1500 BCE dataset. Nodes highlighted in red denote those that are modified during network refinement. **(d-f)** Refined networks. Refinements are applied to remove grid cells located in oceans or lakes and to adjust grid placement so that nodes are closer to the sampled individuals.

**Figure S11: Distinguishing forward-in-time biased migration from carrying-capacity gradients**

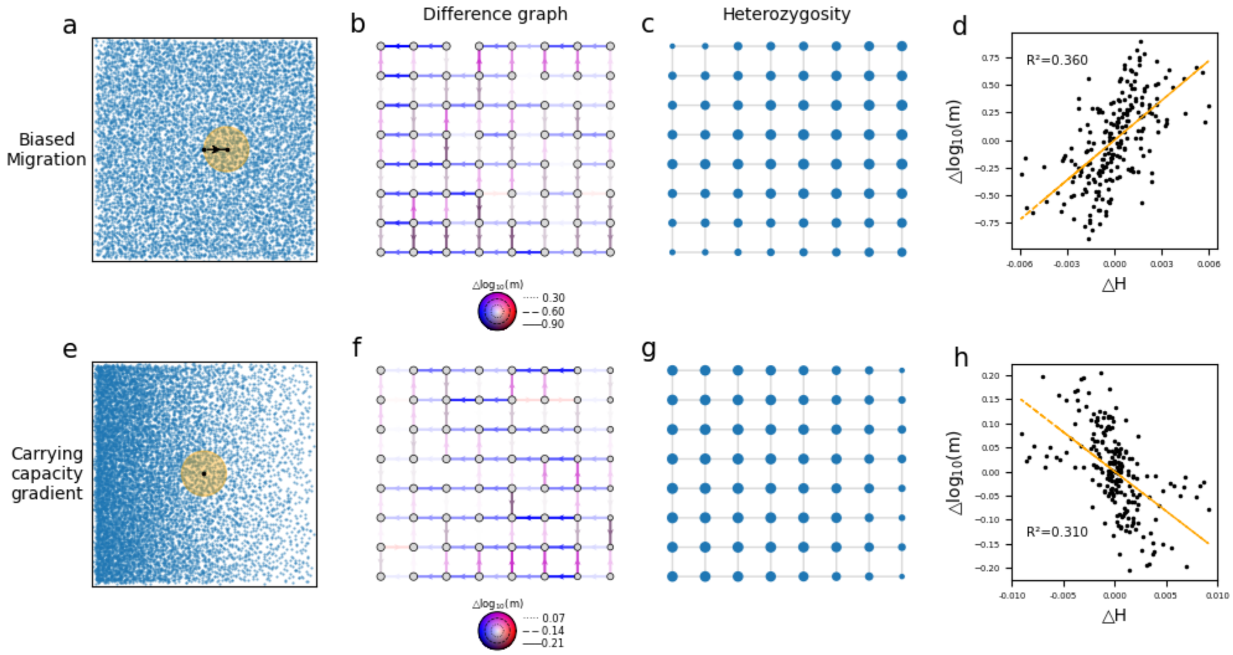

**(a–d)** Under uniform carrying capacity with uniform left-to-right biased migration, heterozygosity increases from left to right. The backward in time migration rate difference (on a log scale) is positively correlated with the heterozygosity difference. **(e–h)** Under a left-to-right decreasing carrying-capacity gradient with uniform, unbiased migration, heterozygosity decreases from left to right. The backward intime migration rate difference (on a log scale) is negatively correlated with the heterozygosity difference.

**Figure S12: Influence of divergence time on FRAME inference**

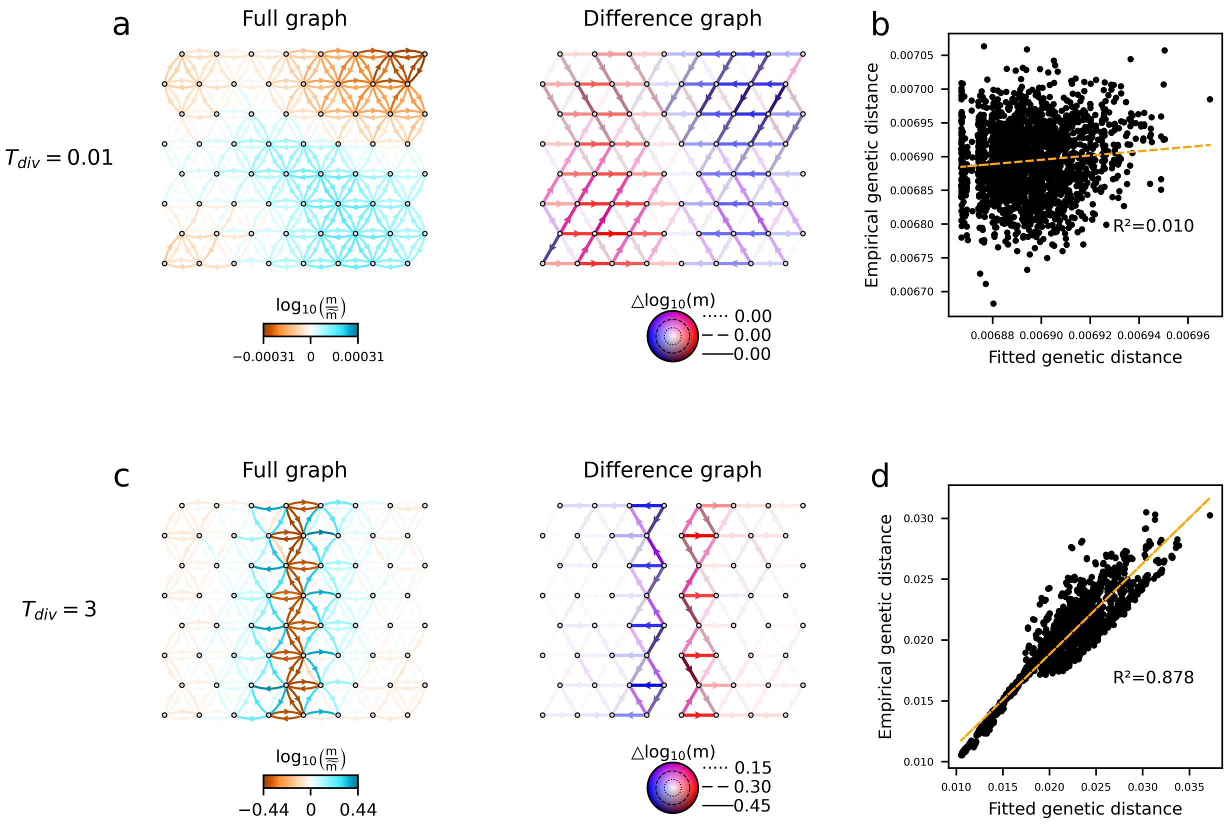

**(a–d)** FRAME fit to a population split-and-divergence model. A panmictic population splits into two subpopulations separated by the central row and diverges for a duration $T_{\text{div}}$. Within each subpopulation, local migration rates between neighbouring demes ($m$) and local effective population sizes ($N$) satisfy $Nm = 1$. **(a–b)** FRAME fit when $T_{div} = 0.01$. Because the divergence time is very short, FRAME detects little structure. Genetic distances are poorly fitted, as the coalescence process is dominated by shared drift in the ancestral panmictic population, and differentiation induced by the migration pattern is weak. **(c–d)** FRAME fit when $T_{div} = 3$. When divergence time is sufficiently long, FRAME clearly detects reduced connectivity due to isolation. Genetic distances are fitted much better, as the migration pattern has persisted long enough to leave a detectable signal.

**Figure S13: Graph Representations**

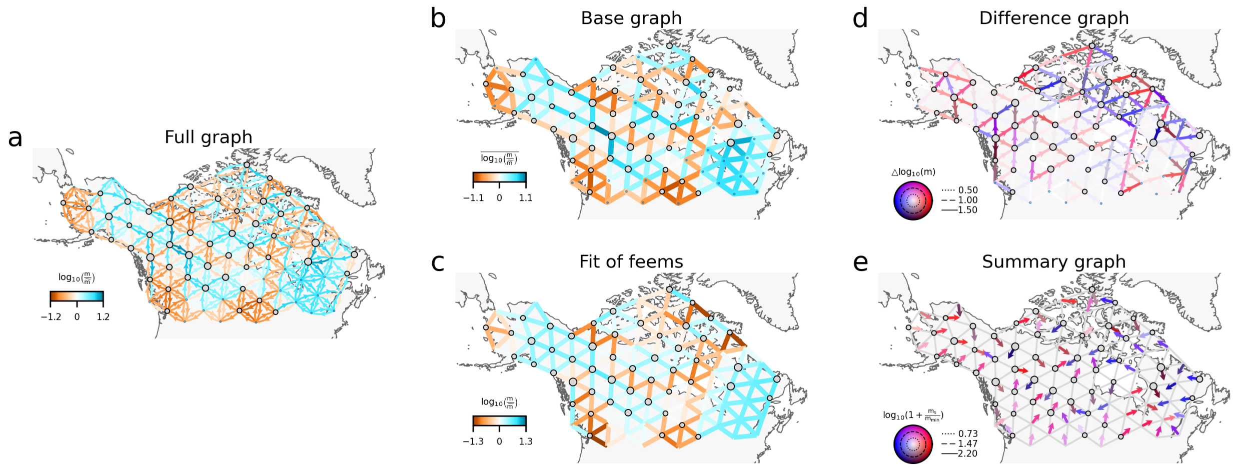

**(a)** Full graph representation uses inferred migration rates as edge weights in a directed graph, depicting the complete migration scenario. **(b)** For a pair of connected demes (A, B), the base graph representation assigns the average log-scaled migration rate $\frac{1}{2}(\log m_{AB} + \log m_{BA})$ to the undirected edge AB. Qualitatively, it resembles FEEMS and EEMS and can be interpreted as a connectivity graph. **(c)** FEEMS fit, shown for comparison with the base graph. **(d)** For a pair of connected demes (A, B) where $m_{AB} > m_{BA}$, the difference graph representation assigns the log-scale difference $\log m_{AB} - \log m_{BA}$ to the directed edge AB. This representation complements the base graph. **(e)** Summary graph representation sums all outgoing edge vectors at each node, producing a migration vector field.

**Figure S14: FRAME fit to the *P. trichocarpa/balsamifera* dataset (full version)**

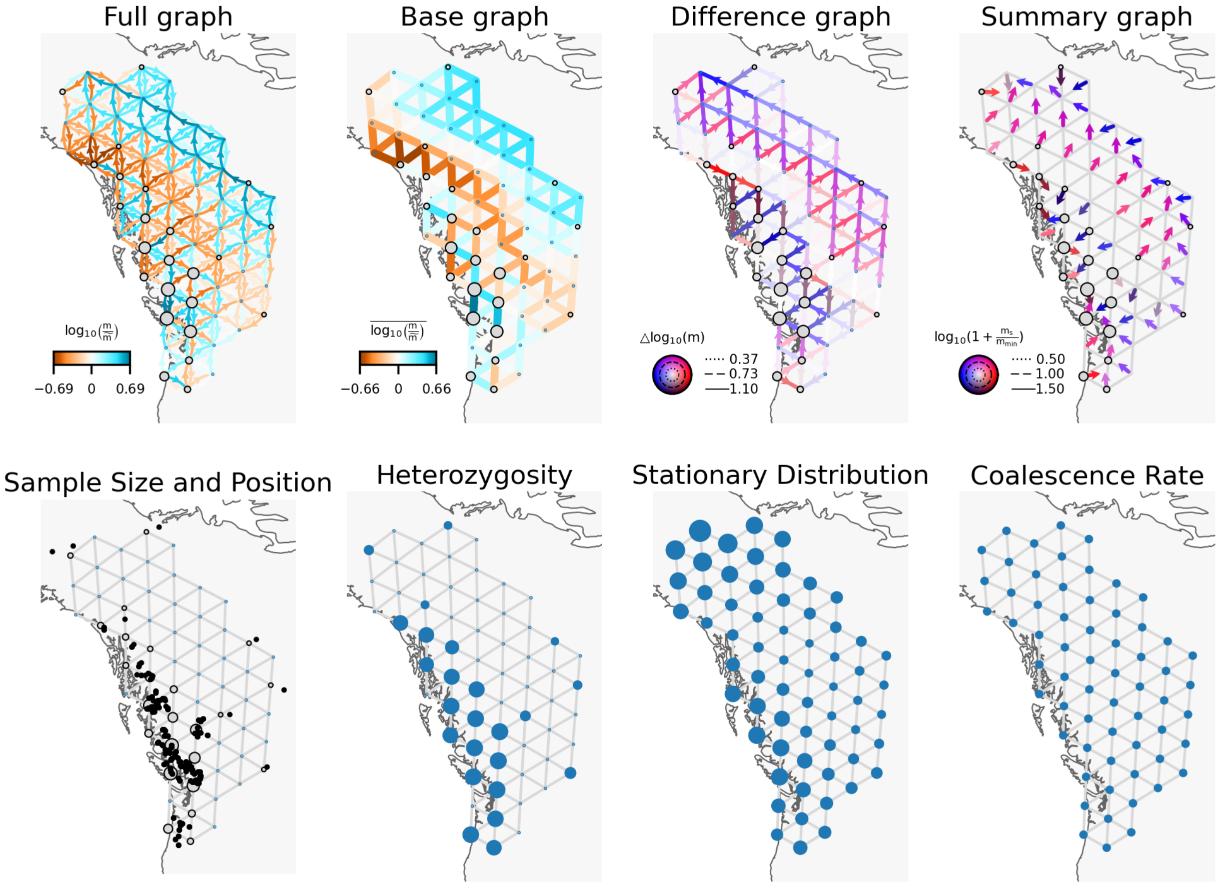

The first row presents all four representations of the FRAME fit to the *P. trichocarpa/balsamifera* dataset. The second row displays the corresponding dataset attributes.

**Figure S15: FRAME fit to the wolf dataset (full version)**

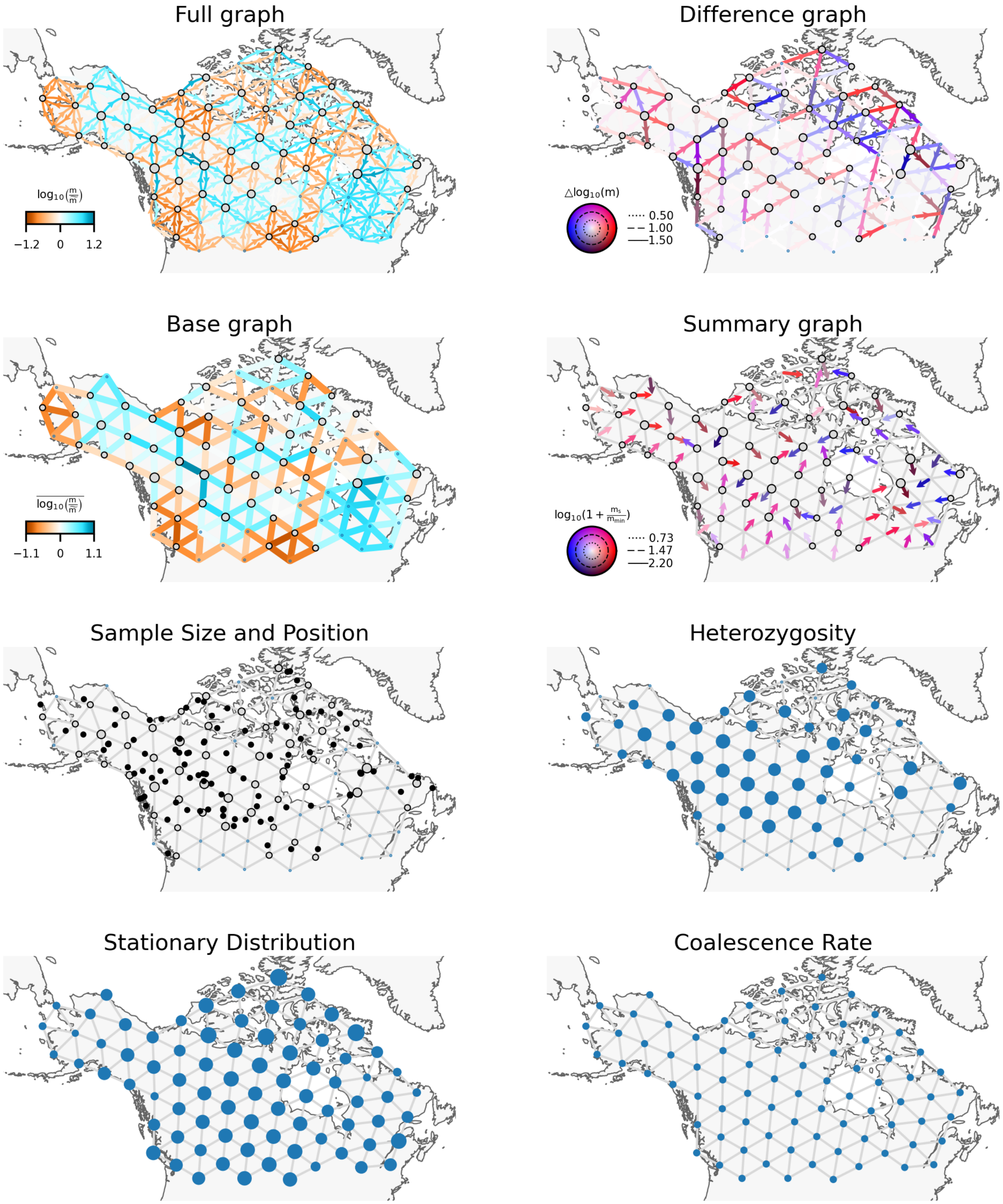

The first two rows presents all four representations of the FRAME fit to the wolf dataset. The third and fourth row displays the corresponding dataset attributes.

**Figure S16: FRAME fit to the 7000-3500 BCE aDNA dataset (full version)**

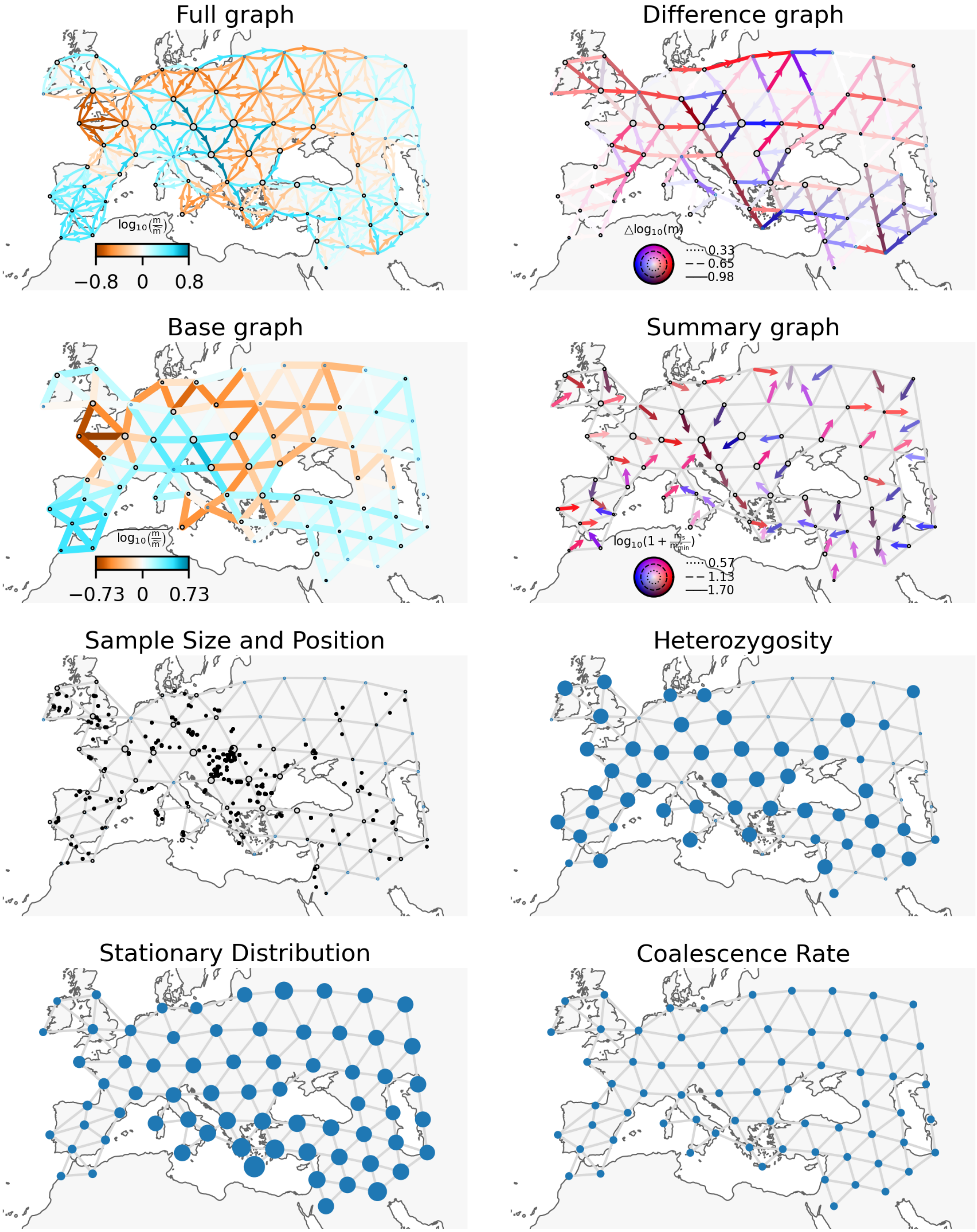

The first two rows presents all four representations of the FRAME fit to the 7000-3500 BCE aDNA dataset. The third and fourth row displays the corresponding dataset attributes.

**Figure S17: FRAME fit to the 3500-1500 BCE aDNA dataset (full version)**

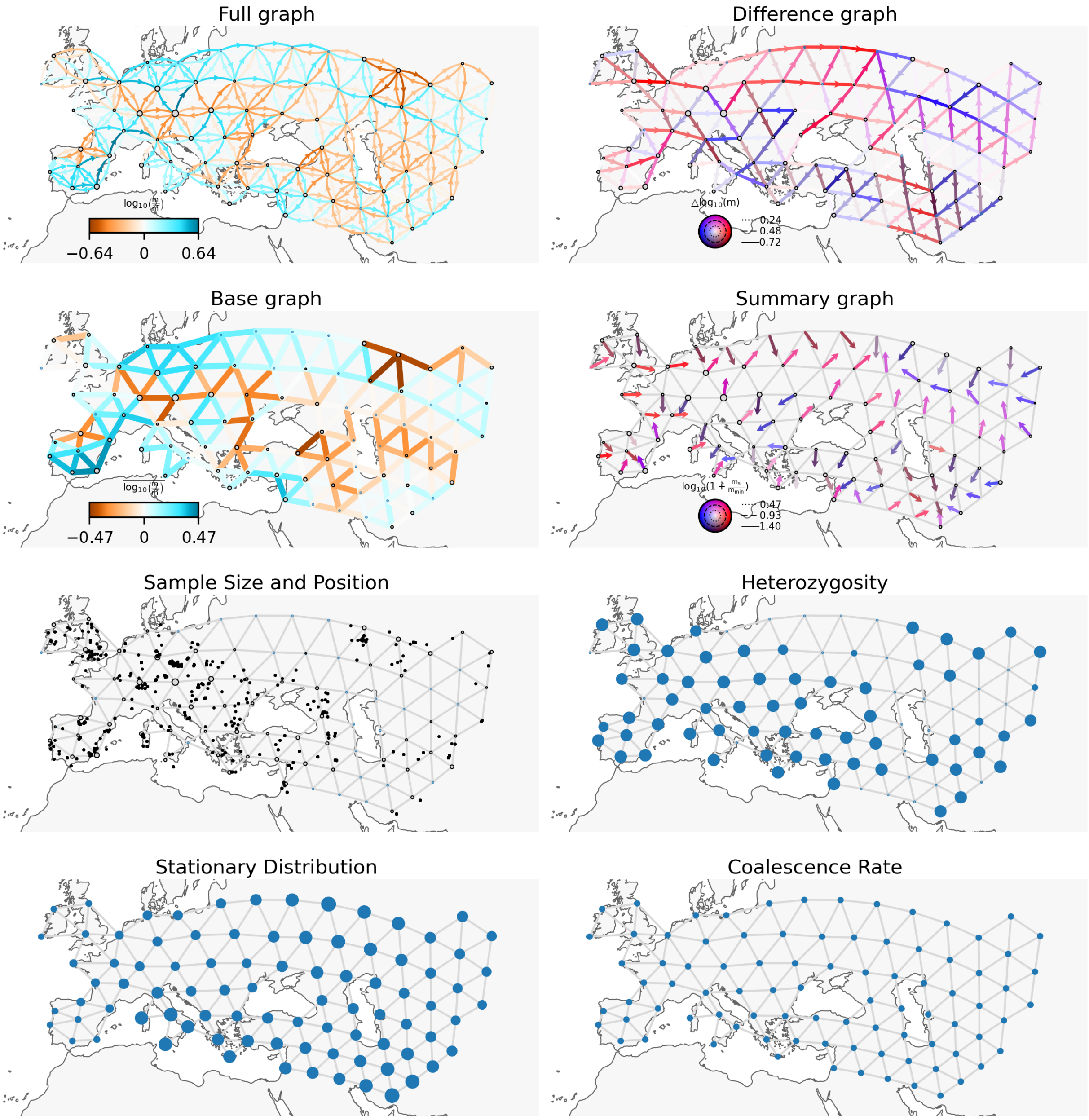

The first two rows presents all four representations of the FRAME fit to the 3500-1500 BCE aDNA dataset. The third and fourth row displays the corresponding dataset attributes.

**Figure S18: FRAME fit to the 4500-3500 BCE aDNA dataset (full version)**

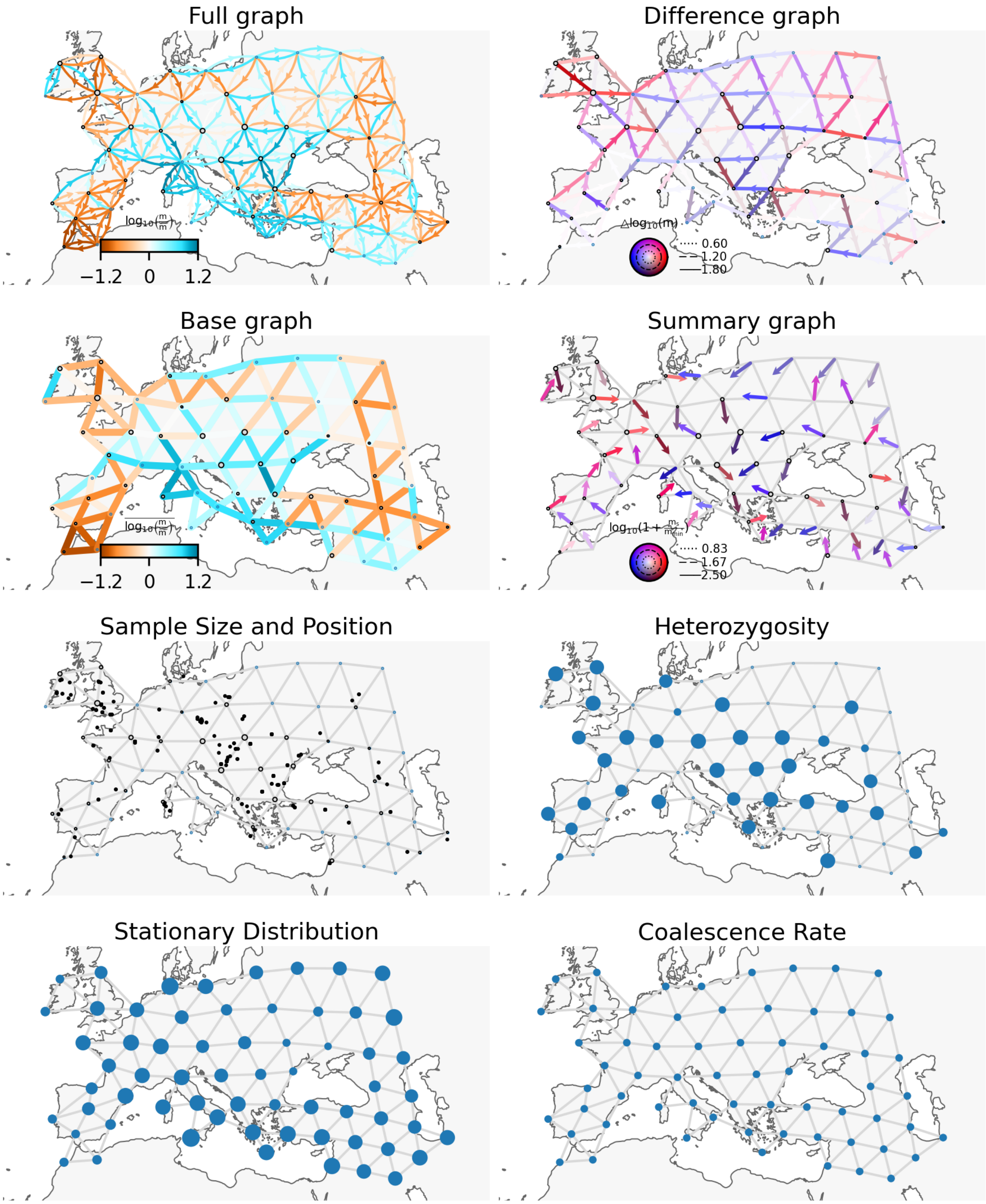

The first two rows presents all four representations of the FRAME fit to the 4500-3500 BCE aDNA dataset. The third and fourth row displays the corresponding dataset attributes.

**Figure S19: FRAME fit to the 2500-1500 BCE aDNA dataset (full version)**

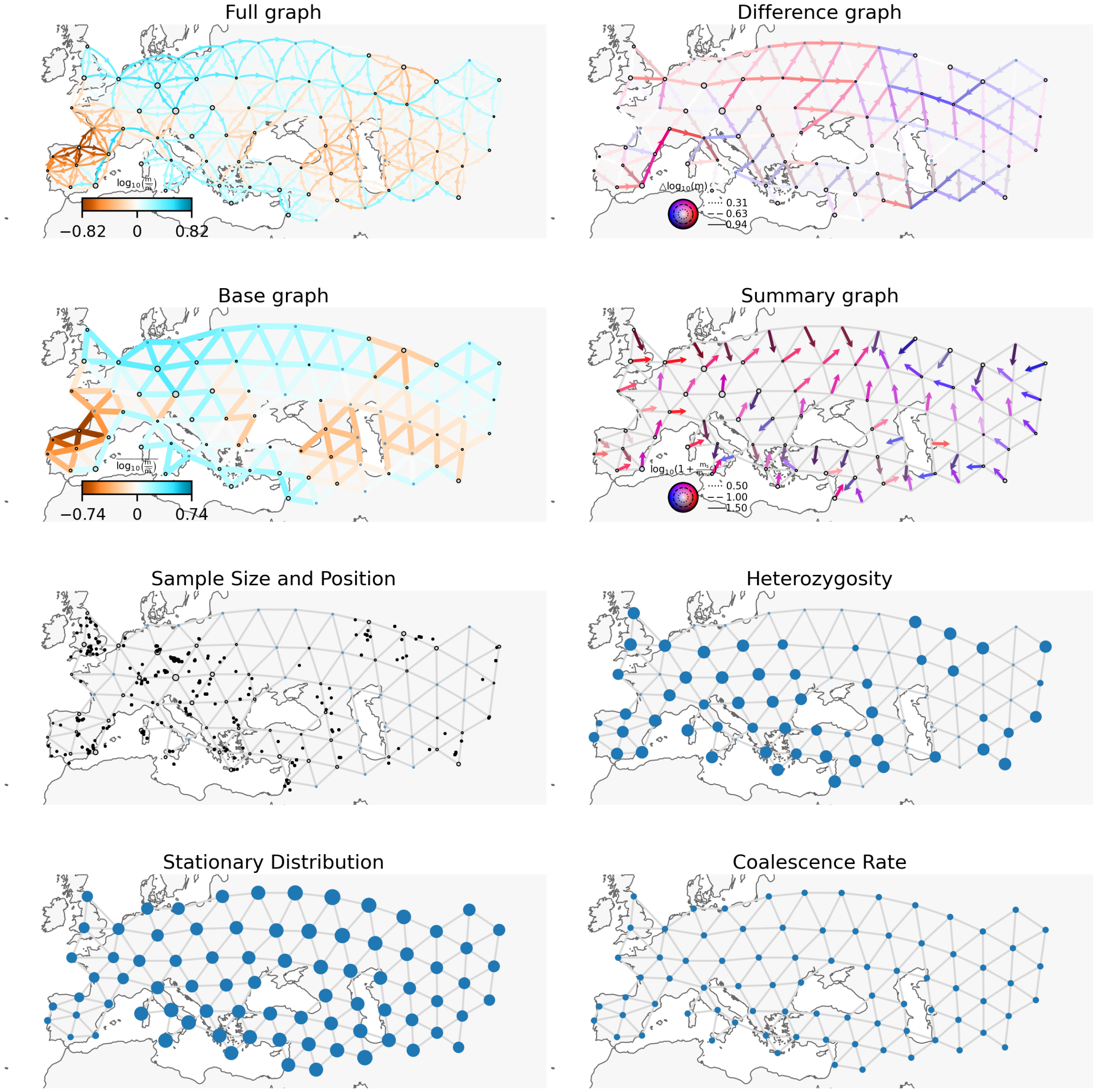

The first two rows presents all four representations of the FRAME fit to the 2500-1500 BCE aDNA dataset. The third and fourth row displays the corresponding dataset attributes.
